# Forecasting oncogene amplification and tumour suppressor deletion

**DOI:** 10.1101/2025.06.06.658212

**Authors:** Barbara Hernando, Angel Fernandez-Sanroman, Alice Cadiz, Patricia G. Santamaria, David Gómez-Sánchez, Maria Escobar-Rey, Blas Chaves-Urbano, Joseph Thompson, Marina Torres, Gorka Ruiz de Garibay, Vera Adradas, Eva Álvarez, The Pan Prostate Cancer Group, Maxime Tarabichi, Tom Lesluyes, Juan Manuel Coya, Jon Zugazagoitia, Luis Paz-Ares, Geoff Macintyre

**Author notes:** These authors contributed equally to this work (Barbara Hernando, Angel Fernandez-Sanroman).

## Abstract

Oncogene amplification and tumour suppressor deletion can drive tumour initiation, progression and treatment resistance. Detection at diagnosis often signals poor prognosis, but it can also enable opportunities for treatment with highly effective targeted therapies. Predicting the likelihood that a patient will acquire these driver alterations in the future using a genomic test represents an opportunity to realise the benefits of interventions earlier, potentially with preventative intent. Here, we present a forecasting framework that takes as input a DNA copy number profile and predicts whether the tumour will acquire an oncogene amplification or tumour suppressor deletion in the future. This framework leverages mutation rate estimates from the input tumour, alongside gene-specific selection coefficients derived from a large cohort of 7,880 tumours. We demonstrate feasibility using 7,042 single-time-point samples and longitudinally collected tumour pairs from 44 prostate and 100 lung cancers, identifying tumours that went on to acquire amplifications at a later time point with an average AUC of 0.87. We show potential clinical utility by forecasting poor prognosis in low-grade gliomas via *CDK4/PDGFRA* amplification or *CDKN2A* deletion, and osimertinib resistance in lung cancers via *MET* amplification. This study serves as a proof-of-concept for a new class of biomarker, wherein selective pressures and mutation-generating processes can be harnessed to anticipate future genomic alterations.

## Introduction

Tumours are highly dynamic entities and, as such, their fates remain difficult to predict. Despite their genomic complexity, analysis of large genome sequencing data across different cancer types indicates that tumour evolution is far from stochastic^1–4^. Instead, frequent recurrence of key driver alterations is observed, alongside convergence to a core set of evolutionary trajectories^3–6^. Recently, pronounced determinism in tumour evolution has been confirmed *in vivo* using bottom-up models of tumour initiation and progression^1,2^. The deterministic nature of tumour evolution lays a foundation for the construction of predictive models where the next genomic alteration and subsequent step in cancer progression can be anticipated. From a clinical perspective, an ability to forecast tumour evolution might allow improved patient risk stratification by predicting the acquisition of somatic alterations foreboding poor outcomes^7–9^, and improved treatment selection by forecasting the acquisition of targetable alterations^10^ or those that confer drug resistance^11^.

The key principles known to govern tumour genome evolution can inform the construction of such predictive models. Namely, the rate at which a genome alteration can spontaneously arise within a tumour cell, and how much that given alteration increases the cell’s fitness. The latter typically depends on complex epistatic and environmental interactions, while the former depends on a complex interplay between global mutation rate, active mutational processes and their biases across the genome due to, among other factors, the overlying DNA topology and epigenetic status. This complexity challenges full forecasting of tumour evolution. However, this complexity is greatly reduced for focal oncogene amplifications and homozygous deletions^12–14^, where only a smaller set of mutational processes can generate the required architectural changes within the genome, and phenotypic effects are usually strongly advantageous. Leveraging this opportunity, we present a predictive framework that models the determinants underpinning the fixation of oncogene amplification and homozygous deletion. Our goal is to perform these predictions in clinically relevant contexts, thus we designed our framework to make predictions using only tumour DNA information derived from a single genomic test that is compatible with current diagnostic assays.

## Results

### A framework for forecasting driver amplifications and deletions

Tumour evolution has long been conceptualised through the lens of population genetics^15^, where the fixation of a mutation in a tumour is governed by the rate at which the event occurs in the tumour cell population and the selective advantage this event confers on the cell that acquires it. Under this framework, if the locus-specific mutation rate (*μ*) and selection coefficient (*s*) for a given tumour can be reasonably estimated, it should conceivably be possible to compute the probability (*P(D)*) of a driver mutation occurring at the locus and becoming fixed within the tumour population in subsequent time periods. However, robustly estimating these parameters presents technical challenges, especially in the context of DNA copy number alterations.

Traditionally, *μ* is expressed in terms of mutations per genome per cell division. However, estimating *μ* poses a unique challenge in the cancer setting, as the number of cells and cumulative cell divisions within a tumour over time can not be readily measured in a clinical context. Thus, various approaches (mainly limited to SNVs) have been developed to infer or approximate *μ*^*16–22*^. Our goal here is to infer *μ* in the context of the DNA copy number changes generated by chromosomal instability (CIN) that result in oncogene amplification or tumour suppressor deletion. As copy number changes can occur in a punctuated manner^23^, and those occurring at one cell division can be linked to further changes at subsequent divisions^24,25^, estimating a per cell division copy number mutation rate is extremely challenging. Therefore, we propose an approach to approximate the locus-specific mutation rate *μ* using a steady-state probability model that reflects the average likelihood of a locus-specific copy number change occurring over the lifetime of a tumour (**Figure 1a, Methods**). This obviates the need for detailed tracking of tumour cell division histories, albeit sacrifices temporal resolution. To calculate this probability, we first estimate the levels and mechanisms of CIN operating in the input tumour, represented by *I*_*CX*_. This is achieved using our previous framework for quantifying CIN that identifies and quantifies distinct patterns of copy number change representing the activity of different CIN processes that shaped the genome^26^. These CIN signatures include patterns representing mitotic errors, impaired homologous recombination and impaired non-homologous end-joining, among others^26^. However, not all of these types of CIN can generate focal amplifications or deletions^14,27–31^. Thus, we also estimate locus-specific CIN signature weights, *ω*_*CX,D*_, that represent the putative causal signatures which can amplify or delete the locus of interest. We learn these weights from examples of oncogene amplifications and tumour suppressor deletions found in TCGA. To enable this, we have adapted our CIN signature framework to assign CIN signatures to individual copy number events and used this to infer causal CIN types for different oncogene amplifications and tumour suppressor deletions. Recognising that local DNA topology features may constrain the possible copy number configuration affecting a given tumour amplification or deletion^32,33^, as part of this inference we also incorporate a correction factor learnt from passenger events scattered across the genome (see **Methods**). The product of the resulting corrected weights, *ω*_*D,CX*_, and the CIN signature activity estimated from the input genome, *I*_*CX*_, results in a steady-state probability approximation of *μ*, the locus-specific mutation rate (**Figure 1a**).

**Figure 1.**
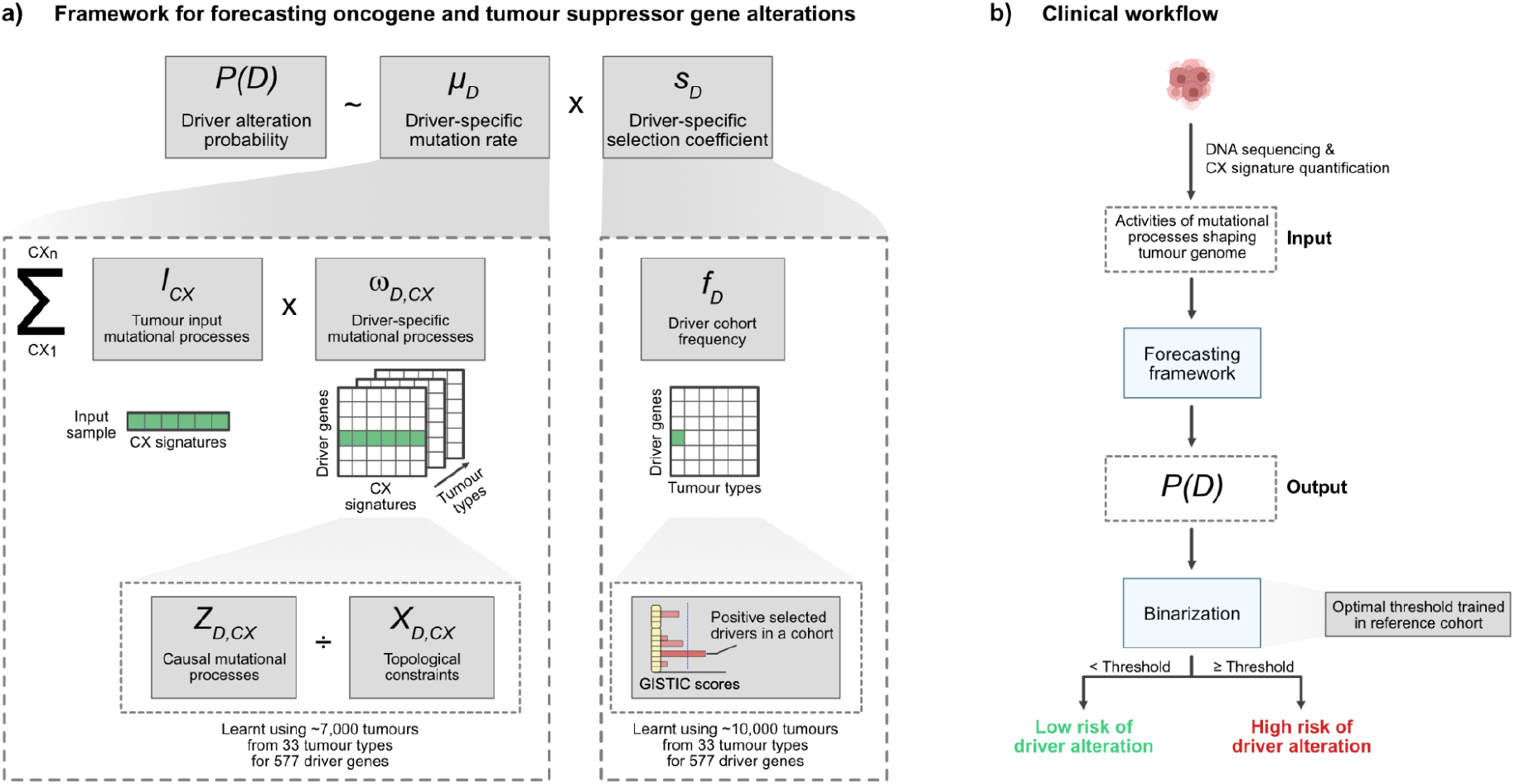
A framework for forecasting oncogene amplifications and tumour suppressor gene deletions. **a)** A schematic representation of our forecasting framework. The framework approximates the probability P(D) of a driver copy number alteration to be acquired and fixed within a tumour population in the future. This P(D) results from the product of the driver-specific mutation rate (μ_D_) and the driver-specific selection coefficient (s_D_) in a given tumour. μ_D_ is estimated taking into account the type of chromosomal instability (CIN) causing the focal amplifications and deletions of driver genes by combining the levels of CIN operating in the tumour (I_CX_) and the relative likelihood that a CIN type cause the driver alteration (ω_D,CX_). These driver-specific mutational processes result from factoring out the topological constraints (X_D,CX_) impacting on the configuration of observed copy number alterations caused by CIN (Z_D,CX_). s_D_ is approximated to the driver frequency at cohort level (f_D_) for drivers under positive selection (see Supplementary Note 1). **b)** Workflow for framework implementation. The framework takes as input the copy number signature activities (CX signatures) estimated from the input tumour and outputs the probability to acquire the amplification or homozygous deletion of interest. Then, an optimal threshold is used to classify the tumour as having high or low risk to acquire the given driver alteration in the future (see Supplementary Note 2 for details).

The selection coefficient, *s*, for a given mutation can traditionally be estimated by comparing growth rate differences between cells with and without the mutation. However, such measurements are not feasible in a clinical setting. Recent studies observing the emergence and fixation of key drivers at single cell resolution in premalignant lesions and tumours have facilitated the estimation of fitness effects for a limited number of drivers^34–38^. While these effects are specific for the conditions in which the cells were cultured or grown and may not represent physiological selective pressures, these studies do show that the estimated fitness effects often correlate with the frequency at which the driver is observed in human tumour populations. This suggests that driver mutation frequency in a cohort can serve as a proxy for *s* (see **Supplementary Note 1** for further discussion). Therefore, to approximate *s*, we used the frequency of oncogene amplification or tumour suppressor homozygous deletion in The Cancer Genome Atlas (TCGA) cohort, calculating these frequencies independently for each tumour type to account for tissue-specific selective pressures. Only confirmed driver genes^39–42^ showing evidence of positive selection, as determined by GISTIC^43,44^, were considered (**Figure 1a, Figure S1-S2**).

The product of the approximated values for *μ* and *s* for a given tumour yields *P(D)*, the probability that the driver mutation will arise and become fixed within the tumour at some (undetermined) point in the future. To enhance the clinical utility of this probability, we also developed methods to define thresholds for *P(D)*, enabling the prediction of a high or low risk that the tumour might acquire a focal amplification or deletion of a driver gene(s) in the future (**Figure 1b**). Further details on thresholding approaches to use for different application areas can be found in **Supplementary Note 2**. These strategies aim to translate *P(D)* into actionable forecasts, supporting risk stratification and personalised clinical decision-making.

### Model training and performance evaluation

We trained *ω*_*D,t,CX*_ and *s*_*D,t*_ for 352 oncogene-tumour type pairs and 52 tumour suppressor gene-tumour type pairs (**Figures S1-S2)** using 7,880 samples across 33 tumour types from the TCGA cohort. We then tested the performance of these models using two independent validation cohorts: The Pan-Cancer Whole Genome (PCAWG) project^45^, including 2,114 samples from primary tumours; and the Hartwig Medical Foundation (HMF)^46^, including 4,784 metastatic tumour samples. In total, we evaluated performance for 241 of the models with sufficient data for validation (≥5 samples with the target driver alteration in the tumour type of interest). To avoid target leakage during this evaluation, we removed *in silico* the amplification or homozygous deletion spanning the target driver from the tumour copy number profiles prior to forecasting (see **Methods** for details).

This retrospective evaluation showed that the framework often yielded models with the capacity to predict oncogene amplification or tumour suppressor homozygous deletion accurately, with an overall mean area under the receiving operating curve (AUC) greater than 0.7 (**Figure 2a, Table S1**). Performance in these independent cohorts was similar to the reapplication of the models to the training cohort (**Figure S3**), suggesting that models did not overfit training data. At the level of individual models, 147 models showed an AUC > 0.7, suggesting good predictive capacity. The remaining 94 models presented an AUC < 0.7 in either HMF or PCAWG cohorts, and as such we caution against their application without the use of further training data (**Supplementary Note 2**). We also validated the different thresholding approaches for binarised outputs (**Figures S4-S5**).

**Figure 2.**
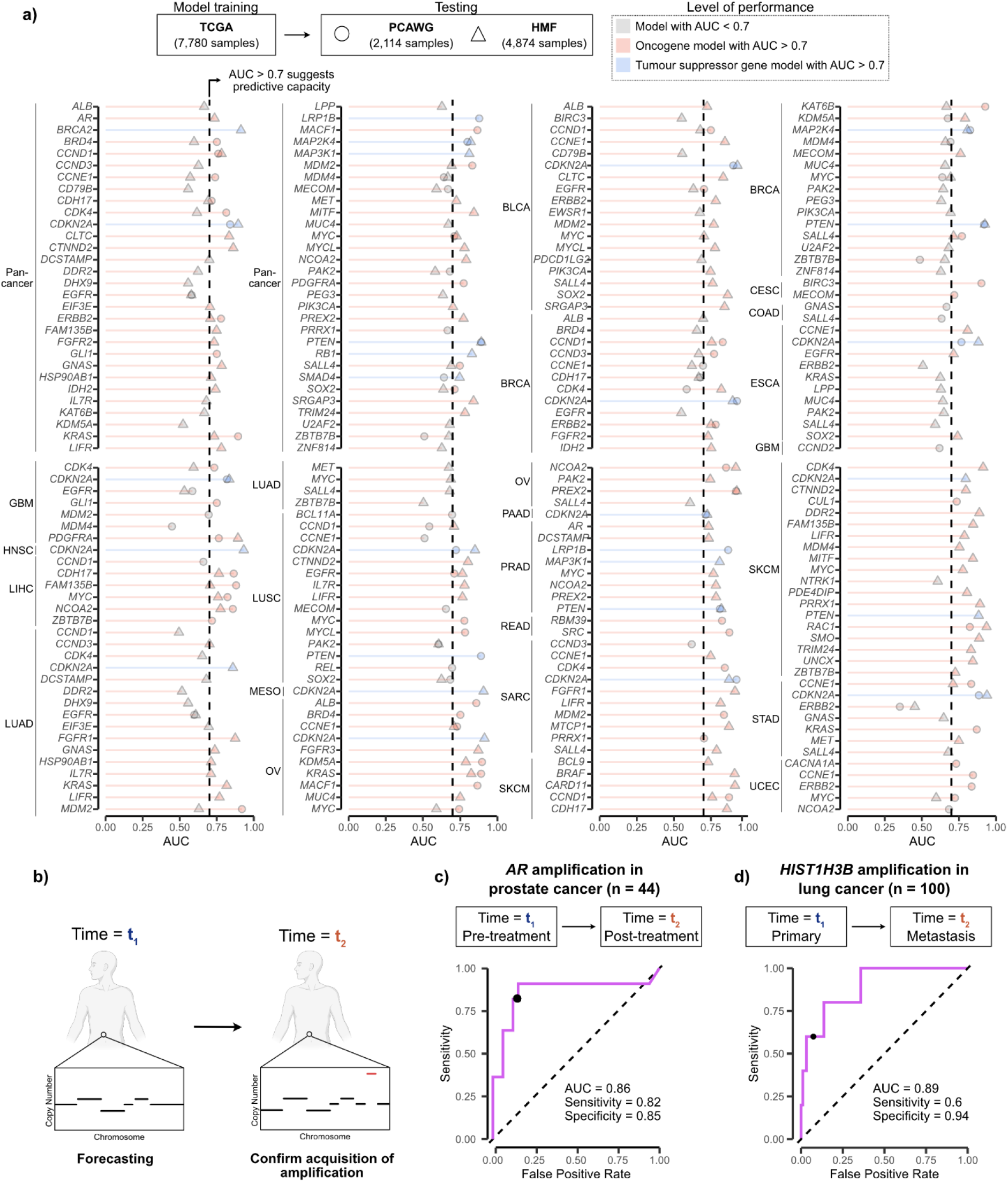
Forecasting performance. **a)** Area under the receiving operating curve (AUC) for 181 tumour-type and 60 pan-cancer forecasting models applied in the Hartwig Medical Foundation (HMF) and Pan-Cancer Analysis of Whole Genomes (PCAWG) datasets. Models were trained in the The Cancer Genome Atlas (TCGA). The probabilities of driver acquisition P(D) for each sample were derived by applying the forecasting models to copy number profiles in which the driver copy number alteration and its adjacent segments were removed to avoid target leaking. Forecasting performance was only evaluated in pan-cancer/tumour-specific HMF or PCAWG cohorts with at least 5 samples harbouring the driver alteration. Circles indicate AUC values in the PCAWG, while triangles denote AUC values in the HMF. Dashed line indicates AUC value of 0.7, the threshold for presumed predictive potential. Red colour denotes models for oncogenes with AUC > 0.7, blue colour highlights models for tumour suppressor genes with AUC > 0.7, and grey colour indicates models with AUC < 0.7. BLCA, bladder urothelial carcinoma; BRCA, breast cancer; CESC, cervical squamous cell carcinoma and endocervical adenocarcinoma; COAD, colon adenocarcinoma; ESCA, esophageal carcinoma; GBM, glioblastoma multiforme; HSCN, head and neck squamous cell carcinoma; LIHC, liver hepatocellular carcinoma; LUAD, lung adenocarcinoma; LUSC, lung squamous cell carcinoma; MESO, Mesothelioma; OV, ovarian serous cystadenocarcinoma; PAAD, pancreatic adenocarcinoma; PRAD, prostate adenocarcinoma; READ, rectum adenocarcinoma; SARC, sarcoma; SKCM, skin cutaneous melanoma; STAD, stomach adenocarcinoma; UCEC, uterine corpus endometrial carcinoma. **b)** Schematic representation of the approach to assess performance of the forecasting framework using longitudinally collected pairs of tumour samples. **c-d**) ROC curves showing performance of the forecasting framework for **c)** AR amplification in post-treatment prostate cancer samples, and **d)** HIST1H3B amplification in metastatic lung cancer samples. The black dot represents the optimal threshold learnt from a reference cohort (see Supplementary Note 2 for details of recommended thresholding estimation).

This performance, although promising, had a limitation in that we had to artificially alter the genome to remove evidence of any amplification/deletion in positive cases to create a gold standard. To further assess performance without this modification, we sought cohorts of longitudinally collected tumour pairs where we could apply our forecasting framework to samples collected at an early time point (without the driver event) and assess prediction performance using the status of samples collected at later time points (with or without the driver event) (**Figure 2b**). Most cohorts of paired samples had insufficient numbers of recurrent drivers for this purpose (**Table S2**). However, we found sufficient data to assess performance for amplification of two oncogenes: *AR* and *HIST1H3B. AR* amplification frequently drives resistance to androgen deprivation therapy (ADT)^47^, allowing us to forecast using pretreatment samples; and *HIST1H3B* amplification has been shown to be exclusively found in non-small cell lung cancer (NSCLC) metastases^48^, allowing us to forecast using primary samples seeding the metastatic sample. Applied to 44 longitudinal pairs of prostate cancers^46,49^, the *AR* model achieved a sensitivity of 0.82, a specificity of 0.85 and AUC of 0.86 for forecasting *AR* amplifications (**Figure 2c**). Applied to 100 primary-metastasis NSCLC pairs^48^, the *HIST1H3B* model achieved a sensitivity of 0.6, specificity of 0.93 and AUC of 0.89 for forecasting *HIST1H3B* amplification (**Figure 2d**). Importantly, alternative approaches to forecasting, for example, using the total number of amplifications, fraction of genome altered, or level of CIN, did not show predictive capacity (**Figure S6**).

### Forecasting poor prognosis

As oncogene amplifications or tumour suppressor gene deletions can drive cancer progression, forecasting whether a patient will acquire these events in the future may help identify early-stage patients with poor outcomes. To test this, we focused on low-grade glioma (LGG), where molecular testing is employed for diagnosis, prognosis and disease management. The current WHO Classification of Tumors of the Central Nervous System^50–52^ provides guidance for diagnosing and managing LGG based on a combination of molecular and pathological features. Focussing here on the molecular features, we provide a summary of the classification scheme: LGG tumours found to be *IDH* wild-type are upgraded to grade 4 glioblastomas with poor prognosis; *IDH* mutant tumours with 1p/19q codeletion are classified as grade 2 or 3 oligodendrogliomas with good prognosis; *IDH* mutant non-codeleted without *CDKN2A/B* deletion are classified as grade 2 or 3 astrocytomas with intermediate prognosis; and *IDH* mutant non-codeleted with a *CDKN2A/B* deletion are upgraded to grade 4 astrocytomas with poor prognosis. As a baseline for our cohort, we applied this classification to a combined patient cohort of 473 patients from the TCGA-LGG and Glioma Longitudinal AnalySiS^53^ (GLASS) consortium studies, performed a survival analysis, and reassuringly recapitulated prognosis across the classification groups (median survival of 12.9, 7.3, 3, 1.7 years for the good, intermediate and two poor prognosis groups, respectively, **Figure 3a**). The largest group was the *IDH*-mutant, non-codeleted, *CDKN2A/B* deletion wild-type, intermediate prognosis, with 51% of patients assigned. In contrast, the smallest group was the *CDKN2A/B* deleted, representing 1% of the cohort.

**Figure 3.**
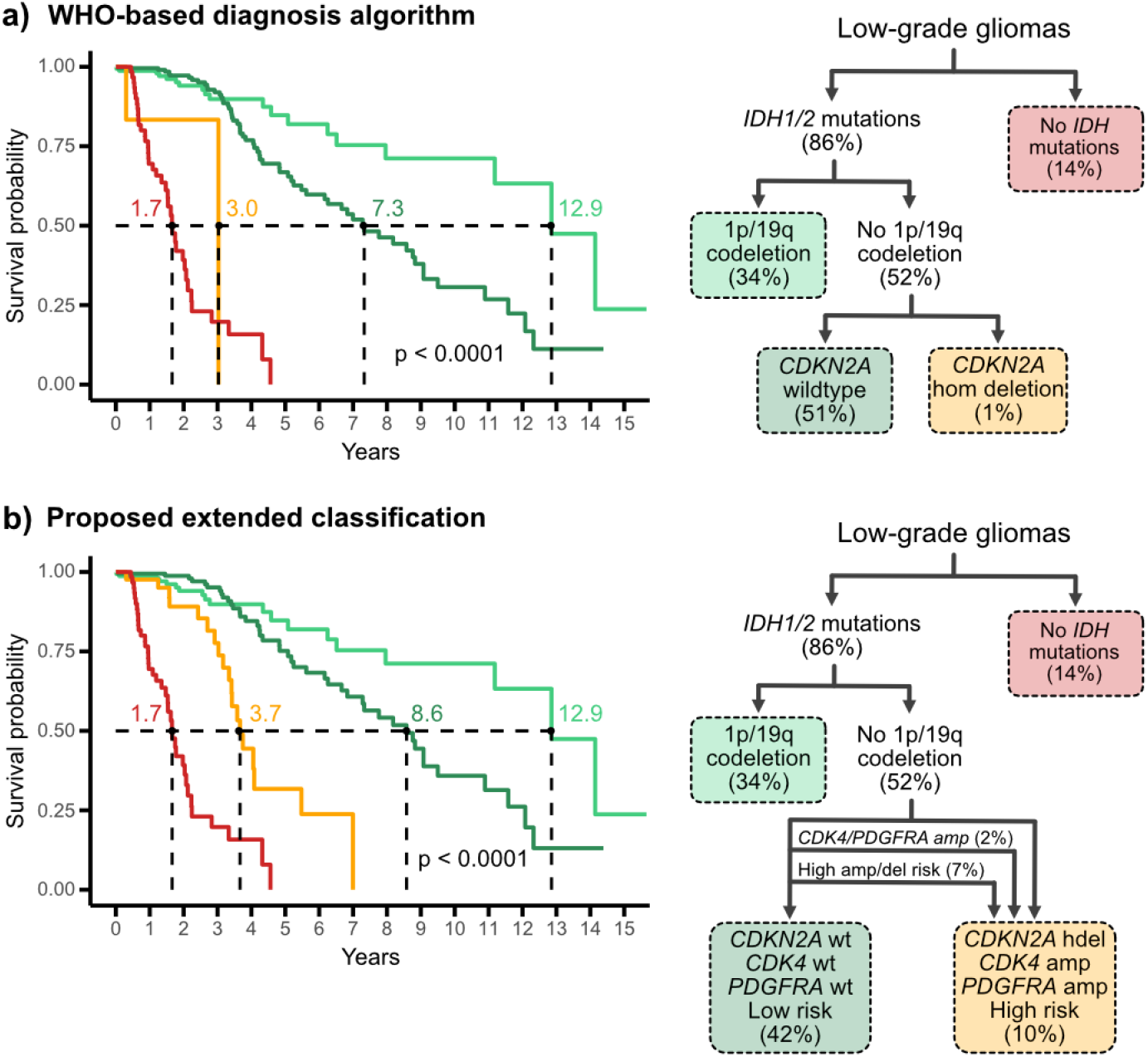
Predicting prognosis of low-grade gliomas. **a-b)** Kaplan-Meier curves showing overall survival for low-grade glioma (LGG) patients from TCGA and GLASS cohorts classified based on **a)** the 2021 World Health Organization Classification of Tumours of the Central Nervous System^51^, and **b)** the proposed extended classification of IDH-mutant-non-codeleted low-grade gliomas patients that includes the actual and predicted CDK4/PDGFRA/CDKN2A alteration status in the diagnosis algorithm. Lines are coloured according to the groups from the diagram in the right. Median survival values are indicated for each group. The p-value was estimated by the log-rank test. The prevalence of each low-grade glioma subtype is shown in brackets.

Recent studies have suggested that amplification of *CDK4* or *PDGFRA* can also indicate poor prognosis, offering an opportunity to identify further *IDH* mutant non-codeleted patients which could be upgraded to grade 4 astrocytomas with poor prognosis^54–56^. Indeed, if we reclassify those patients with these events from the intermediate prognosis group to the poor prognosis astrocytoma grade 4 group, we observe similar median survival estimates with 3% of patients now in this group (**Figure S7a**). This suggests that considering *CDK4/PDGFRA* amplifications may improve future classification of low-grade gliomas. We next hypothesised that our forecasting framework could be used to identify patients at high risk of acquiring *CDK4/PDGFRA* amplification or *CDKN2A* deletion, improving their classification and therefore clinical management. Applying our framework trained on the amplified and deleted cases to the 233 unaltered patients identified a subgroup of 37 patients with a high risk of future *CDK4/PDGFRA* amplification or *CDKN2A* deletion. This reclassification resulted in an increase to 10% of patients appearing in the poor prognosis grade 4 astrocytoma group, maintaining similar median survival estimates to the original WHO classification (**Figure 3b**). Unfortunately, we did not have data to confirm that these patients went on to acquire these amplifications or deletions in the future, however, Cox proportional hazards analysis showed these results to be independent of age at diagnosis and general chromosomal instability, suggesting the poor prognosis in these patients is likely attributable to the increased risk of acquiring these amplifications or deletions (**Figures S7b-S8**).

### Forecasting treatment resistance

Forecasting oncogene amplification has the potential to identify patients at risk of acquiring oncogene-driven drug resistance. To explore this, we focused on forecasting *MET* amplification in NSCLC which can drive resistance to EGFR tyrosine kinase inhibitors (EGFR-TKI) in 10-30% *EGFR*-mutant patients^57,58^. Fortunately, upon resistance to EGFR-TKIs, these patients can be treated with inhibitors against *MET*. However, this usually only occurs after resistance has emerged. We therefore reasoned that patients at high-risk of acquiring *MET* amplification might benefit from upfront inhibition of both EGFR and MET^60^, preventing this mechanism of treatment resistance.

In this instance, we did not have an appropriate training cohort of resistant patients for which to estimate a threshold on *P(D=MET)*. Therefore we opted to use an *in vitro* approach to estimate an appropriate threshold. We performed a long-term culture of 4 *EGFR*-mutant NSCLC cell lines in the presence of EGFR-TKI to track the emergence of *MET* amplification in resistant clones. Importantly, none of these cell lines had a detectable initial *MET* amplification. However, following treatment, 11 out of 12 (92%) resistant clones derived from HCC827 showed a *MET* amplification, while the remaining cell lines showed no evidence of *MET* amplification-driven resistance (**Figure S9**). Our framework applied to the genomes of these 4 cell lines prior to treatment assigned the highest probability to the HCC827 parental cell line (**Figure 4a**). A threshold of 0.0031 was selected to accurately classify these cell lines. This threshold was similar when estimated in an independent long-term culture of the HCC4006, PC9 and HCC827 lines treated with the EGFR-TKI erlotinib^59^ (**Figure 4a, Figure S10**).

**Figure 4.**
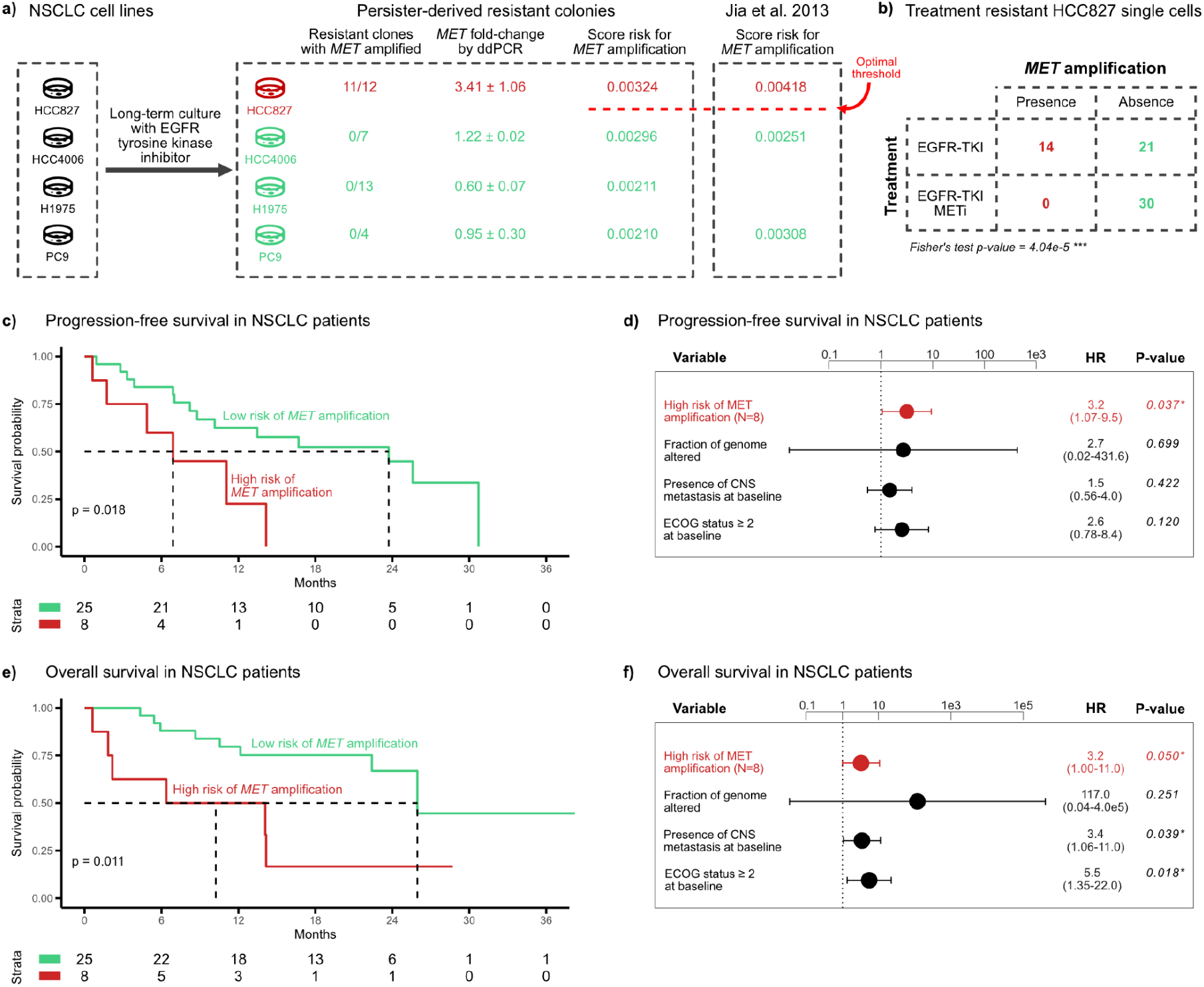
Predicting EGFR tyrosine kinase inhibitor resistance. **a)** Experimental validation of predicting EGFR tyrosine kinase inhibitor (EGFR-TKI) resistance via MET amplification using a panel of non-small cell lung cancer (NSCLC) cell lines treated with EGFR-TKI until the emergence of treatment resistance. MET amplification scores were computed in parental cell lines, and the presence of MET amplification in resistant clones was tested via digital droplet PCR (ddPCR). Red colour indicates resistance driven by MET amplification. **b)** Contingency table showing the number of cells acquiring and not acquiring MET amplification after treating the HCC827 parental cell line with EGFR-TKI alone or in combination with a MET inhibitor (METi). P-value was estimated by Fisher’s exact test. **c**,**e)** Kaplan-Meier curves showing **c)** progression-free survival and **e)** overall survival for NSCLC patients predicted as having low (green) or high (red) risk of acquiring MET amplification as the resistant mechanism to osimertinib. P-value was estimated by the log-rank test. **d**,**f)** Cox proportional hazards regression models showing **d)** progression-free survival and **f)** overall survival for NSCLC patients predicted as having low (green) or high (red) risk of acquiring MET amplification as the resistant mechanism to osimertinib. The model was corrected for the fraction of genome altered, the presence of metastasis in the central nervous system (CNS) at treatment initiation, and the physical status based on the Eastern Cooperative Oncology Group (ECOG) scale at treatment initiation. The threshold for classifying the patients was obtained from our in vitro study in **a)**.

Next, we wanted to test if treatment upfront with both *EGFR* and *MET* inhibitors could prevent *MET*-driven resistance *in vitro*. We treated the HCC827 parental cell line with erlotinib and a selective inhibitor of *MET*, harnessed resistant clones and performed single-cell shallow Whole Genome Sequencing (sWGS) to detect *MET* amplification. The application of both inhibitors to the HCC827 line resulted in an absence of resistant cells exhibiting *MET* amplification. In contrast, 40% of the resistant cells treated with only the EGFR-TKI showed *MET* amplification (p-value = 4.04e-05; Fisher exact test; **Figure 4b, Figure S11**). These findings indicate that our forecasting framework could be used to stratify patients for dual inhibition treatment strategies, thus preventing the emergence of this resistance mechanism.

To assess feasibility in a clinical setting, we collected a cohort of 68 *EGFR*-mutant NSCLC patients treated with the EGFR-TKI osimertinib (**Figure S12**). Whole-Exome Sequencing (WES) and sWGS of tumours collected at diagnosis showed an absence of *MET* amplification prior to treatment for all 33 patients with high-quality sequencing data. We applied our framework to these primary, treatment-naive tumour samples to forecast *MET* amplification using the threshold established from lung cancer cell lines, and integrated clinical data to assess osimertinib response. Eight cases were predicted to have a high risk of future *MET* amplification of which 6 were classified as non-responders via RECIST criteria. This subset of patients, representing 46% of the osimertinib resistant population, had shorter progression-free survival intervals compared to low-risk of amplification patients (median 6 vs 24 months, HR = 3.2, p-value = 0.037; independent of the fraction of genome altered, the presence of metastasis in the central nervous system (CNS) at treatment initiation, and the physical status based on the Eastern Cooperative Oncology Group (ECOG) scale at treatment initiation; **Figure 4c-d**) and shorter overall survival (median 10 vs 26 months, HR = 3.2, p-value = 0.050; independent of the fraction of genome altered, the presence of CNS metastasis at treatment initiation, and the ECOG status at treatment initiation; **Figure 4e-f**). This suggests patients predicted as high risk of *MET* amplification may indeed be suitable for upfront treatment with dual inhibitors of EGFR and MET.

## Discussion

One of the greatest challenges for precision medicine is the adaptive nature of tumours, which has so far been challenging to predict. Here, we present an approach to anticipating the emergence of tumour-suppressor gene homozygous deletions or oncogene amplifications, and how it may contribute to forecasting tumour progression and/or treatment resistance.

Our framework models established evolutionary forces using a modular structure. It combines the activity of mechanisms driving copy number changes in a tumour sample with their propensity to hit certain genomic locations, thus capturing the tendency of a tumour to amplify or homozygously delete a specific gene. The ability of the framework to integrate multiple mutational processes simultaneously is key, given the diverse mechanisms whereby we (**Supplementary Note 5**) and others have found driver genes to be amplified or homozygously deleted across different tumour types. By further combining this information with the selective advantage conferred by the driver copy number alteration, it aims to forecast whether a cell with such alteration is likely to clonally expand. The ability of the framework to adequately capture these factors is dependent on high-quality training and input data. The more similar the technical, clinical and biological properties of the training data are to the scenario where the user aims to deploy the framework, the better the framework will perform. Thus, we provide a set of guidelines to help users navigate how best to train and apply the framework (see **Supplementary Note 2**). These guidelines also include an important discussion on how to establish an appropriate threshold to generate binary predictions. The ability to reduce the complexity of forecasting to a binary prediction may simplify workflows for potential future clinical adoption.

To explore the clinical utility of this forecasting framework, we presented two suggestive examples of how forecasting specific genetic changes can unlock new clinical opportunities. First, in the context of risk stratification, we showed that up to 10% of low-grade glioma patients could be reclassified under extended WHO molecular prognosis guidelines as poor prognosis given their high risk of acquiring *CDK4/PDGFRA* amplifications or *CDKN2A* homozygous deletion. While longitudinal biopsies are warranted to confirm future acquisition of these alterations, these results highlight the capacity to forecast tumour progression via some of the models herein presented. Second, our framework also offers opportunities to enable new treatment modalities. Currently, NSCLC patients resistant to EGFR inhibition via *MET* amplification only receive second-line MET inhibitors once resistance has been confirmed. As our framework provides a method to identify patients at high risk of *MET* amplification at first-line, it can be used as a tool to stratify patients for treatment with new dual inhibitors of both EGFR and MET. New trials have established a 30% reduction in risk of progression when treating first-line with a dual inhibition of EGFR and MET, however, this comes at the cost of high toxicities^60^. We hypothesise that our predictor could prevent exposure to combination-specific toxicities in patients with a low risk of *MET* amplification who might derive similar benefit from monotherapy. While our *in vitro* results support this strategy, its application in patients warrants further investigation, as our clinical proof-of-concept lacked a follow-up biopsy at relapse to confirm *MET* amplification. Future trials could address this limitation by incorporating liquid biopsies to monitor the emergence of *MET* amplification^61^. Nevertheless, given that resistance to targeted therapies can arise from diverse oncogene amplifications^47,62^, predictive modeling could extend beyond the *MET* use case presented here. For instance, it could be applied to forecast resistance to ADT via *AR* amplification in prostate cancer or to stratify patients for novel therapies targeting oncogene amplifications.

When constructing our forecasting framework, we made an explicit design choice to use only tumour DNA copy number profiles as input. The advantage of this approach is its low cost and clinical utility, potentially facilitating forecasting using only a standard tumour genome sequencing test. However, as multiomic testing becomes more commonplace in the clinic, it may be possible to improve the estimation of the parameters of the forecasting framework and increase its complexity. Additionally, in this study, we assumed homogeneous selection pressures across patients within the same tumour type, but there is the opportunity to introduce complex epistatic interactions in our model that could improve forecasting accuracy by assigning fitness effects on a per-patient basis. We also assumed that the CIN processes identified using our CIN signatures are ongoing and contributing to the future potential of acquiring an amplification. However, some of these processes may be historical and no longer active. Improved profiling of tumours at single-cell resolution may help distinguish ongoing from historical CIN and potentially improve forecasting performance.

Overall, our results suggest that anticipating genomic progression is achievable and can ultimately be harnessed for improved clinical decision-making.

## Methods

See Methods.

## Supplementary Information

Supplementary Information contains Supplementary Notes 1-5, and Figures S1-S36

## Acknowledgements

First and foremost, we thank cancer patients and their families who generously contributed to this study. We thank Maria J Garcia, Oscar Llorca and Duncan Odum for valuable feedback on the manuscript. A.F-S., B.H., A.C., D.G-S., M.E-R., B.C-U., J.T., M.To., P.G.S. and G.M. are hosted by the Centro Nacional de Investigaciones Oncológicas (CNIO), which is supported by the Instituto de Salud Carlos III and recognized as a ‘Severo Ochoa’ Centre of Excellence (ref. CEX2019-000891-S) by the Spanish Ministry of Science and Innovation (MCIN/AEI/10.13039/501100011033). This work was supported by grants to G.M. by the Spanish Ministry of Science and Innovation (PID2019-111356RA-I00 funded by MCIN/AEI /10.13039/501100011033 and PID2022-137042OB-I00 funded by MCIN/AEI /10.13039/501100011033 and by the European Union, ERDF “A way of making Europe”). A.F-S. received the support of a fellowship from “La Caixa” Foundation (ID 100010434; LCF/BQ/DR21/11880009). B.H.’s postdoctoral contract was supported by philanthropists via the ‘Amigos/as del CNIO’ Programme. B.H. was also supported by ‘La Caixa’ Foundation (ID 100010434; LCF/BQ/PR23/11980033). B.C-U. received the support of a fellowship from “La Caixa” Foundation (ID 100010434; LCF/BQ/DR23/12000023). M.T. was supported as a postdoctoral researcher of the F.R.S.-FNRS. T.L. was supported by the Francis Crick Institute which receives its core funding from Cancer Research UK (CC2008), the UK Medical Research Council (CC2008), and the Wellcome Trust (CC2008). J.M.C.’s postdoctoral research contract was supported by Comunidad de Madrid via Programa de Atracción de Talento (2018-T2/BMD-11945). The OSIRESP clinical study was funded by AstraZeneca (ESR-20-20709). The Pan Prostate Cancer Group was supported by the Dallaglio Foundation. This publication and the underlying study have been made possible partly based on data that Hartwig Medical Foundation has made available to the study through the Hartwig Medical Database.

## Author contributions

A.F-S. and B.H. contributed equally to this work. G.M., B.H. and A.F-S. conceived and designed the study. A.F-S. and B.H. developed the methodology and software of the study. D.G-S., A.C., P.G.S., M.E-R., B.C-U., J.T., M.To., L.P-A., J.Z., V.A., E.A., G.R.dG., J.M.C, T.L., M.T. and The Pan Prostate Cancer Group provided access to data and/or contributed to gathering, processing and curating data. A full list of group members can be found in Supplementary File 1. A.F-S., B.H., G.M. and D.G-S. contributed to the formal analysis presented in this study. A.F-S., B.H. and G.M. wrote the manuscript. A.F-S., B.H. and G.M. produced and contributed to the visualisations of the study. G.M supervised the project. All authors had access to all of the data in the study. All authors contributed to the review and the editing of the manuscript. All authors approved the manuscript before submission.

## Declaration of interests

G.M. is co-founder, director and shareholder of Tailor Bio Ltd. J.Z. has served as a consultant for Sanofi, Pfizer, AstraZeneca, BMS, Novartis, NanoString, and Guardant Health. J.Z. reports speakers honoraria from Pfizer, BMS, Roche, AstraZeneca, NanoString and Guardant Health, all outside the submitted work. J.Z. receives grant support from AstraZeneca, Roche, and BMS. L.P-A. has received honoraria for scientific advice and speaker fees from Lilly, Merck Sharp & Dohme, Bristol-Myers Squibb, Roche, PharmaMar, Merck, AstraZeneca, Novartis, Boehringer Ingelheim, Celgene, Servier, Sysmex, Amgen, Incyte, Pfizer, Ipsen, Adacap, Sanofi, Bayer and Blueprint, and participates as an external member of the board of Genómica. L.P-A. is founder and board member of Altum sequencing and has received institutional support for contracted research from Merck Sharp & Dohme, Bristol-Myers Squibb, AstraZeneca and Pfizer. The Spanish National Cancer Research Centre (CNIO) have filed a patent application (EP23383179.1) covering the methodology for assigning copy number signatures to individual copy number events that lists B.H. and G.M. as inventors. CNIO and the Foundation of the Health Research Institute Hospital 12 de Octubre have filed a patent application (EP23383180.9) covering the forecasting framework methodology and clinical applications that lists A.F-S., B.H., D.G-S., L.P-A., J.Z., J.M.C. and G.M. as inventors.

