## Supplementary Information for "Forecasting oncogene amplification and tumour suppressor deletion"


**Supplementary Text** **2**

Supplementary Note 1 - Leveraging the approximate linear relationship between s and f 2

Supplementary Note 2 - Guidelines for forecasting 9

Supplementary Note 3 - On the feasibility of forecasting oncogene amplifications 12

Supplementary Note 4 - Particularities of tumour suppressor genes homozygous deletions 15

Supplementary Note 5 - Exploring the mutational processes putatively causing oncogene amplifications and tumour suppressor deletions 16

**Supplementary Figures** **18**

**References** **49**

### Supplementary Text

#### Supplementary Note 1 - Leveraging the approximate linear relationship between *s* and *f*

For an individual tumour, the fixation of a mutation in a driver gene (amplification, deletion or otherwise) depends primarily on the product of two key factors: (1) the rate at which the alteration occurs in the tumour cell population (mutation rate, $\mu$), and (2) the selective advantage this alteration confers on the cell that acquires it (selection coefficient, *s*). Given this, deriving an estimate for *s* for a given tumour is challenging as it is confounded by $\mu$. However, at a tumour population level, recent studies (outlined below) suggest that the effect of $\mu$ can be "averaged out". This implies that there may be a relationship between the frequency, *f,* at which a mutation occurs in the population and *s*. Indeed, these studies show an approximately linear relationship between *s* and *f.* This provides an opportunity to use *f* in a population of tumours to approximate *s* in our framework. Below, we provide a further theoretical justification for the approximate linear relationship between *s* and *f.*

##### Previous evidence of an approximate linear relationship between *s* and *f*

There are examples in the literature that provide evidence for an empirical direct relationship between the *s* and *f*.

In Watson *et al*. 2024[10] an *in vitro* forward genetic screen was employed to observe selection of aneuploid states. This study used a starting population of breast cells with induced aneuploidy (all possible chromosome loss/gain copy number alterations (CNAs), equivalent to a constant mutation rate). These were allowed to evolve in proliferative competition assays. The resulting fit clones, across replicates, showed evidence of selection of particular CNAs, i.e. those which were recurrent. The frequency at which these recurrent CNAs were observed across the clones correlated with the frequency of CNAs observed across an independent population of breast cancers (Figure 1 in Watson *et al*. 2024[10]). This suggests that in this instance, the population frequency of a CNA approximates the selection coefficient attributed to that CNA, at least within the same tumour type.

Davoli *et al*. 2013[11] modeled the functional impact of arm-level CNAs by integrating the density and potency of tumour suppressors, essential genes, and oncogenes within a chromosome arm. By jointly assessing these genes, they assign a functional score to each arm, serving as a surrogate for CNA functional impact and fitness. This score correlated strongly with the relative frequency at which the corresponding chromosome arm is lost or gained (Figure 5 in Davoli *et al*. 2013[11]).

Shih *et al*. 2023[12] present a method to disentangle mutation rate effects from selection when estimating the relative fitness impact of CNAs. Rather than analyzing recurrence, they infer peak-level relative fitness by examining deviations in the distribution of telomeric- and centromeric-bounded CNA (*partial CNA*) lengths (Figure 1). Their key hypothesis is that positively selected loci are more frequently altered by *partial CNAs*, while negatively selected loci are less frequently affected. This approach enables relative fitness estimation independent of mutation rate, relying solely on deviations between the observed and expected distributions of partial CNA configurations. Extending these peak-level relative fitnesses to arm-level relative fitnesses, Shih *et al*. identify a strong relationship between chromosome arm relative fitness and the frequency of its gain or loss across tumors (TCGA, Figure 4b, Figure 4c in Shih *et al*. 2023[12]), a trend consistent across tumor types (Figure 4e in Shih *et al*. 2023[12]). This shows that an estimation of relative fitness that theoretically uncouples a potential bias generated by mutation rate offers similar and proportional estimates to the relative frequencies used in this study.

Dinh *et al*. 2024[13] present a mathematical framework for modeling karyotype diversification and selection during tumour evolution. Assuming uniform missegregation rates across chromosomes and patients, the authors used approximate bayesian computation to fit the arm selection parameters and the chromosome missegregation probability to PCAWG and TCGA CNA data. In Figure 2d, 2e, 2f and 2j, Dinh *et al.* showed that the selection rate of losses and gains strongly and linearly correlated with the frequency of loss or gain.

#### A theoretical argument supporting the approximate linear relationship between *s* and *f*

Through a mathematical modelling approach we aimed to investigate how both the mutation rate and the selection coefficient of a given driver, *D*, contribute to its final cohort frequency, in order to support that both contributions can be disentangled. To establish such a model, we make the following assumptions:

- At the cohort level, as a first approximation, we assume that driver mutations with a strong selection coefficient confer a similar selective advantage across all tumours, regardless of patient-specific factors. Consequently, the driver selection coefficient is uniform across patients and remains constant with age and tumour stage.
- We assume that the overall mutation rate varies between patients but is constant for the tumour lifespan.
- We assume that the patient cohort is representative enough to consider that patient variability in mutation rate, age, cell division rate, and locus-specific mutation rate can be approximated by a consistent probability density function (PDF).

To construct our model, we first establish a representation of the effective mutation rate in an individual tumour. Here, each patient $i$ has their own:

- $M_i$: overall mutation rate (number of mutations per cell division).
- $T_i$: tumour *age* (generations elapsed since the first tumour cell).
- $r_i$: cell division rate (related to the average number of daughter cells per cell division, accounting for both birth and death rates).
- $\mu_{i,D}$: driver locus mutation rate (likelihood of acquiring mutations specifically at the genomic region where the driver is located).

Altogether, these variables can be consolidated into a single parameter that we call $\Lambda_{i,D}$ which indicates the number of opportunities for a specific driver alteration to arise, and depends on both the mutation rate and the tumour growth dynamics and stage. It can be computed as:

$$\Lambda_{i,D} = \mu_{i,D} \times M_i \times N_i \times T_i$$

where $N_i$ is the *effective tumour size*, i.e. the time-averaged number of tumour cells. For a cell population undergoing exponential growth with rate $r_i$, this is calculated as the temporal harmonic mean of the population size[14]:

$$N_i = \frac{T_i}{\int_0^{T_i} e^{-r_i t} dt} = \frac{r_i T_i}{1 - e^{-r_i T_i}} \approx r_i T_i$$

Now, since we have variability in $M_i$, $\mu_{i,D}$, $T_i$, and $r_i$ across our patient cohort, the resulting values of $\Lambda_D$ follow a distribution, denoted as $p(\Lambda_D)$, which gives the relative amount of patients with a certain value of $\Lambda_D$.

Using this, we can compute the probability that a single tumour harbours a mutation. Suppose we know the value of $\Lambda_{i,D}$, the number of opportunities that driver $D$ had to arise in a particular patient $i$ by the time of sampling. For notation purposes, we use $\Lambda$. Once the mutation of interest appears in one cell of the population, its associated selection coefficient will determine its *fixation probability*. This is the probability that, once a given alteration appears in the population, it will fix (all cells will eventually carry it), and its definition depends on the evolutionary model under consideration. For example, in the (haploid) Wright–Fisher model, this probability can be calculated as[15]:

$$\rho(s) = \frac{1 - e^{-s}}{1 - e^{-Ns}}$$

where $s$ is the selection coefficient (defined for this instance in the context of the Wright–Fisher model) and $N$ is the *effective tumour size*. For the purposes of our work, we can assume that the effective tumour size is large enough so that:

$$\rho(s) \approx 1 - e^{-s}$$

and $s$ in our context can be referred to as either “selection coefficient” or “fixation probability”.

We can expand this result to the case where an alteration may appear multiple times during evolution. Assuming that occurrences of the mutation can be considered independent events and are approximately Poisson($\Lambda$) distributed, the probability that no mutant ever manages to clonally expand is equal to $e^{-\Lambda s}$. Hence, the probability that the mutation has been fixed within the tumour at tumour sampling is:

$$P(\text{D fixes}|\Lambda, s) = 1 - e^{-\Lambda s}$$

This probability is defined for a single patient. To extend this argument to a cohort of patients and calculate the fraction of them where the driver is clonally mutated, we have to average this probability over the distribution of $\Lambda$. Given that there are different values of $\Lambda$ in the cohort with relative proportions $p(\Lambda)$, this average is obtained by integrating:

$$f(s) = \int_0^\infty \left[1 - e^{-\Lambda s}\right] p(\Lambda) d\Lambda$$

Although the explicit definition of $p(\Lambda)$ is unknown, we can try to put a prior on it to continue with this analysis. Once this integration is solved, we can estimate the value of $s$ given an observed value of $f$.

Because we consider the accumulation of mutations as a sequential Poisson process with rate $\Lambda$, the distribution of the number of such mutations across many patients is, consequently, also Poisson-distributed[16,17]. Let us then assume that $p(\Lambda)$ can be fitted to a Poisson distribution with parameter $\lambda$:

$$p(\Lambda) = \frac{e^{-\lambda} \lambda^{\Lambda}}{\Lambda!}$$

with $\lambda$ representing the cohort average of $\Lambda$. Then, the previous integral becomes solvable:

$$f = 1 - \exp(-\lambda s)$$

Therefore,

$$s = -\frac{1}{\lambda} \ln(1 - f)$$

which is nearly linear in the biologically relevant regime of $\lambda s \ll 1$.

Note from these expressions that we have been able to decouple patient-specific aspects such as patient predisposition to acquire a specific driver, or patient age at diagnosis, from the resulting cohort frequency at which the driver is present. Indeed, the frequency dependence on the mutation rate and tumour history is manifested only through the cohort average $\lambda$, for which we can estimate reasonable lower and upper bounds from real life data (see below).

We test this theoretical model by choosing parameter values that approximate biological scenarios derived from our analysis on a TNBC cohort (**Supplementary Note 1**) where we extracted features of unique focal amplifications. We computed the average effective arrival rate parameter using the definition of $\Lambda_{i,D}$ and substituting:

$$\lambda = \frac{3 \times 10^6}{3234000000} \times \frac{13.8}{1000} \times T$$

We assume a locus-specific mutation rate of $1/n_{\text{loci}}$ (the expected locus-specific mutation rate assuming uniform burden across the genome) where $n_{\text{loci}}$ is an estimate of the number of regions susceptible to amplifications and is estimated as the reference genome length,

$3234000000$ bp, divided by the average segment size of the focal amplifications observed in the TCGA cohort, $3 \times 10^6$ bp.

The overall amplification rate $M$ for the cohort is estimated from the $13.8$ amplifications per $1000$ cells found for the TN4 TNBC tumour samples analysed in **Supplementary Note 1**. This corresponds to the maximum number of alterations per tumour type observed in this cohort, thus providing an upper bound on $\Lambda$ and therefore $\lambda$. For an exponentially growing cell population with net growth rate $r$ for $T$ cell cycles, the expected number of unique amplifications that can be found in a sample of $n$ cells is approximately:

$$\mathbb{E}[\#\text{ unique amplifications}] = nrTM$$

, where $rT$ can be understood as the length (in cell divisions) of each of the $n$ external branches in the sampled cells' history. Hence,

$$M = \frac{\mathbb{E}[\#\text{ unique amplifications}]}{nrT} = \frac{13.8}{1000} \cdot \frac{1}{rT}$$

Combining these parameter values is how we find the formula for $\lambda$ as a function of the only unknown parameter in this instance: $T$. Note that $\lambda$ no longer depends on the cell growth rate but only on the mutation rate and tumour age. To finalize this argument, in **Supplementary Note Figure 1** we vary $T$ in a realistic range of tumour ages[18], that span from ~30 cell divisions (corresponding to $\lambda \approx 4 \cdot 10^{-4}$) up to ~1500 (corresponding to $\lambda \approx 2 \cdot 10^{-2}$).

**Supplementary Note Figure 1** illustrates how $s$ can be estimated from $f$ at a fixed cohort average value of $\lambda$ for the Poisson prior on $p(\Lambda)$ and shows that the inferred $s$ indeed can be accurately approximated from the linear correlation assumption $s = f/\lambda$ (dotted lines). In all cases depicted, the curves illustrating $s$ as a function of the cohort frequency $f$ are monotonically increasing: as $f$ grows, so does the inferred selection coefficient $s$. Notice that when the average mutation rate and tumour age increases, the distribution of $\Lambda$ skews toward higher values, and the gap between the curve of $s$ vs. $f$ and the linear approximation $s = f/\lambda$ becomes more pronounced. Thus, larger $\lambda$ values push the inferred $s$ to deviate more strongly from $f/\lambda$, underscoring the interplay between mutation rate (captured by $\Lambda$) and selection in determining the observed cohort frequency. However, this discrepancy remains small for the biologically relevant regime in which our work is focused (frequencies from 0.001 up to 0.5 and the mutation rates calculated above).

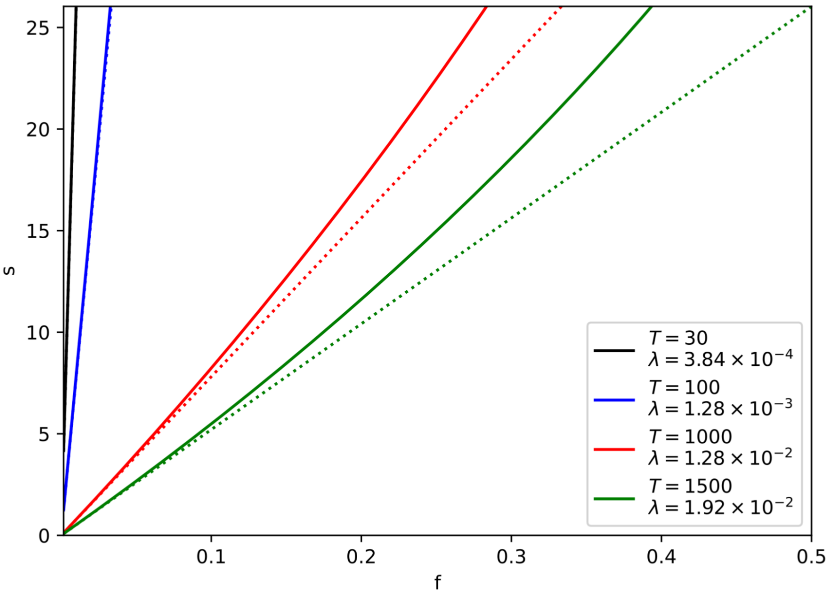


**Supplementary Note Figure 1.** Dependence of $s$ with the cohort frequency $f$. Dotted lines indicate our assumption in the main text that $s$ is linearly correlated with $f$ with slope $1/\lambda$. Different parameter values are proposed, corresponding to different plausible tumour ages[18] (expressed in number of tumour cell divisions) and mutation rates from the cohort of TNBC samples analysed and described in **Supplementary Note 3**.

Our analysis shows that the cohort frequency, $f$, is determined by the interplay between the mutation rate, the tumour growth dynamics (these two combined in what we call $\Lambda$), and the selective advantage $s$. In practice, the overall heterogeneity in tumour-specific parameters (mutation rate, age, cell division rate, and locus-dependent predisposition) forces us to integrate over a distribution $p(\Lambda)$. Because the precise form of $p(\Lambda)$ is unknown, we have assumed a Poisson-distributed prior. In this case, the resulting mapping between $f$ and $s$ is nearly linear in the biologically relevant regime of $\lambda s \ll 1$, where

$$s \approx \frac{f}{\lambda}$$

A central concern in estimating selection coefficients from tumour frequencies is that the observed cohort frequency $f$ is jointly determined by $\lambda$—the cohort-average product of the driver specific mutation rate and the tumour growth stage—and by $s$—the selection coefficient. One may point out that a high observed frequency could be driven by a high mutation rate alone, rather than by a genuinely large selective advantage. In that view, reading a high $f$ as evidence for a high $s$ risks being an over–simplification of the problem.

To address this concern, we explicitly incorporate a distribution $p(\Lambda)$ over the cohort of tumours and examine how $s$ depends on the resulting mean frequency $f$. **Supplementary Note Figure 1** illustrates the results of this analysis for several plausible upper bound values for $\lambda$. Over these parameter sweeps, we find that the inferred $s$ remains nearly linear with $f$

in magnitude, indicating that assuming $s \approx f$ (subjected to renormalization) does not drastically overstate a driver's selective advantage in typical TNBC-like conditions. Importantly, the risk of overestimating $s$ (i.e. attributing a high frequency solely to a high selective advantage when in reality a high $\Lambda$ might be responsible) is typically modest in biologically relevant parameter ranges. In most regimes of tumour sizes and mutational rates, the predicted $s$–vs–$f$ curves remain close to the linear approximation.

Additionally, note that the frequency $f$ is mathematically expressed as the product of two independent functions of $\Lambda$ and $s$, respectively. This factorization implies that if independent estimates or assumptions on the driver locus-specific mutation rate are available (for example, across several drivers within the same patient cohort), one can effectively renormalize the problem to remove the dependency on $\Lambda$. In doing so, the contribution of the mutation rate is accounted for using empirical data, leaving the selection coefficient $s$ as the sole free parameter governing the cohort frequency. This renormalization provides additional reassurance that our estimation of $s$ from $f$ is robust and not confounded by differences in mutation rates among patients.

To conclude, our results indicate that, while the observed frequency $f$ reflects the combined effects of mutation rate and selective advantage, integrating over a plausible distribution for $\Lambda$ yields a mapping from $f$ to $s$ that is approximately linear for reasonable cohort-average mutation rate values. Hence, the possibility of misinterpreting high frequencies as solely due to high mutation rates is minimized. These points, along with the ability to renormalize the problem using independent mutation rate information, support the empirical assumption $s \approx f$ as a useful and conservative approximation for estimating the driver's fitness effect from tumour sequencing data.

### Supplementary Note 2 - Guidelines for forecasting

Our forecasting framework can be used to answer both genomic and clinical research questions. In the genomics setting, a typical use case might be the application of the framework to a sample to determine which oncogene amplification or tumour suppressor gene homozygous deletion is likely to drive the future evolution of the tumour (if any). In this instance, the framework can output a list of oncogenes and tumour suppressor genes at high risk of future amplification or homozygous deletion, respectively. Default framework parameters and corresponding thresholds learnt using TCGA samples can generally be used, particularly if the forecasting model shows predictive capacity in independent validation using PCAWG or HMF cohorts (**Figure 2, Table S1**). However, the user needs to take care to ensure that the local mutation rates and selection coefficients across genes are comparable. In a clinical research setting, the framework is more likely to be used with a specific gene in mind. In this case, the target gene is specified upfront and the framework should be trained on appropriate data and the optimal threshold set accordingly. Here, we provide suggestions on how to use the framework in both research settings.

#### What do I need as input?

The framework takes as input an unrounded, absolute copy number profile of the tumour genome. These profiles can be obtained by profiling the genome using SNP arrays, whole-exome or whole-genome sequencing and applying software such as ASCAT, ABSOLUTE, or our own approach described in the "Generating copy number profiles" section in the Methods. It is particularly important to ensure the resolution of this profile is suitable (ideally 30-50 kb) for quantifying CIN signatures activities, the input $I_{CX}$ to the forecasting framework. ASCAT processed SNP6-derived profiles can be input natively, however, we recommend a 30-50 kb bin resolution for whole exome and whole genome sequencing (**Figure S18**). The detectable levels of CIN are also important: the tumour genome must have at least 15 copy number changes to enable forecasting of tumour suppressor gene homozygous deletion, and tumours with less than 20 copy number changes are considered at low risk of acquiring an oncogene amplification by default. In our framework, we made an explicit decision to only include signatures that were robustly quantifiable across sequencing technologies (**Figure S34**). We additionally identified that the forecasting models show robust predictions across sequencing technologies (**Figure S17**). Despite the robustness of the proposed framework, we recommend profiling the tumour DNA using the same modality as the training cohort where possible. We also recommend to fit copy number profiles with a uniform software pipeline across the training cohort and the input samples.

*Genomics research setting:* Here, the user does not need to specify a tumour type or gene as input (although we recommend specifying the tumour type when possible). This allows full exploration of potential oncogene amplifications or tumour suppressor gene homozygous deletion that are likely to emerge in the future evolution of the tumour. If the user specifies the tumour type, only the *driver-specific mutational processes* ($\omega_{D,CX}$) and *driver-specific selection coefficients* ($s_D$) from that tumour type are used to calculate *P(D)*. An optimal threshold within that tumour type can then be applied to binarise *P(D)*. If the tumour type is not specified, forecasting is performed in pan-cancer mode by averaging the *P(D)* obtained for all tumour types in which the amplification or homozygous deletion of the target gene is putatively under positive selection (**Figures S1-S2**). In this case, a pan-cancer threshold can be estimated and then applied to binarise *P(D)*.

*Clinical research setting:* Here, in most instances, forecasting will be performed with a specific gene and tumour type in mind. Thus, it is recommended to specify this as input so that only the *driver-specific mutational processes* ($\omega_{D,CX}$) and *driver-specific selection coefficients* ($s_D$) for the given tumor type are used in the framework.

#### How do I set the optimal threshold?

Choosing an optimal threshold is highly dependent on the specific research question, tumour type, treatment status and disease stage. The threshold can be tuned such that the framework is conservative in its prediction of high-risk cases (maximising specificity), comprehensive (maximising sensitivity) or balanced (maximising a trade-off between sensitivity and specificity). In this study, we devised three different approaches for the automated derivation of thresholds tailored to achieve these desired outcomes:

1. Youden's index threshold: This strategy uses a reference cohort to determine a threshold that maximises prediction sensitivity and specificity via the Youden's index. A weighted version of Youden's index can also be applied if users prefer to emphasize either sensitivity or specificity.

2. Expected alteration frequency: This approach determines a threshold that matches prior knowledge on the fraction of samples expected to acquire an amplification, prioritizing prediction specificity.

3. Minimum *P(D)* in carriers: This strategy uses a training cohort to determine a matching the minimum *P(D)* value across all cases that already have the driver alteration, maximizing prediction sensitivity.

Users also have the flexibility to determine thresholds based on their preferred approaches, using the distribution of *P(D)* values provided for TCGA samples in this study or, preferably, in a more closely matched reference cohort. Below, we provide some broad guidelines on how to practically proceed with threshold estimation in different scenarios:

*Genomics research setting:* Here, the user may not have a strong idea of the prior probability of acquiring a driver alteration. Thus, they are likely to want a balance between specificity and sensitivity. In this instance, we recommend using the Youden's index threshold estimated from the training data (TCGA by default). This threshold will work best if the samples intended to be studied match the characteristics of the training cohorts (e.g. primary tumours). If they differ substantially (e.g. post treatment) then it may be more appropriate to follow the clinical research setting suggestions below.

*Clinical research setting:* In this setting, it is assumed that the user has some prior knowledge of the likelihood of acquiring the driver copy number alteration of interest and that the clinical research question requires a much more precise focus on either maximising sensitivity or specificity. Selecting the optimal threshold in this instance requires a training cohort with similar properties to the input tumour samples. The more precisely matched the cohort characteristics are, the better the model will perform. Two strategies for setting the optimal threshold can be adopted:

1. If the training cohort is extremely well matched to the clinical research setting, a threshold can be chosen to maximise the performance of the framework on the cohort (e.g. maximising sensitivity, specificity or both, potentially using the approaches presented above).

2. In the absence of a suitably matched reference cohort, one possibility the user may want to consider is compiling a training cohort prior to deploying the forecasting framework. For example, we opted for this approach when using an *in vitro* dataset of osimertinib resistant cell lines to determine the optimal threshold for forecasting *MET* amplification. If that's not possible, albeit non-ideal, it is possible to rely on prior knowledge on the expected frequency of the driver amplification or deletion to yield a potentially reasonable threshold. For example, after androgen deprivation therapy, 45-52% of prostate cancer patients are expected to develop resistance through *AR* amplification[19]. In this instance, the user should first apply the trained *driver-specific mutational processes* ($\omega_{D,CX}$) and *driver-specific selection coefficients* ($s_D$) to each sample in the reference cohort to obtain a distribution of *P(D)* values. A threshold can then be selected to classify ~45% of the training cohort (in our example, the TCGA PRAD cohort) as having high risk of acquiring *AR* amplification. In practice, this threshold corresponds to the quantile $Q_{1-x}$ of the *P(D)* distribution in the reference cohort, where *x* represents the expert- or literature-derived expected frequency of driver gene amplification or deletion in the forecasting scenario.

#### How do I learn the framework parameters?

The default parameters for the framework, including *driver-specific mutational processes* ($\omega_{D,CX}$) and *driver-specific selection coefficients* ($s_D$) were learnt from the TCGA cohort. Ultimately, the quality of the final predictions hinges on how well the clinico-pathological and biological characteristics of this cohort matches the patient population where application is intended. For most users, the default values for these parameters will be suitable and no further training of the framework will be required. However, if the selective pressures or mechanisms of driver gene amplification or deletion differ greatly between the query sample and these cohorts, the user may want to relearn these parameters using a more suitable training cohort. In general, we recommend training all the parameters of the model using a cohort of tumours with oncogene amplifications or tumour suppressor gene homozygous deletions at an evolutionary end-point with respect to the samples for which the forecasting model will be applied. For instance, if the model is used to forecast an amplification driving treatment resistance, a cohort of resistant samples that was subjected to the same treatment selection pressures is preferred. If the user has a suitable reference cohort available, the *driver-specific mutational processes* parameter can be estimated using the procedure described in the Methods' section "Estimating locus-specific mutational processes ($\omega_{D,CX}$)", and the *driver-specific selection coefficient* can be estimated using the procedure described in the Methods' section "Estimating selection coefficients (*s*)".

### Supplementary Note 3 - On the feasibility of forecasting oncogene amplifications

According to classical tumour evolutionary theory, mutation and selection (plus genetic drift) determine how likely an oncogene is to be amplified in the future[1]. In simple terms, individual cells within a tumour acquire new amplifications via various DNA damage mechanisms, generating cell-to-cell karyotypic variation. Selection then favours the expansion of cells with an amplification that confers a fitness advantage.

Single-cell whole genome DNA sequencing data offers an opportunity to observe this process in action. Amplifications observed exclusively in single cells provide a detailed view of the process of karyotypic diversification in a tumour (**Supplementary Note Figure 2a**). Meanwhile, amplifications detectable in bulk (or pseudo-bulk) DNA sequencing data indicate those that are present in multiple cells, which means that they were present in a cell that underwent a clonal expansion in the past, potentially driven by the fitness advantage conferred by the amplification itself (**Supplementary Note Figure 2a**).

Here, we compared the number of clonally-expanded and unique amplifications in 16,178 single cells from 8 human triple-negative breast cancers (TNBC) and 4 cell lines[2]. We observed unique amplifications generated during recent karyotypic diversification in 11 out of 12 samples (**Supplementary Note Figure 2b**) and clonally-expanded amplifications in 7 out of these 11 samples. This suggests that amplifications are actively generated in the evolution of most TNBC samples. However, the rate at which new amplifications emerged varied across TNBC samples, ranging from 2.7 unique amplifications per 1,000 cells in sample TN3 to 13.8 unique amplifications per 1,000 cells in TN4. This indicates significant variation in the ongoing activity of amplification-generating mechanisms across TNBC samples. Notably, one sample showed neither unique nor clonally-expanded amplifications (**Supplementary Note Figure 2b**), likely reflecting inoperative mechanisms causing amplifications throughout its evolutionary history. Overall, these findings highlight that the acquisition of amplifications is an ongoing consequence of karyotypic diversification in most TNBCs, occurring at different rates.

Selection ultimately determines whether these amplifications expand to dominate the tumour population. Notably, we observed that 4 samples did not show any clonally-expanded amplification despite showing a moderate number of unique amplifications (mean of 7.15 and range from 2.72 to 9.93 unique amplifications per 1,000 cells) (**Supplementary Note Figure 2b**). If these mechanisms of amplification were active during the previous evolution of the tumour, they must have generated amplifications during karyotypic diversification. Their absence at the bulk level suggests that, in such a scenario, the generated amplifications did not confer a positive advantage or were negatively selected.

To formally assess the extent to which selection drives the expansion of amplifications generated during karyotypic diversification, we designed a positive selection test for amplifications. We reasoned that positively selected amplifications would predominantly amplify oncogenes, contrary to passenger amplifications. Applying this test to unique amplifications, we found no enrichment of breast cancer oncogenes (**Supplementary Note Figure 2c**). In contrast, clonal amplifications were significantly enriched in breast cancer oncogenes (**Supplementary Note Figure 2c**). This underscores the role of selection in constraining the initial pool of amplifications generated during karyotypic diversification, resulting in a landscape of clonally-expanded amplifications enriched in oncogenes.

The absence of detectable positive selection in unique amplifications enabled us to exclude selection as a confounder when studying the genome-wide activity of mechanisms of amplification. We reasoned that if these mechanisms were to operate uniformly across the genome, then we would see an even distribution of unique amplifications across a chromosome (i.e. nearest unique amplifications would be at a distance equal to the length of the chromosome divided by the total number of unique amplifications observed in the chromosome). However, across all 18 samples we observed that unique amplifications clustered together, with distances between adjacent unique amplifications being significantly shorter than expected (**Supplementary Note Figure 2d**). This suggests that amplification rate varies across genomic regions, potentially depending on the active mechanisms of amplification within a sample.

To understand how these evolutionary factors are intertwined in determining how likely a *specific* oncogene amplification is to arise, we simulated tumour evolution using different selection coefficients for the oncogene amplification and rates at which it is acquired in the tumour population (**Supplementary Note Figure 2e**). The latter depends both on the activity of the mechanisms of amplification and how frequently these mechanisms operate on the locus harbouring the oncogene. Our simulations revealed that the varying rates of amplification acquisition observed during karyotypic diversification in TNBCs could significantly affect the likelihood of a new amplification emerging and clonally expanding (**Supplementary Note Figure 2f**). Selection also had a notable effect, with high selection coefficients promoting the clonal expansion of amplifications even when their rate of acquisition was lower (**Supplementary Note Figure 2f**). These findings emphasise the need to jointly model the varied selective coefficients and amplification rates to accurately forecast oncogene amplification.

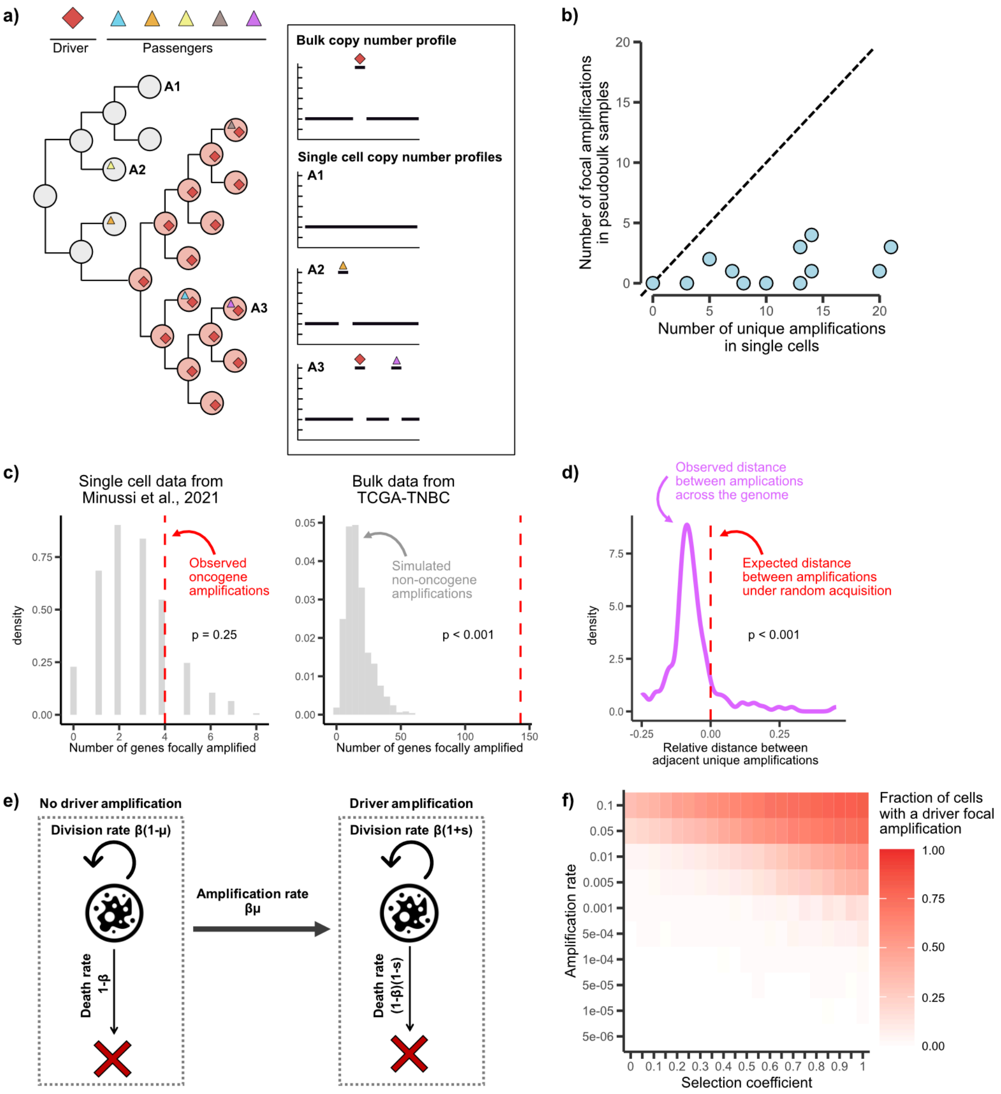


**Supplementary Note Figure 2. Acquisition, selection and expansion of focal amplifications in triple-negative breast cancer. a)** Schematic showing the ability to identify unique and clonally expanded focal amplifications in single-cell and bulk DNA sequencing data. **b)** Scatter plot comparing the number of unique focal amplifications detected across all single cells with the number of clonally expanded focal amplifications seen at pseudobulk resolution in 8 triple-negative breast cancer (TNBC) samples and 4 TNBC cell lines. **c)** Density plots describing the expected number of gene amplifications in single cells (left) and in the TCGA-TNBC cohort (right) based on 1,000 iterations with lists of random genes. The red dashed line highlights the observed number of oncogenes amplified. Statistical significance was measured by calculating the empirical p-value. **d)** Density plot showing the relative observed distance between adjacent unique amplifications observed across 16,178 cells from 12 TNBC samples. The red dashed line highlights the relative expected distance between adjacent amplifications under random acquisition. P-value is estimated by one-sample Wilcoxon's two-sided test. **e)** Schematics of birth-death simulations of tumour growth (fitness advantage: s, death rate: α, birth rate: β, amplification rate: μ). Tumour cells without amplification (left box) die with rate α, divide yielding two cells without amplification with rate β(1-μ) and divide giving rise to one cell with amplification and another without with rate βμ. Tumour cells with amplifications (right box) have a fitness advantage and as such die with rate α(1-s) and divide with rate β(1+s), giving rise to two cells with amplification. **f)** Mean fraction of cells harbouring an oncogene amplification at the end of 20 replicate birth-death tumour growth simulations using different combinations of selection coefficient (x-axis) and amplification rate (y-axis).

### Supplementary Note 4 - Particularities of tumour suppressor genes homozygous deletions

The loss of a tumour suppressor gene (TSG) represents a key event in the development of many cancers. Much like the forecasting of oncogene amplification, our framework can be adapted to predict homozygous TSG loss. However, the nature of these events has some important differences compared to oncogenes.

First, these events can occur early in tumour evolution, where evidence of CIN operating in the genome is sparse. Indeed, in the TCGA cohort, 13% of samples without detectable CIN had evidence of homozygous TSG loss. This is in contrast to oncogene amplification where 1.9% of samples without detectable CIN showed evidence of these events (**Figure S35)**.

Second, TSG loss typically occurs via two independent genomic hits[20], where each copy of the gene is lost at different stages during tumour evolution. To understand the nature of each hit, we compared single copy number loss events spanning TSGs to double hits. We leveraged our 3-feature copy number encoding to compare the size, changepoint and breakpoint density in segments harbouring TSGs across samples with hemizygous versus homozygous deletions. To reduce the impact of noise from CIN processes arising later in tumour evolution, we initially examined diploid samples (ploidy < 2.7). In diploid samples, hemizygous deletions were defined by long-sized segments with 1-copy changes; while homozygous deletions spanned short-sized segments with 1-copy change (**Figure S15**). These differences suggest that single and double copy losses may be driven by different mechanisms generating long vs focal losses, with focal homozygous deletions potentially occurring following an initial whole or arm-level chromosome loss. We observed similar configurations of single and double losses in tetraploid samples (ploidy ≥ 2.7), except for higher changepoints in homozygous deletions where 1-to-3 copy changes appear (**Figure S15**). These findings support the hypothesis that homozygous deletions are early events, followed by subsequent mutational processes, such as whole-genome duplication (WGD) that would amplify adjacent segments. Indeed segments flanking homozygous deletions tend to show a copy number of 1 in diploid tumours, but 2 or higher in tetraploid tumours (**Figure S16**). Thus when training our model, we need to adjust the changepoint feature values to account for subsequent WGD events. In addition, we need to adjust our risk predictions based on the current number of copies in a tumour as higher copy number states will result in a lower risk score.

Third, homozygous genome loss is commonly deleterious and thus heavily negatively selected. Beyond tumour suppressor genes, the most frequent homozygous deletions are found in the T-cell receptor alpha locus, immunoglobulin heavy and light chain loci, and fragile sites[20]. In fact, approximately 70% of all homozygous deletions that did not involve TSGs were located in these regions. It is well known that the mechanisms driving the acquisition of homozygous deletions in these regions are completely different from those impacting on TSGs, indicating that the evidence of CIN-related passenger homozygous deletions in the genome is limited. Therefore, it is not possible for us to account for local topology constraints in this instance.

### Supplementary Note 5 - Exploring the mutational processes putatively causing oncogene amplifications and tumour suppressor deletions

The "locus-specific mutational processes" matrix shows important etiological relationships between putative causal signatures and particular oncogene amplifications/tumour suppressor deletions (**Supplementary Note Figure 3**).

CX13, representing a form of replication stress linked with circular ecDNA formation[3] (**Figure S36**) known to be a major source of DNA amplification[4], was the most commonly enriched putative causal signature across oncogenes (**Supplementary Note Figure 3a**). CX8 and CX9, other replication stress signatures, were also enriched across oncogenes. A subset of oncogenes had CX1 enriched, a signature associated with fold-back inversion[3], which is another mechanism known to generate amplifications[5]. In the case of homozygous deletions (**Supplementary Note Figure 3b**), CX2 and CX3, two signatures linked to impaired homologous recombination[3] and extensive deletions across the genome, were enriched in the majority of tumour suppressor genes.

Interestingly, we observed both shared and variable patterns of signature enrichment across tumour types for specific oncogenes (**Figure S31**) and tumour suppressor genes (**Figure S33**). For example, most tumours with *EGFR* amplification showed an enrichment of CX13; however, lung adenocarcinomas showed a distinct enrichment of CX8, a signature putatively linked with R-loop accumulation[3]. A similar trend was observed for *PTEN* deletions, which were typically enriched for CX2 except for sarcomas, where CX1 enrichment suggests missegregation errors as the putative mechanism.

We also observed shared and variable enrichment patterns across genes within a tumour type (**Figures S30 and S32**). For instance, in prostate cancer, amplification of *AR* and *NCOA2* were enriched for CX8. As the amplification of both oncogenes is linked to resistance to ADT[6,7], the mechanisms underpinning CX8 may enable multiple paths to resistance. In contrast, amplification of *MYC* in prostate cancer, also known to modulate AR signalling[8], was enriched for CX9, suggesting multiple DNA damage mechanisms can drive resistance in the disease. Similarly, in colorectal cancer, we observed different signatures causing the loss of *SMAD3* and *SMAD4*, two members of the TGF-β signalling pathway that normally act as tumour suppressors in early colorectal cancer development[9].

Altogether, our results show that the "locus-specific mutational processes" matrix ($\omega_{D,t,CX}$) may be useful for exploring amplification and deletion aetiology independent of their use in the forecasting framework.

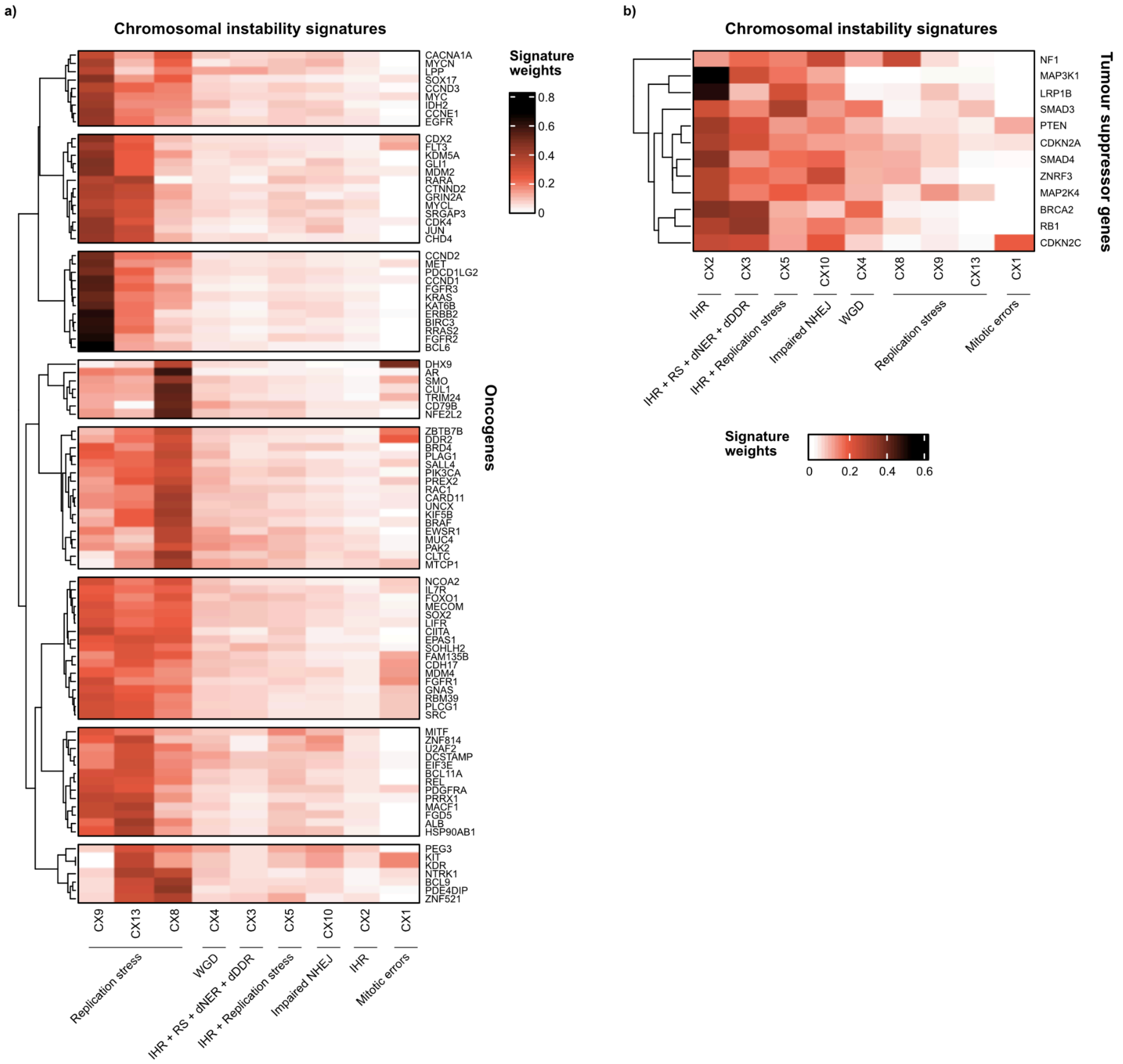


**Supplementary Note Figure 3. Types of chromosomal instability underpinning oncogene amplifications and tumour suppressor gene deletions.** Heatmaps highlighting the enrichment of CIN signatures (signature weights) in **a)** amplified segments harbouring a specific oncogene across TCGA, and **b)** homozygous deleted segments harbouring a specific tumour suppressor gene. Signature weights were learnt from the TCGA cohort by computing the ratio of the driver-specific signature score (i.e. the average of the signatures assigned to each altered copy number event harbouring the specific driver gene) to the background signature score (i.e. the average of the signatures assigned to all copy number segments). Signature aetiologies are stated at the bottom. IHR, impaired homologous recombination; RS, replication stress; dNER, nucleotide excision repair deficiency; dDDR, DNA damage repair deficiency; WGD, whole-genome duplication; NHEJ, non-homologous end joining. Here, we focused on the 9 CIN signatures which were robustly quantifiable across sequencing platforms to enable forecasting from multiple DNA sequencing assays (see Methods).

### Supplementary Figures

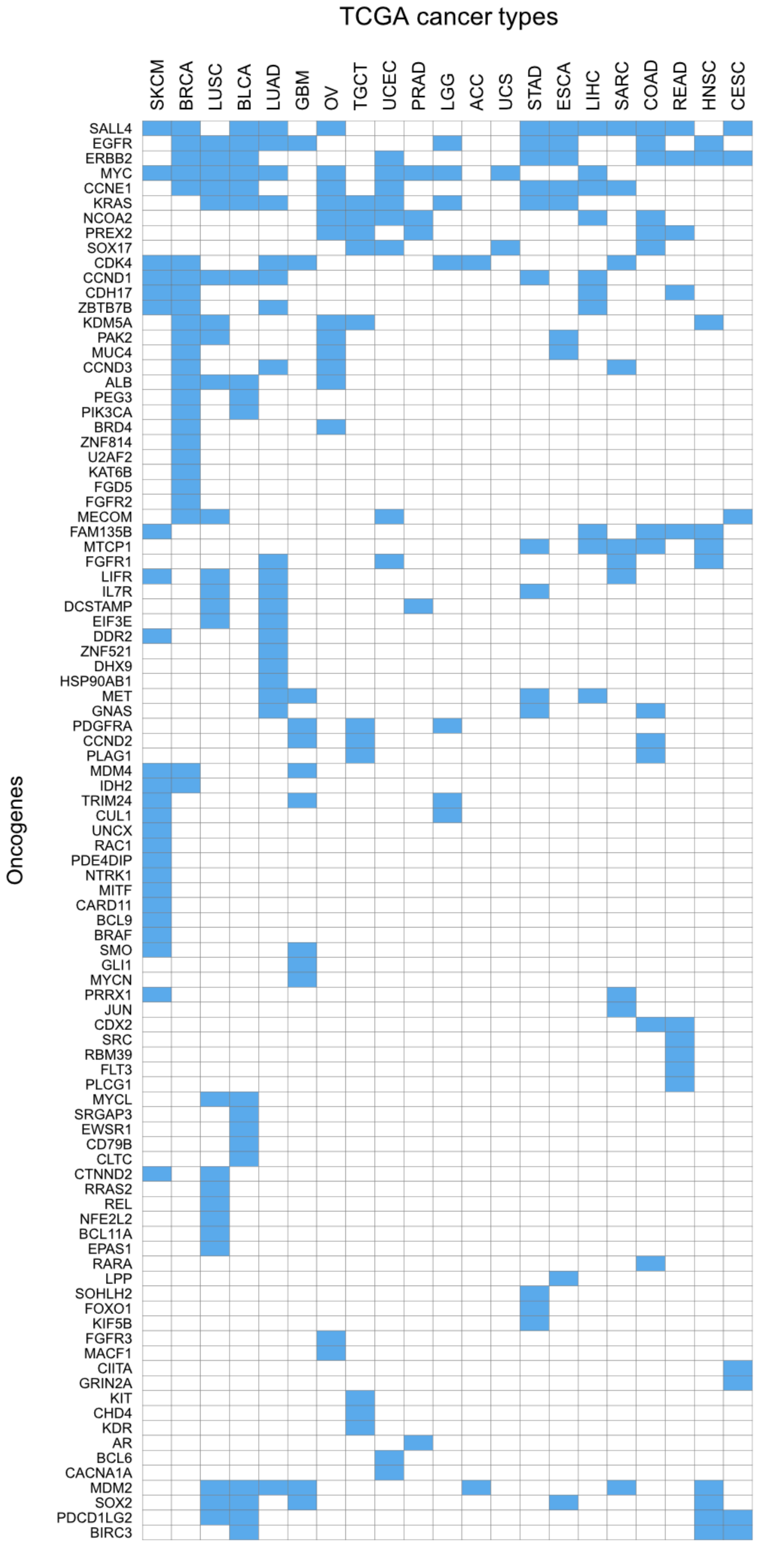


**Figure S1. Summary of the different oncogenes positively selected to be amplified per tumour type.** An oncogene was considered a putatively amplified oncogene in a tumour type when it was amplified in ≥ 5 samples in the TCGA tumour-specific cohort, and it was previously identified as significantly amplified in the given tumour type by GISTIC.

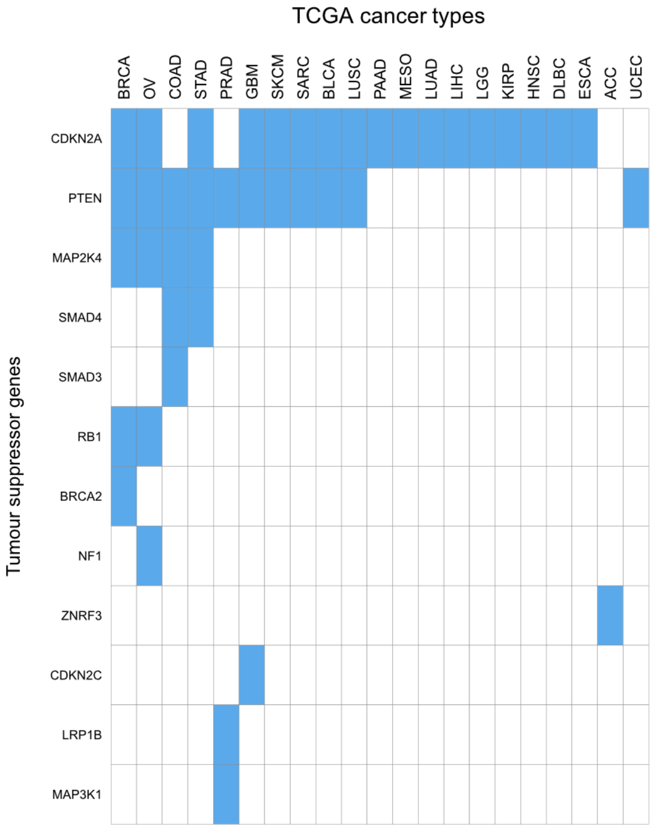


**Figure S2. Summary of the different tumour suppressor genes positively selected to be deleted per tumour type.** A tumour suppressor gene (TSG) was considered a putatively deleted TSG in a tumour type when it was homozygous deleted in ≥ 5 samples in the TCGA tumour-specific cohort, and it was previously identified as significantly deleted in the given tumour type by GISTIC.

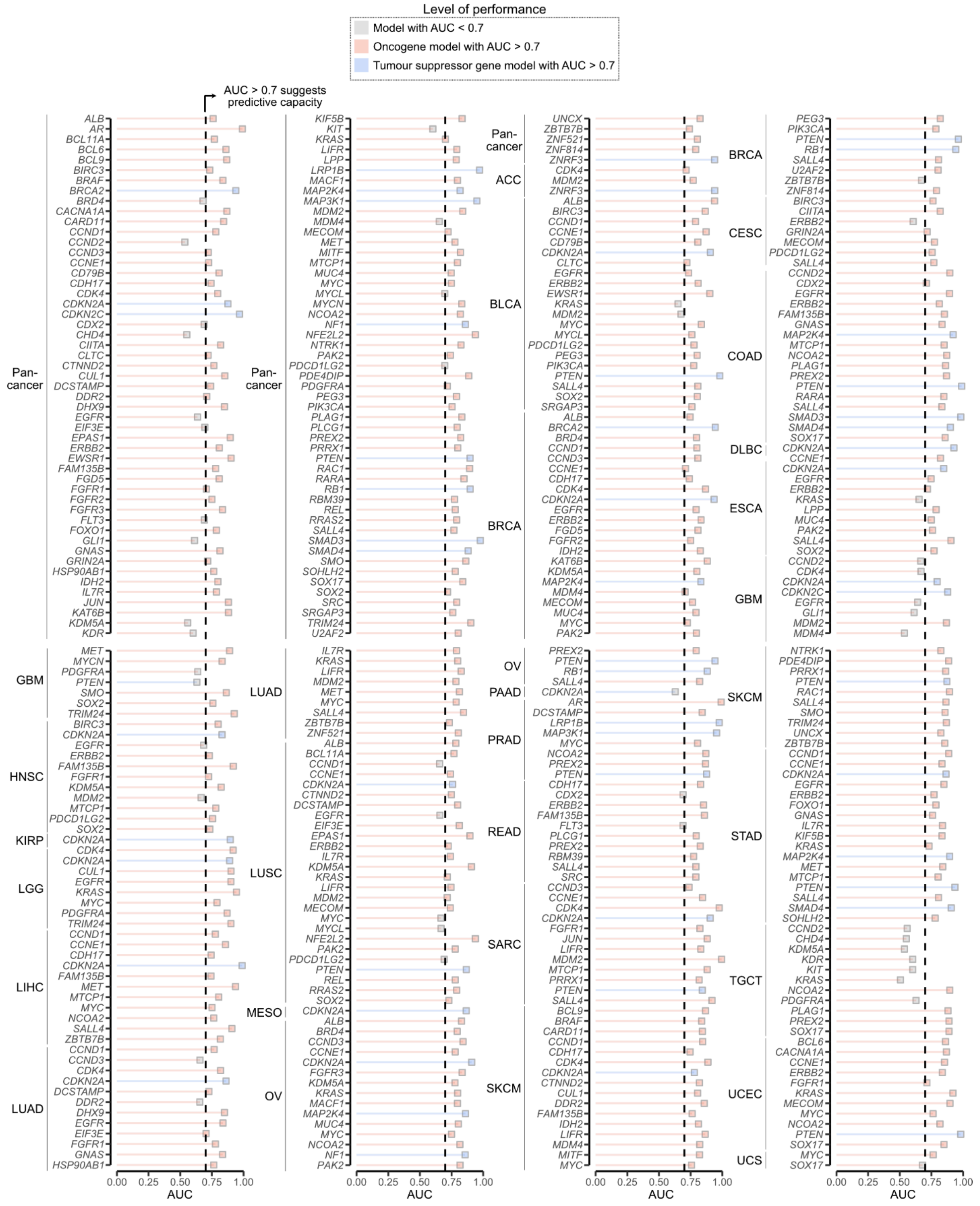


**Figure S3. Evaluation of the framework's predictive performance in the TCGA cohort.** Area under the receiving operating curve (AUC) for 300 tumour-type and 107 pan-cancer forecasting models reapplied in the The Cancer Genome Atlas (TCGA) dataset. Models were trained using the TCGA dataset. Dashed line indicates AUC value of 0.7, the threshold for suggestive predictive potential. Red colour denotes models for oncogenes with AUC > 0.7, blue colour highlights models for tumour suppressor genes with AUC > 0.7, and grey colour indicates models with AUC < 0.7. Pan, pan-cancer; ACC, adrenocortical carcinoma; BLCA, bladder urothelial carcinoma; BRCA, breast cancer; CESC, cervical squamous cell carcinoma and endocervical adenocarcinoma; COAD, colon adenocarcinoma; DLBC, diffuse large B-cell lymphoma; ESCA, esophageal carcinoma; GBM, glioblastoma multiforme; HNSC, head and neck squamous cell carcinoma; KIRP, kidney renal papillary cell carcinoma; LGG, low grade glioma; LIHC, liver hepatocellular carcinoma; LUAD, lung adenocarcinoma; LUSC, lung squamous cell carcinoma; MESO, mesothelioma; OV, ovarian serous cystadenocarcinoma; PAAD, pancreatic adenocarcinoma; PRAD, prostate adenocarcinoma; READ, rectum adenocarcinoma; SARC, sarcoma; SKCM, skin cutaneous melanoma; STAD, stomach adenocarcinoma; TGCT, testicular germ cell tumors; UCEC, uterine corpus endometrial carcinoma; UCS, uterine carcinosarcoma. Performance was only evaluated for models with at least 5 samples with the driver copy number alteration.

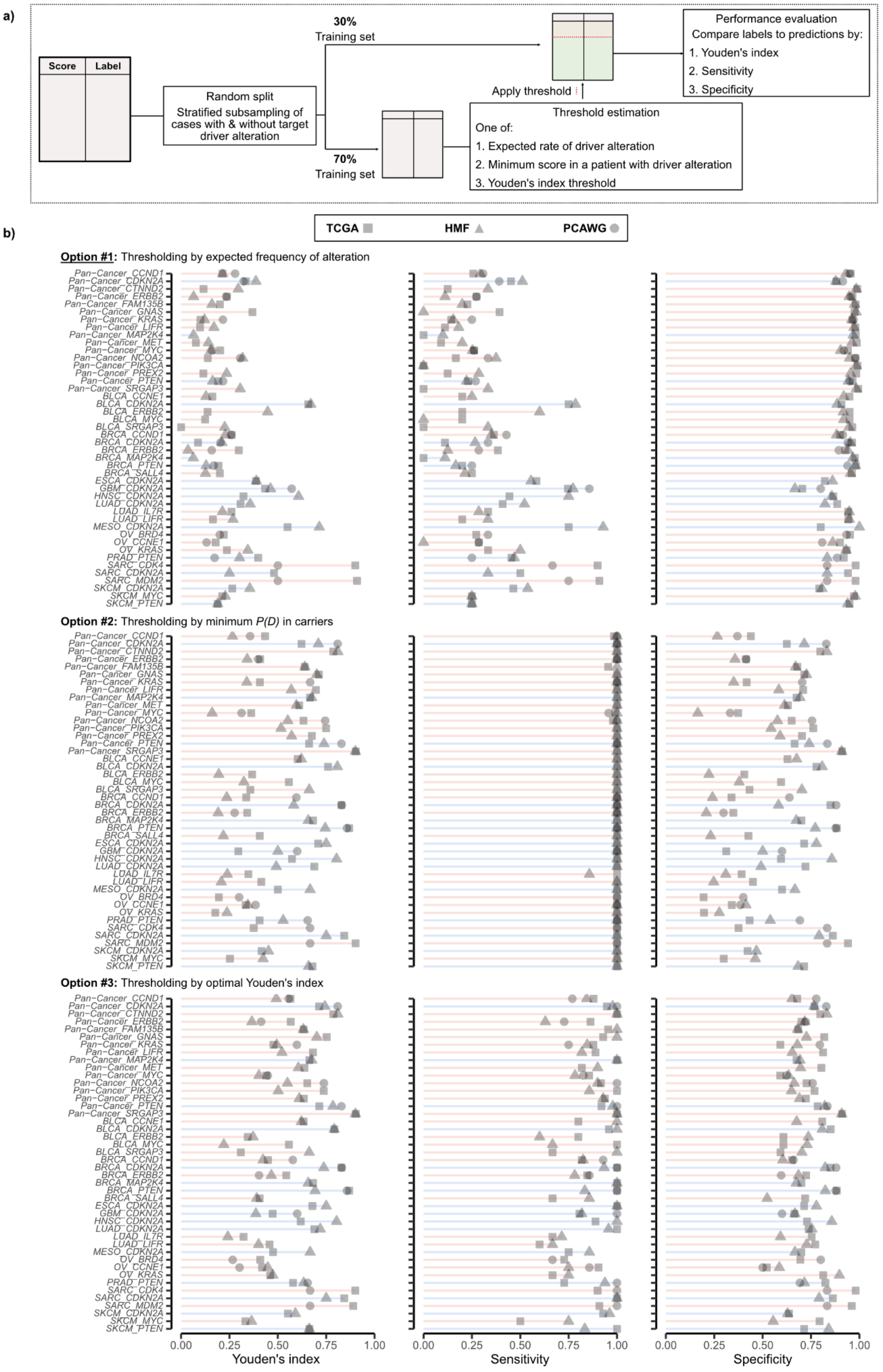


**Figure S4. Evaluating the performance of risk classifications in TCGA, PCAWG and HMF using a threshold derived in an independent cohort-specific subset. a)** Schematic representation of the procedure to split the cohort in a training and a testing subset, derive the threshold in the training subset, and ultimately apply the derived threshold to classify samples in the testing subset. Threshold was derived using three different approaches. This procedure was repeated 100 times, with the average values presented in panel **b**. The forecasting models were trained in the TCGA and applied in the different cohorts, while the threshold to binarise predictions are optimised in the training subset in each cohort. This emulates a scenario in which a researcher has a reference cohort with similar clinico-pathological and biological characteristics to the cohort where the model will be applied. See Methods and Supplementary Note 4 for additional details. **b)** Youden's index (1 - sensitivity + specificity, left panel), sensitivity (middle panel) and specificity (right panel) for the predictions binarised following the procedure described in **a**. The probabilities of driver acquisition *P(D)* for each sample were derived by applying the forecasting models to copy number profiles in which the driver copy number alteration and its adjacent segments were removed to avoid target leaking. In the top row, the threshold was set such that the proportion of high-risk samples in the reference cohort matched the expected frequency of driver alterations. In the middle row, the threshold was defined as the minimum *P(D)* value observed in samples harbouring the driver alteration. In the bottom row, the threshold was set to maximize the Youden's index in the training subset. Red lines indicate models for oncogenes, blue lines indicate models for tumour suppressor genes. Forecasting performance was only evaluated for models with evidence of predictive performance (AUC > 0.7, Figure 2, Table S1) and in tumour-specific TCGA (squares), HMF (triangles) or PCAWG (circles) cohorts with at least 10 samples harbouring the driver alteration.

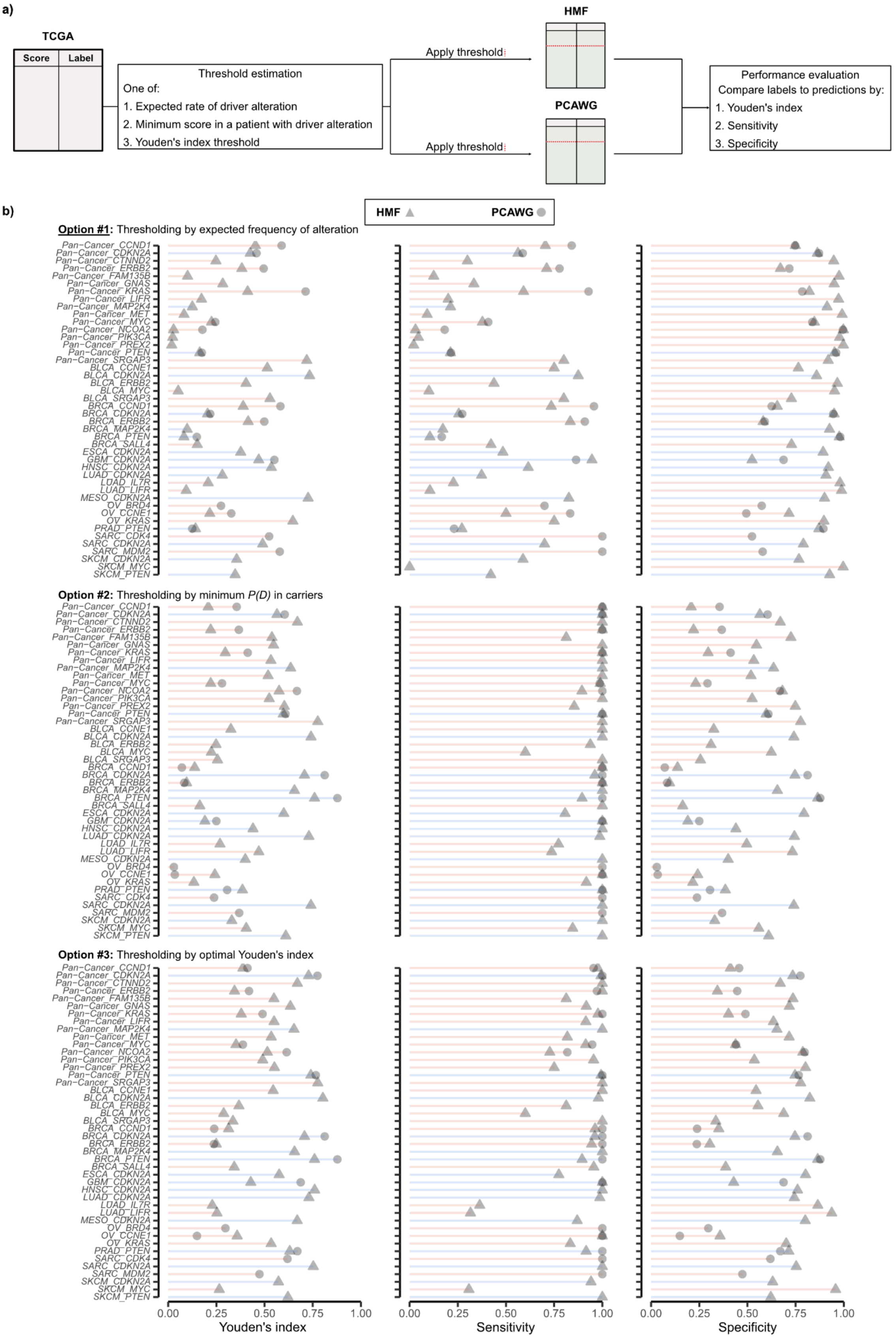


**Figure S5. Evaluating the performance of risk classifications in PCAWG and HMF using a threshold derived in the TCGA cohort. a)** Schematic representation of the procedure to derive a threshold in the TCGA cohort, and ultimately apply the derived threshold to classify samples in the

PCAWG and HMF cohorts. Threshold was derived using three different approaches. The forecasting models and the threshold for binarising predictions were trained in the TCGA and applied in the different cohorts. This emulates a scenario in which a researcher does not have a reference cohort with similar clinico-pathological and biological characteristics to the cohort where the model will be applied. See Methods and Supplementary Note 4 for additional details. **b)** Youden's index (1 - sensitivity + specificity, left panel), sensitivity (middle panel) and specificity (right panel) for the predictions binarised following the procedure described in **a**. The probabilities of driver acquisition *P(D)* for each sample were derived by applying the forecasting models to copy number profiles in which the driver copy number alteration and its adjacent segments were removed to avoid target leaking. In the top row, the threshold was set such that the proportion of high-risk samples in the reference cohort matched the expected frequency of driver alterations. In the middle row, the threshold was defined as the minimum *P(D)* value observed in samples harbouring the driver alteration. In the bottom row, the threshold was set to maximize the Youden's index in the training subset. Red lines indicate models for oncogenes, blue lines indicate models for tumour suppressor genes. Forecasting performance was only evaluated for models with evidence of predictive performance (AUC > 0.7, Figure 2, Table S1) and in pan-cancer/tumour-specific TCGA (squares), HMF (triangles) or PCAWG (circles) cohorts with at least 10 samples harbouring the driver alteration.

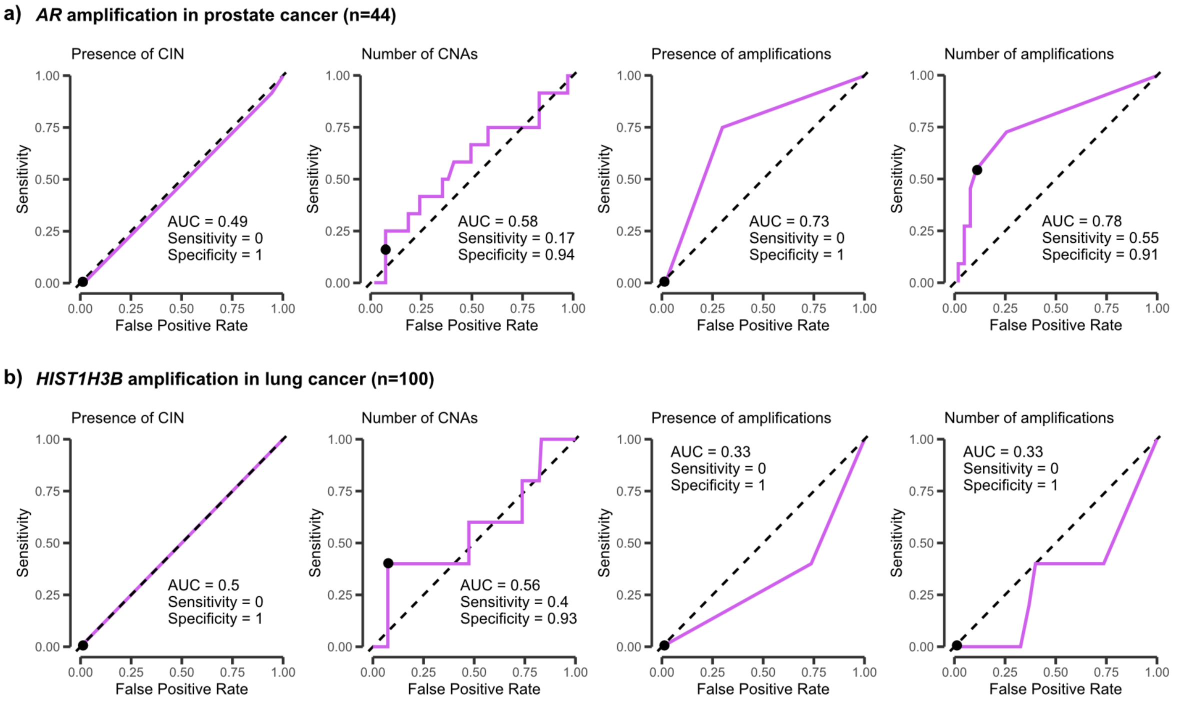


**Figure S6. Forecasting performance of four genomic features across longitudinal data. a-b)** ROC curves showing performance of four genomic features (presence of CIN, number of CNAs, presence of amplifications, and number of amplifications) for forecasting the acquisition of **a)** *AR* amplification in postreated prostate cancer samples, and **b)** *HIST1H3B* amplification in metastatic lung cancer samples. The predictions were performed in the early time point sample. The black dot highlights the effectiveness of the predictions by applying the optimal threshold derived based on the expected frequency of positive samples.

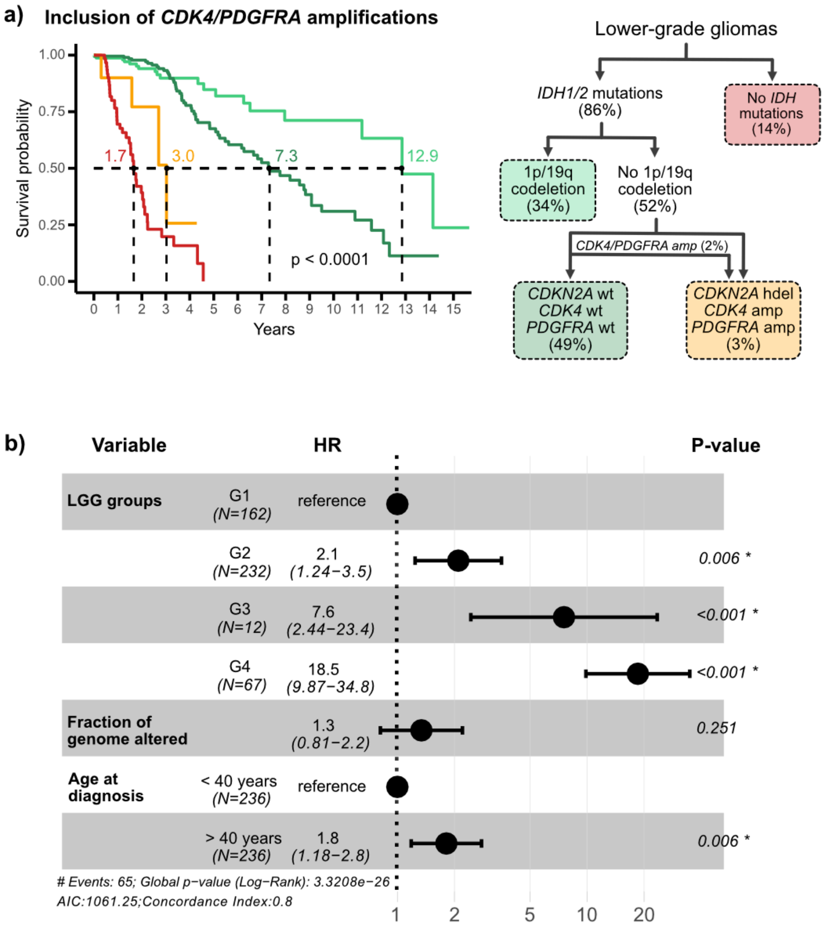


**Figure S7. Survival analysis in low-grade gliomas patients from the TCGA and the GLASS cohorts classified based on the inclusion of *CDK4/PDGFRA* amplification status in the diagnosis algorithm for classifying IDH-mutant-non-codeleted cases. a)** Kaplan-Meier curves showing survival times for the different LGG groups. Patients are classified based on the 2021 World Health Organization Classification of Tumours of the Central Nervous System, with IDH-mutant-non-codeleted cases being additionally classified based on the *CDK4/PDGFRA* amplification status. Lines are coloured according to the groups from the diagram in the right. Median survival values are indicated for each group. The p-value was estimated by the log-rank test. The prevalence of each low-grade glioma subtype is shown in brackets. **b)** Cox proportional hazards model using the classification algorithm described in a). The model was corrected for the fraction of genome altered and age at diagnosis. Age at diagnosis was categorised into two groups (<40 versus ≥40 years) based on clinical risk. G1, IDH-mutant-codeleted; G2, IDH-mutant-non-codeleted; G3, IDH-mutant-non-codeleted with *CDKN2A* homozygous deletion and/or *CDK4/PDGFRA* amplification; G4, IDH wild-type.

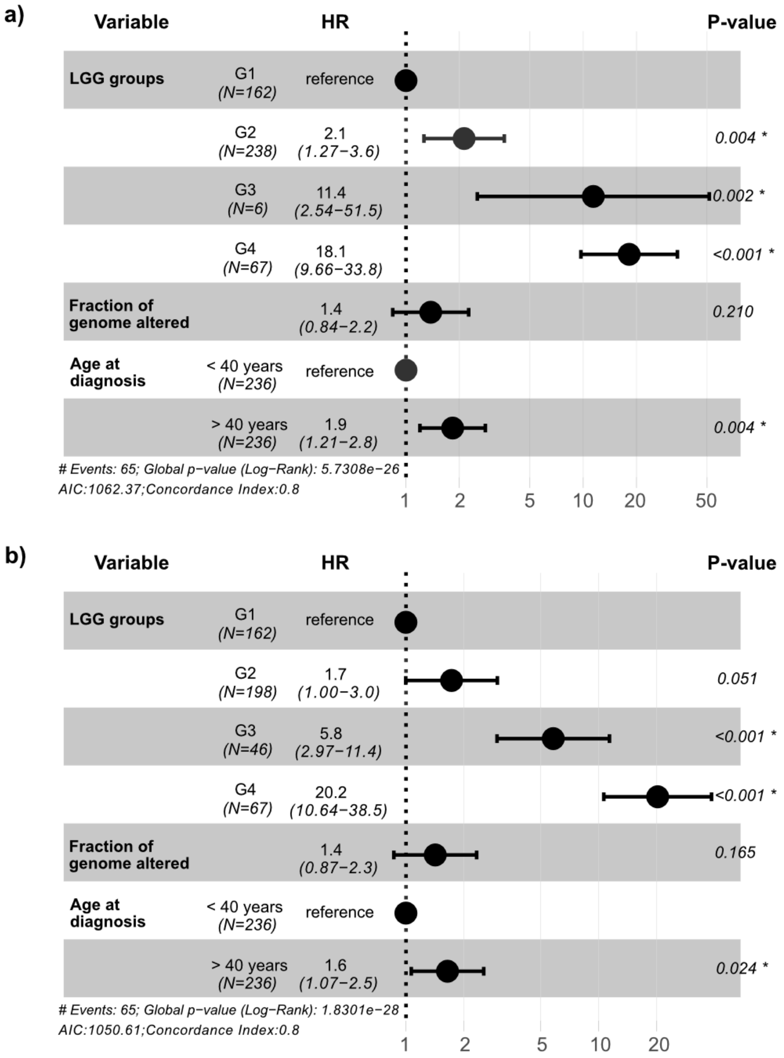


**Figure S8. Results of a Cox proportional hazards model for TCGA-LGG and GLASS patients based on the 2021 WHO diagnosis algorithm and the proposed extended classification.** Cox proportional hazards models were corrected for the fraction of genome altered and age at diagnosis. Age at diagnosis was categorised into two groups (<40 versus ≥40 years) based on clinical risk. **a)** Patients were classified based on the 2021 WHO diagnosis algorithm (as described in Figure 3a): G1, IDH-mutant-codeleted; G2, IDH-mutant-non-codeleted; G3, IDH-mutant-non-codeleted with *CDKN2A* homozygous deletion; G4, IDH wild-type. **b)** Patients were classified based on the inclusion of the actual and predicted *CDK4/PDGFRA* amplification status and predicted *CDKN2A* deletion status in the diagnosis algorithm for classifying IDH-mutant-non-codeleted cases (as described in Figure 3b): G1, IDH-mutant-codeleted; G2, IDH-mutant-non-codeleted; G3, IDH-mutant-non-codeleted with or with high-risk for *CDKN2A* homozygous deletion and/or *CDK4/PDGFRA* amplification; G4, IDH wild-type.

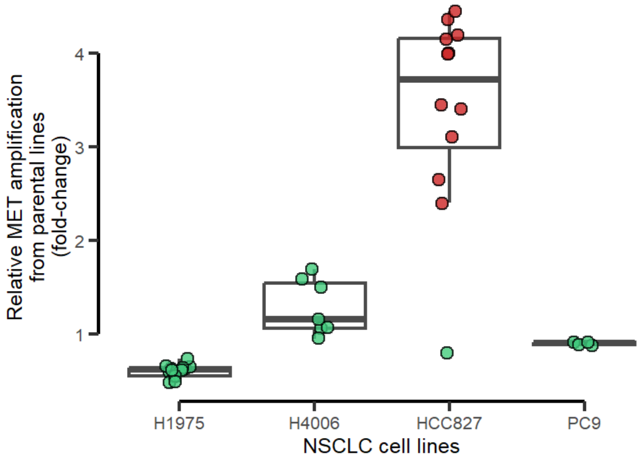


**Figure S9. Acquisition of *MET* amplification in EGFR-TKI resistant clones.** Resistant clones were derived from 4 NSCLC cell lines after long-term culture with EGFR-TKI. Red colour denotes clones acquiring *MET* amplification as the resistant mechanism, while green colour highlights clones not acquiring *MET* amplification. Droplet digital PCR was used to detect *MET* amplification, and a 2-fold increase from the parental cell line was considered as acquiring a *MET* amplification.

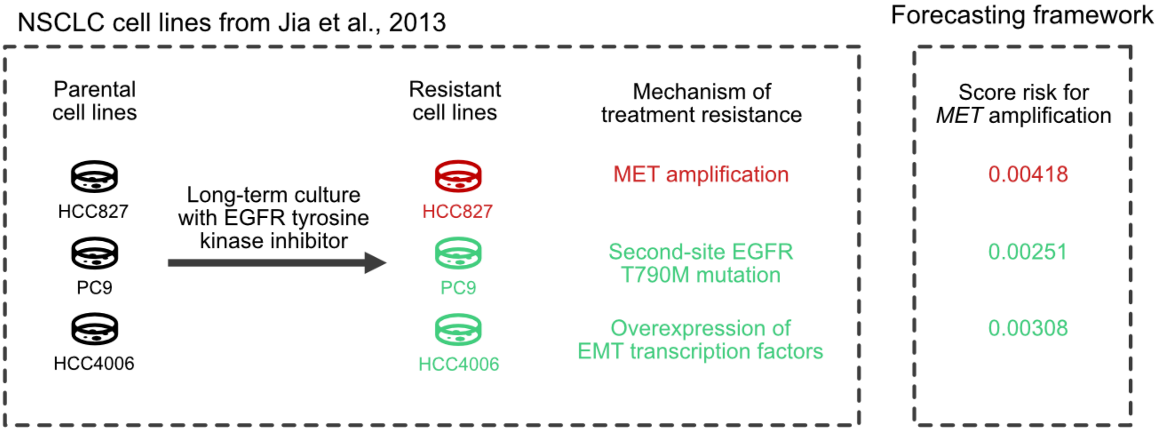


**Figure S10. Forecasting *MET* amplification-driven resistance to erlotinib in lung cancer cell line cultures.** Experimental validation of predicting *MET* amplification using a panel of NSCLC cell lines treated with erlotinib until the emergence of treatment resistance [21]. Scores of *MET* amplification risk were computed in parental cell lines, and the mechanism of treatment resistance acquired in resistant cultures was evaluated using different experimental techniques as previously reported [21]. Red colour indicates resistance driven by *MET* amplification.

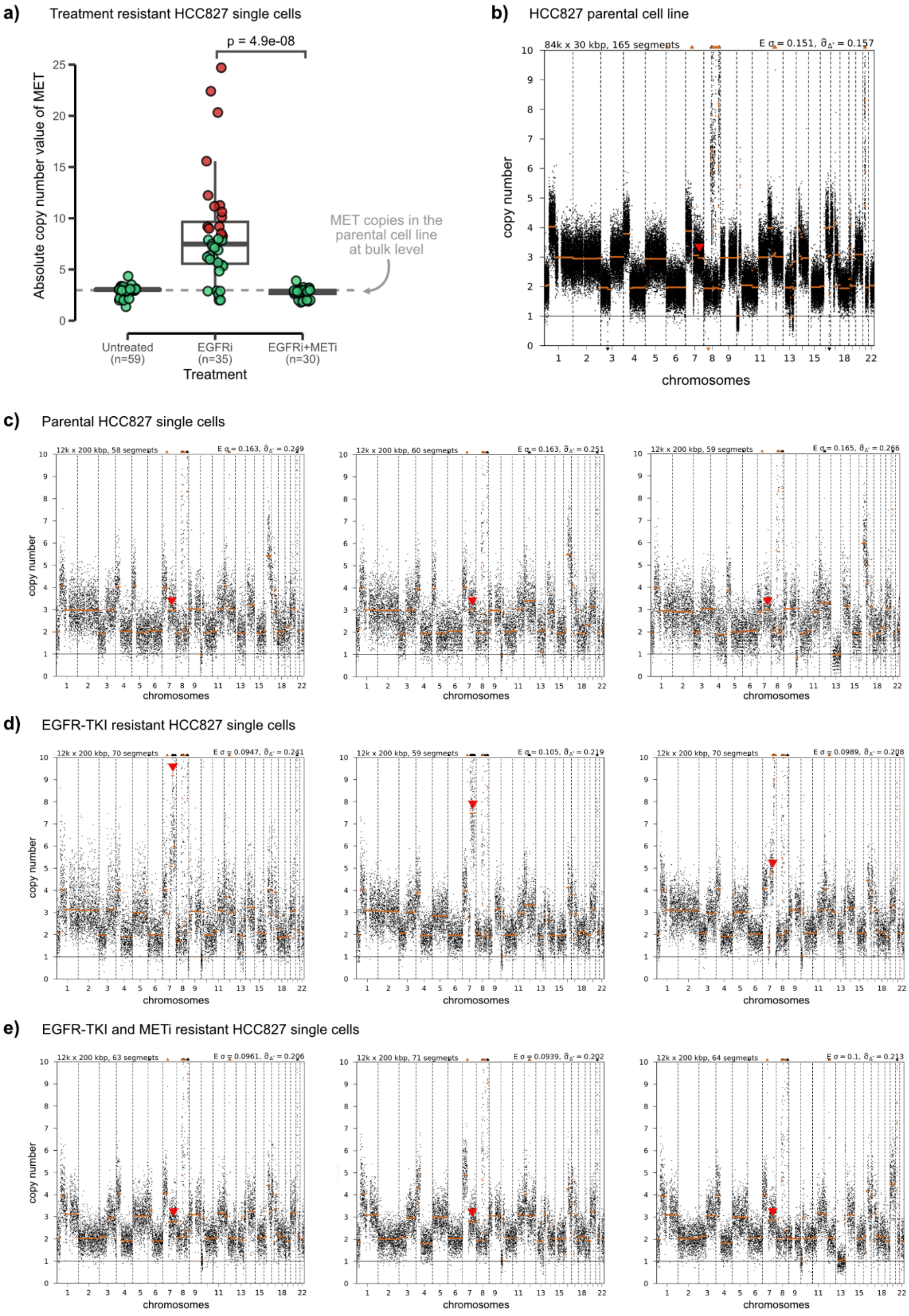


**Figure S11. Single-cell DNA sequencing of treatment-resistant HCC827 cells. a)** Boxplot showing the absolute copy number value of *MET* in HCC827 single cells after 1-month treatment with EGFR-TKI or EGFR-TKI and METi. The box bounds the interquartile range divided by the median. Red colour denotes cells with at least 8 copies of *MET*, while green colour denotes cells without focal amplification of *MET*. Horizontal dashed line indicates the absolute copy number value of *MET* in the parental cell line. P-value was estimated by Wilcoxon two-sided rank test. Untreated cells are used as control. **b)** Copy number profile of the HCC827 parental cell line. **c)** Copy number profiles of three untreated HCC827 single cells. **d)** Copy number profiles of three HCC827 single cells under EGFR-TKI treatment. **e)** Copy number profiles of three HCC827 single cells under EGFR-TKI and METi treatment. The segment mapping the genomic location of *MET* (7q31) is highlighted with a red triangle. Orange lines indicate copy number segments, while black dots indicate bins. Bulk copy number profile is derived at 30 kb bin resolution, while single-cell copy number profiles are derived at 200 kb bin resolution.

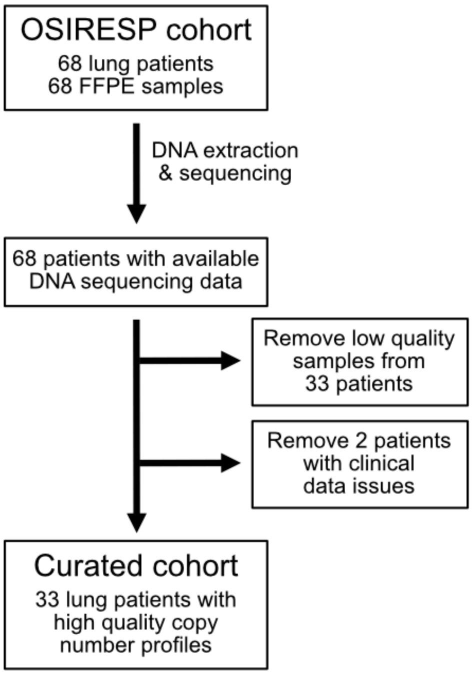


**Figure S12. REMARK diagram.** Flow diagram summarising the quality control filtering of patients to get the curated cohort for forecasting the acquisition of *MET* amplification as a resistance to osimertinib treatment.

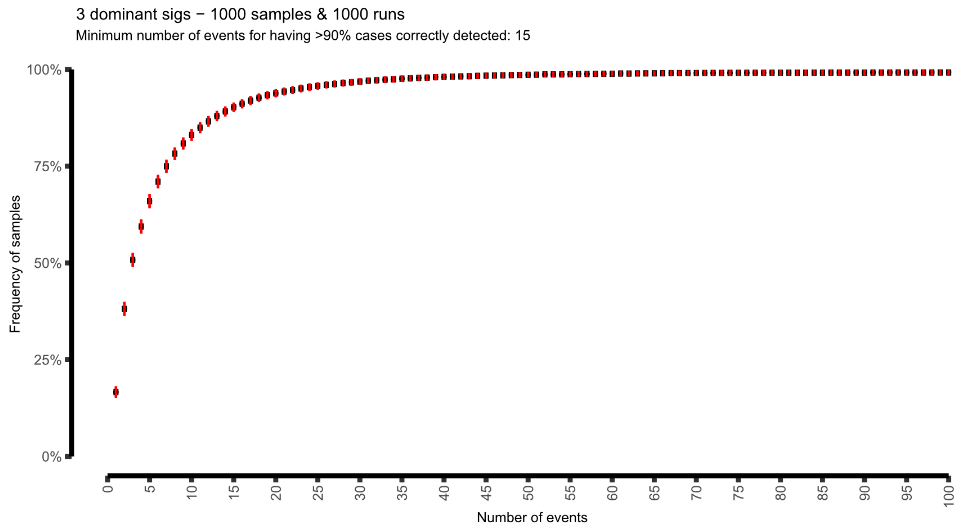


**Figure S13. Frequency of samples with correct signature decomposition based on the number of events retained in the copy number profile.** Plot shows the success rate of capturing the sample's signature composition (defined by its three dominant signatures) by progressively increasing the number of copy number events from 1 to 100. This procedure was repeated 1000 times with 1000 TCGA samples and copy number events used being randomly selected for each run. Black horizontal lines represent the mean across iterations, while red vertical lines indicate the standard deviation of the mean across iterations.

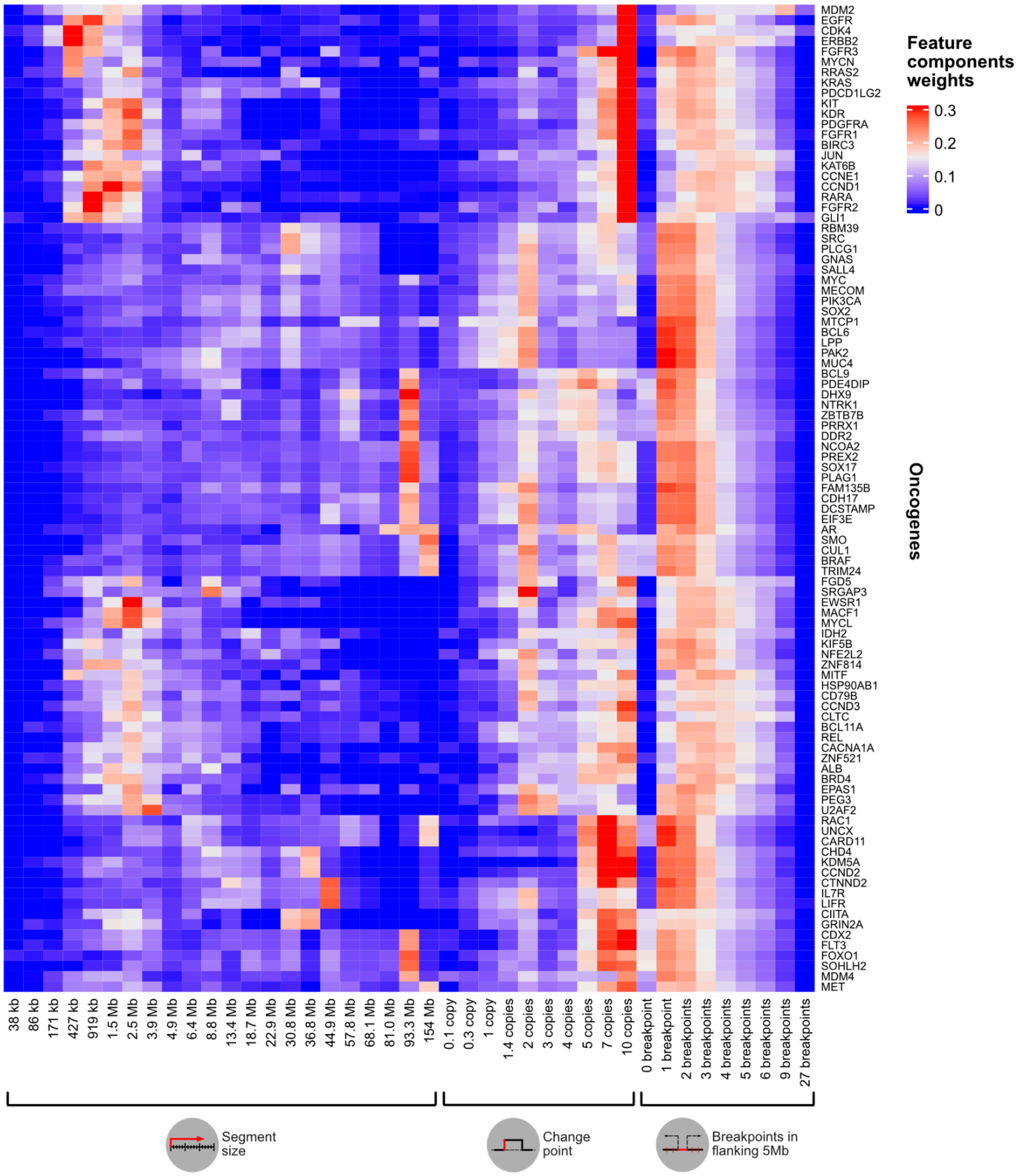


**Figure S14. Copy number feature distributions across oncogene amplifications.** Heatmap showing the mean of the sum-of-posterior probability component vectors of all copy number segments harbouring a particular oncogene amplification (oncogene-specific feature components weights). Weights are normalised per feature for ease in denoting the differences in terms of amplicon size, amplitude and density of flanking breakpoints across oncogenes. Displayed on the bottom are the three copy number features and their mixture components (22 components for the segment size feature, 10 components for the change point feature and 9 components for the flanking breakpoints feature). Mixture components were derived from feature distributions (see Methods).

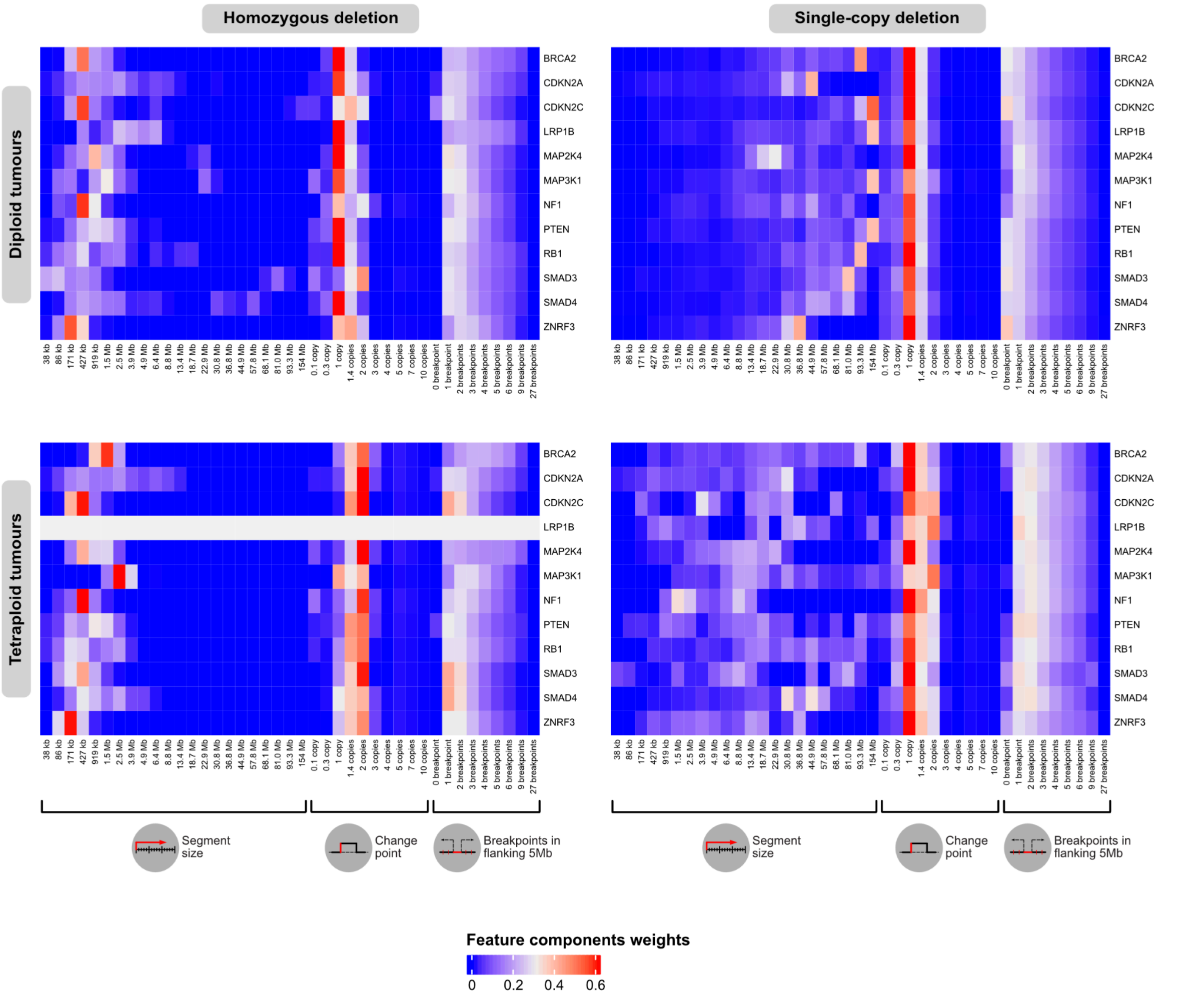


**Figure S15. Copy number feature distributions across tumour suppressor gene deletions.** Heatmaps showing the mean of the sum-of-posterior probability component vectors of all copy number segments harbouring homozygous (left) or single-copy deletions (right) in a given tumour suppressor gene (TSG-specific feature components weights). Weights are normalised per feature for ease in denoting the differences in terms of amplicon size, amplitude and density of flanking breakpoints across oncogenes. Displayed on the bottom are the three copy number features and their mixture components (22 components for the segment size feature, 10 components for the change point feature and 9 components for the flanking breakpoints feature). Mixture components were derived from feature distributions (see Methods).

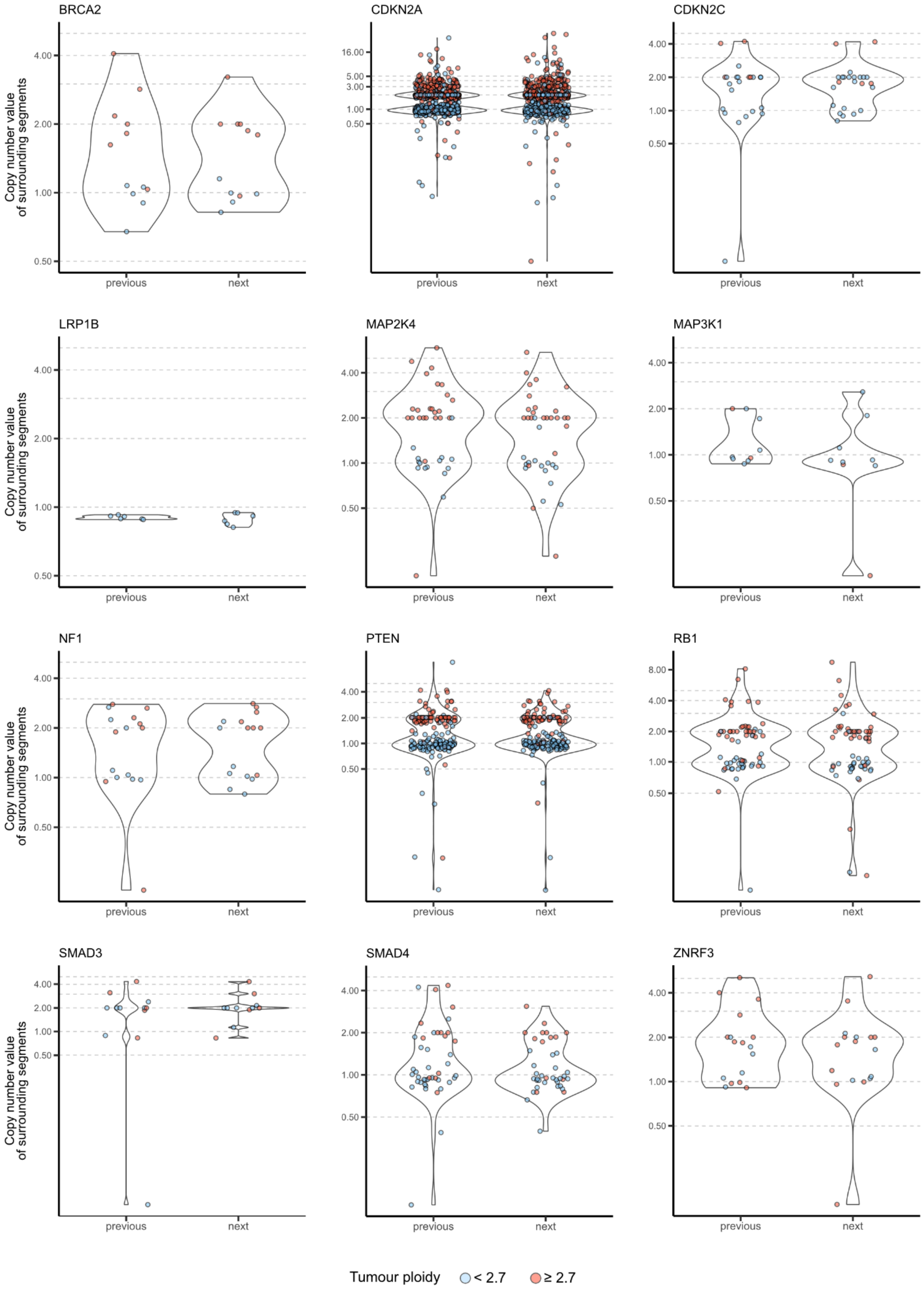


**Figure S16. Copy number values of segments surrounding TSG homozygous deletions.** Violin plots showing distribution of copy number values in the segments flanking homozygous deletions harbouring a given tumour suppressor gene (TSG). Each dot represents an adjacent segment, either preceding (previous) or following (next). Tumour samples are grouped as diploid (blue) or whole-genome duplicated (red). Dashed lines represent integer number states from 0.5 to 5 copies.

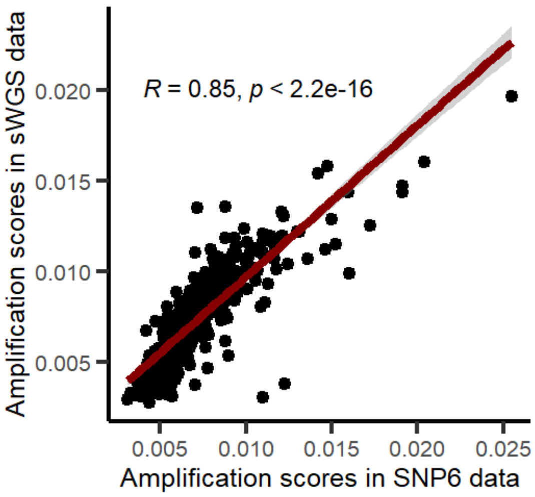


**Figure S17. Predictor stability across different copy number profiling technologies.** Scatter plot showing amplification scores derived from SNP6 (x-axis) and sWGS (y-axis) data from the same set of 478 tumours. Pearson correlation test used for comparing amplification scores between technologies. For comparison purposes, we averaged the oncogene-specific amplification scores per sample.

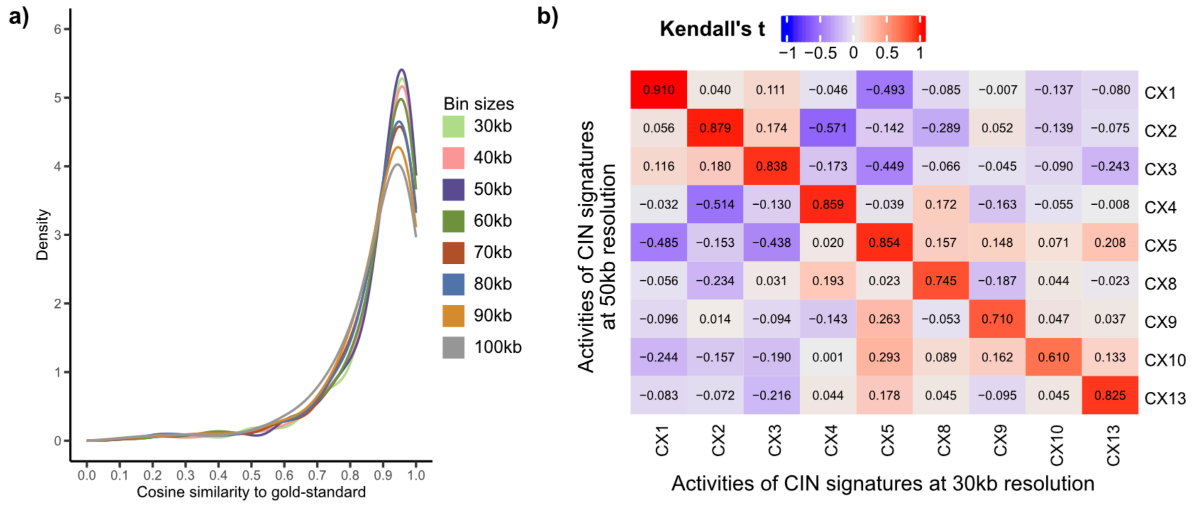


**Figure S18. Signature stability across different copy number profiling resolutions. a)** Density plot showing cosine similarities of CIN signature exposures in SNP6-derived profiles with those obtained from 478 sWGS-derived profiles using different bin resolutions. **b)** Heatmap showing Kendall's tau correlation coefficients between CIN signature exposures derived from sWGS data segmented using a bin size of 30 kb (x-axis) and 50 kb (y-axis). The same set of 478 tumours was used for comparisons. Red colour denotes a high positive correlation between exposures, while blue colour denotes a high negative correlation.

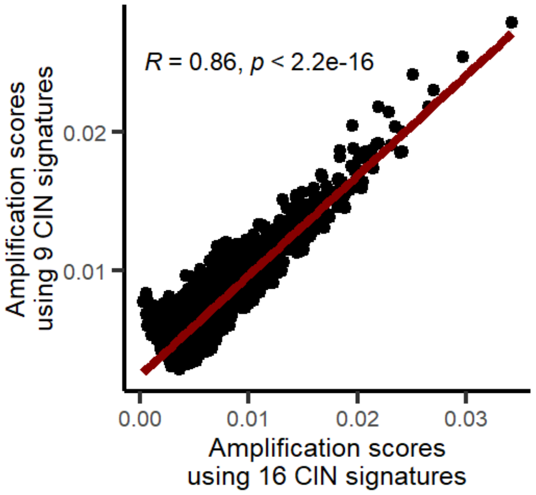


**Figure S19. Correlation between amplification scores computed using the full or the universal set of CIN signatures in the TCGA cohort.** Scatter plot showing amplification scores derived from the TCGA dataset using all 16 CIN signatures (x-axis) and those obtained using only the 9 universal signatures (y-axis). Pearson correlation test used for comparing amplification scores between signature sets. For comparison purposes, we averaged the oncogene-specific amplification scores per sample.

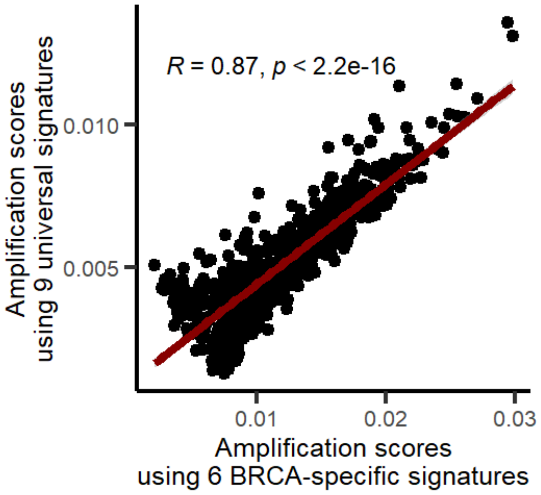


**Figure S20. Correlation between amplification scores computed using the breast-specific or the universal set of CIN signatures in the TCGA-BRCA cohort.** Scatter plot showing amplification scores derived from the TCGA-BRCA dataset using the signatures specifically extracted in the cohort (x-axis) and those obtained using the 9 universal signatures (y-axis). Pearson correlation test used for comparing amplification scores between signature sets. For comparison purposes, we averaged the oncogene-specific amplification scores per sample.

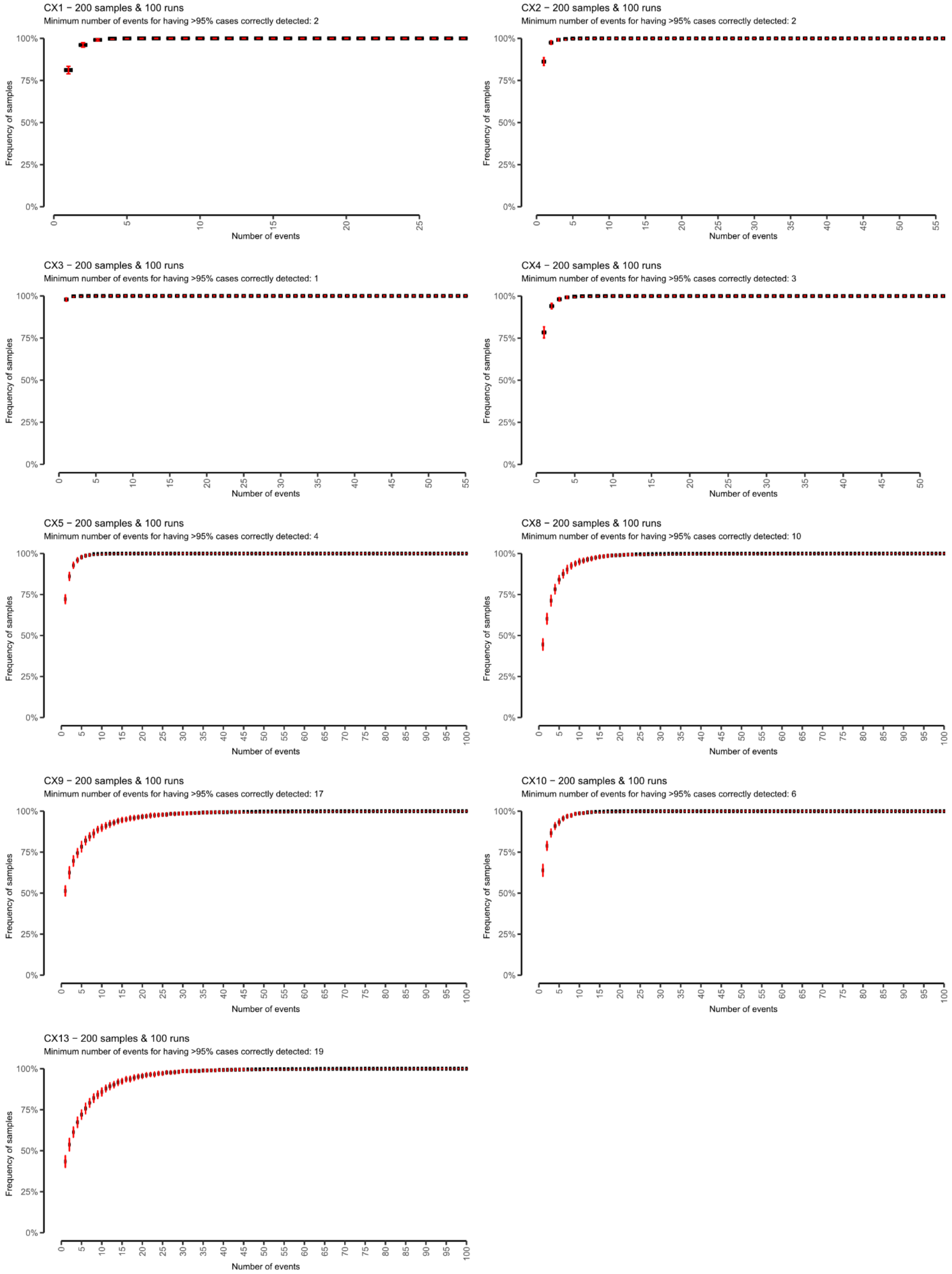


**Figure S21. Frequency of samples with correct detection of their dominant signature based on the number of events retained in the copy number profile.** Plot shows the success rate of detecting the dominant signature in a sample by progressively introducing an increasing number of events. For each specific signature, this procedure was repeated 100 times with a random selection of copy number events from the subset of 200 TCGA samples exhibiting the highest signature activity across the cohort. Black horizontal lines represent the mean across iterations, while red vertical lines indicate the standard deviation of the mean across iterations.

**a)** Amplification threshold independent to tumour's ploidy

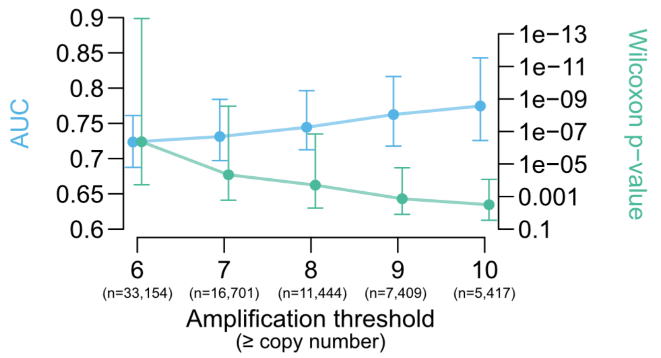


**b)** Amplification threshold based on tumour's ploidy

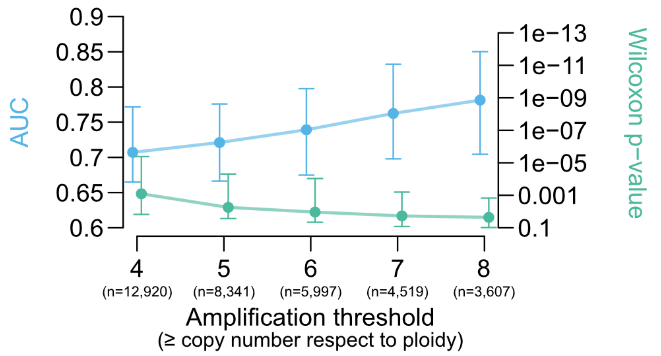


**Figure S22. Retrospective performance of our forecasting framework across a range of thresholds for defining focal amplifications. a)** Results from the ploidy-independent strategy. **b)** Results from the ploidy-aware strategy. Plots show the median and interquartile range across oncogenes in each amplification threshold. Performance was evaluated by comparing amplification scores between samples with and without oncogene amplifications using Wilcoxon two-sided test, and by computing the area under the receiver operating characteristic curve (AUC).

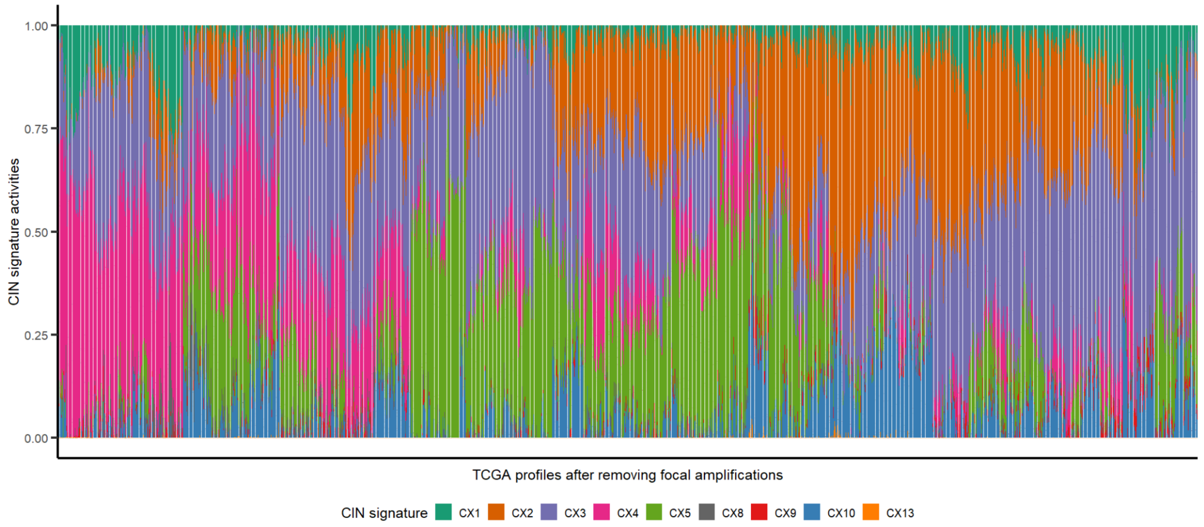


**Figure S23. CIN signature activities in the absence of focal amplifications.** Bar plot showing the activities of the 9 universal CIN signatures across the TCGA cohort after removing all focal amplifications (segments with copy number value equal or higher than 8).

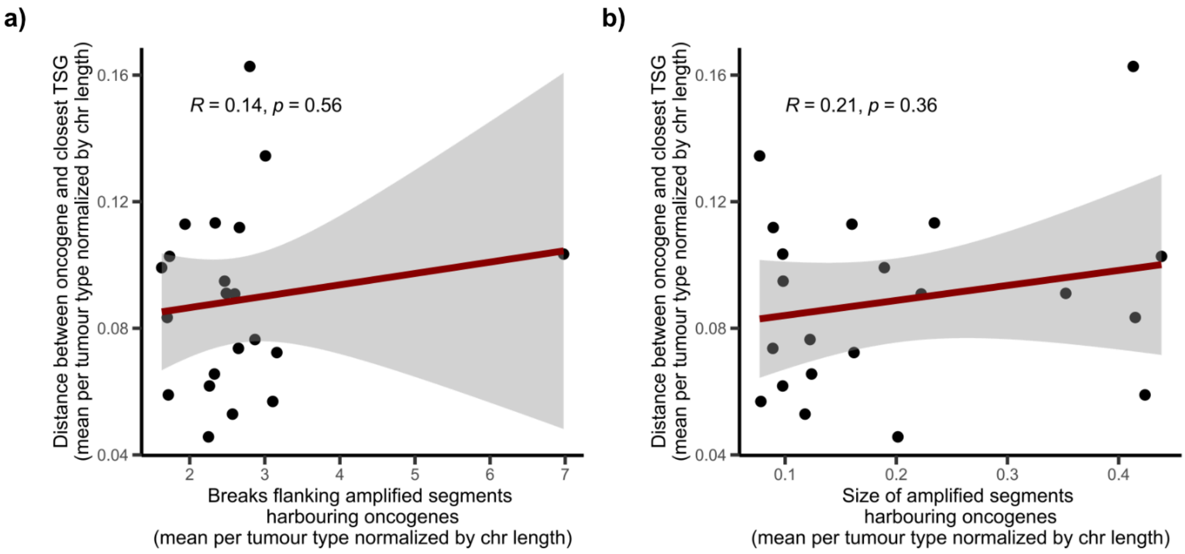


**Figure S24. Impact of genomic context on amplicon configurations.** Scatter plot showing **a)** flanking breaks and **b)** sizes of amplicons across tumour types (x-axis) with respect to the distance between oncogenes and their closest tumour suppressor genes (y-axis). Pearson correlation test was used for testing correlation significance. For comparison purposes, we averaged values across oncogenes per tumour type. Dots represent each tumour type.

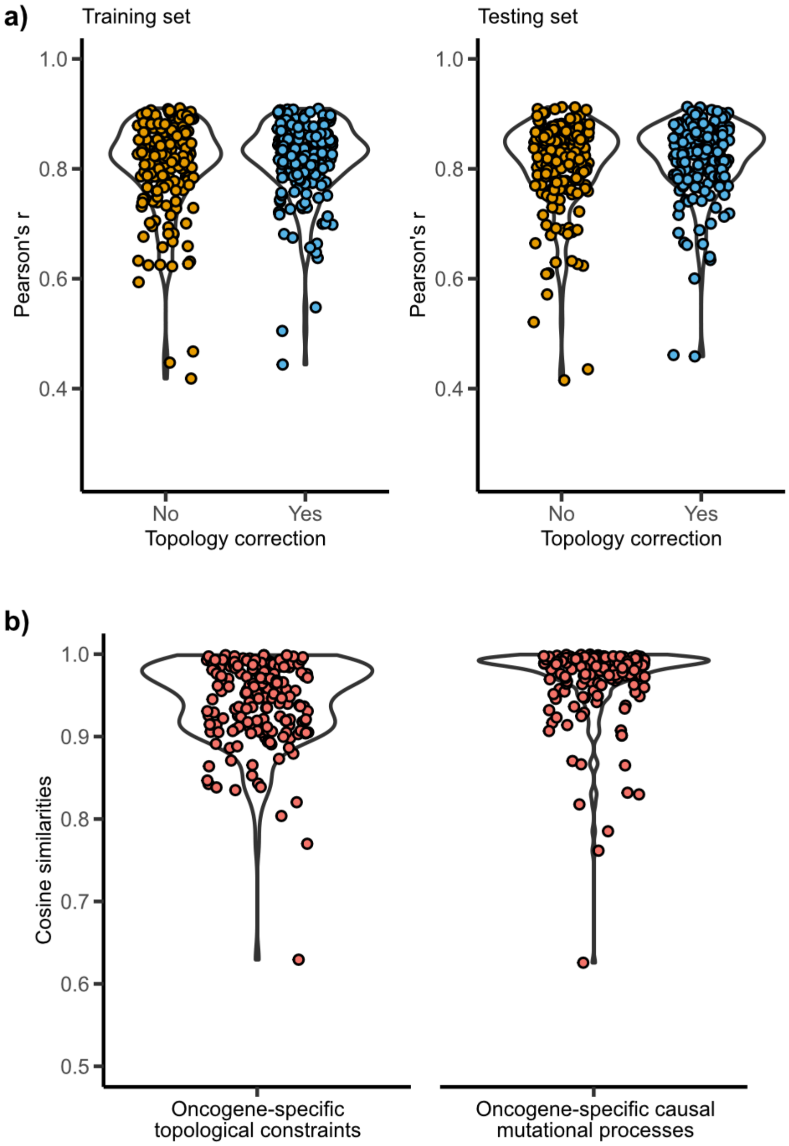


**Figure S25. Evaluation of the framework's ability to capture mutation rate using simulations. a)** Violin plots showing Pearson's correlation coefficients between simulated and estimated mutation rate across simulated cohorts. Mutation rate estimation was conducted with and without correction for topological constraints on locus-specific mutation rates. The model was trained on the training cohorts and applied to estimate mutation rates in both the training and testing cohorts (see Methods for details). **b)** Violin plots showing cosine similarities between simulated and estimated topological constraints in the oncogene region (left) and between simulated and estimated signatures causing the oncogene amplification (right).

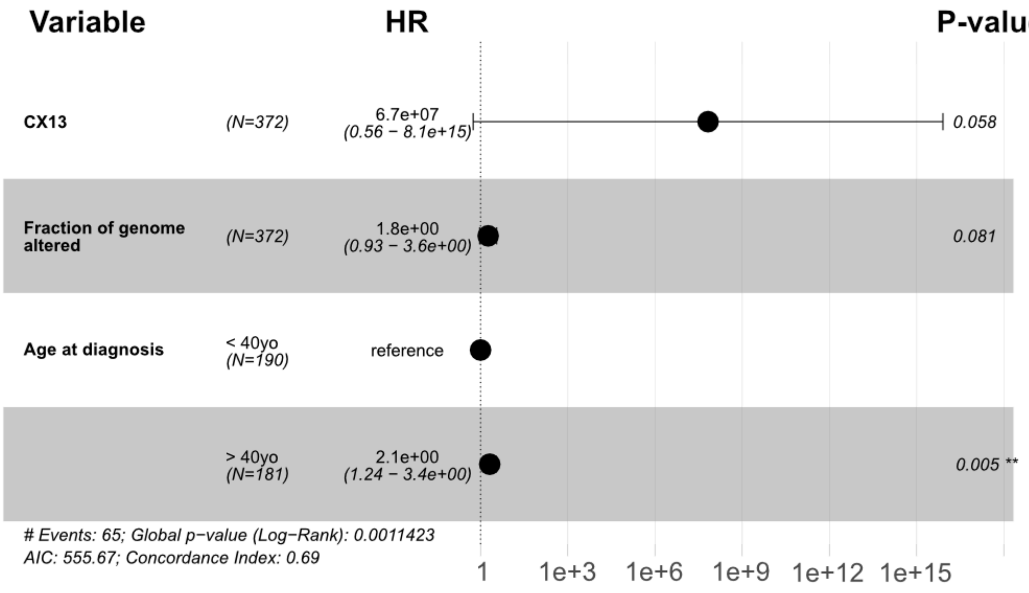


**Figure S26. Results of a Cox proportional hazards model for TCGA-LGG patients according to CX13 levels.** The model was corrected for the fraction of genome altered and age at diagnosis. Age at diagnosis was categorised into two groups (<40 versus ≥40 years) based on clinical risk.

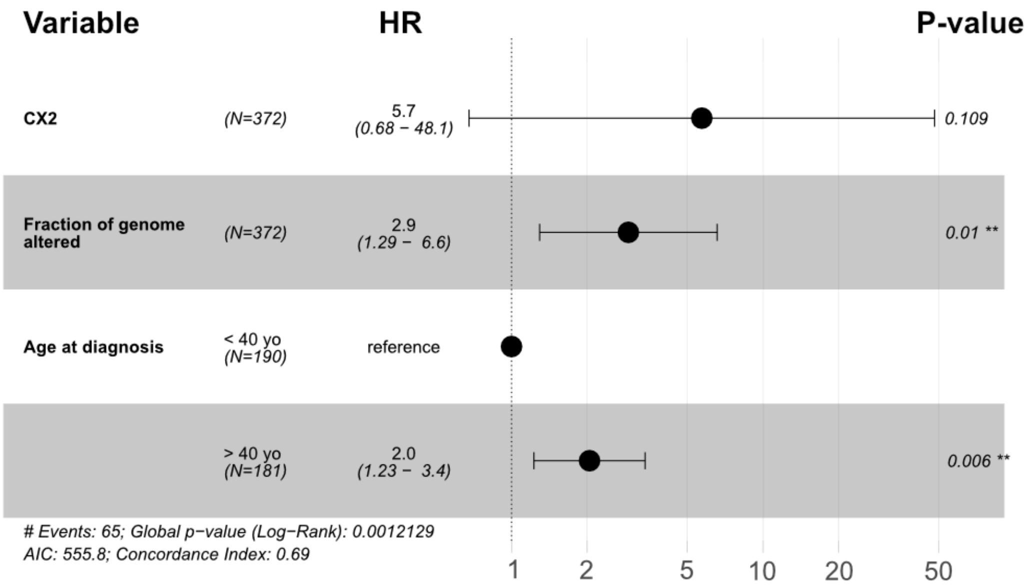


**Figure S27. Results of a Cox proportional hazards model for TCGA-LGG patients according to CX2 levels.** The model was corrected for the fraction of genome altered and age at diagnosis. Age at diagnosis was categorised into two groups (<40 versus ≥40 years) based on clinical risk.

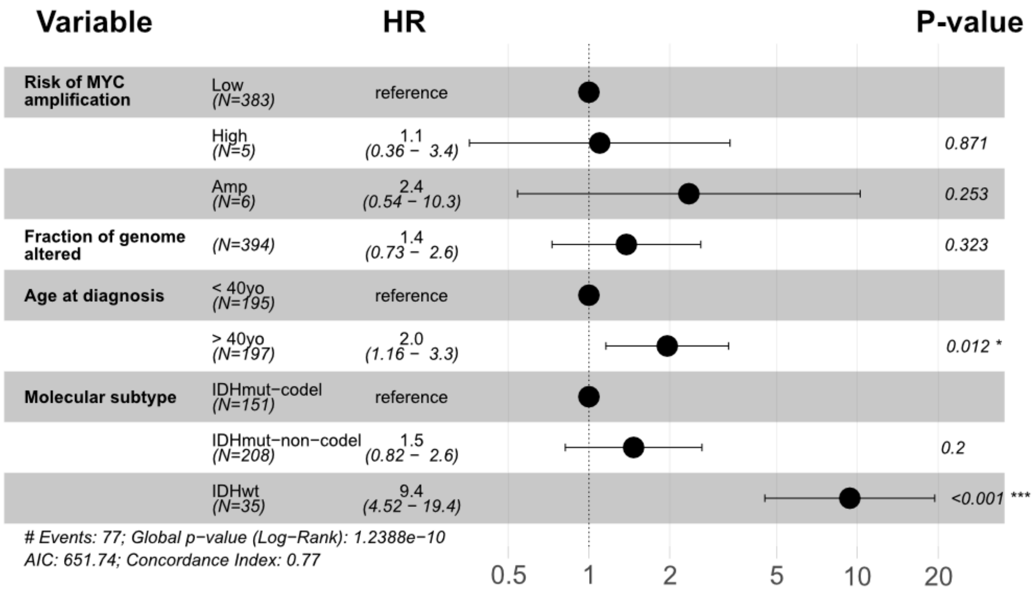


**Figure S28. Results of a Cox proportional hazards model for TCGA-LGG patients with and without *MYC* amplification.** Patients without *MYC* amplification are classified as high or low risk of acquiring *MYC* amplification in the future by our forecasting framework. The model was corrected for the fraction of genome altered, age at diagnosis and molecular subtype. Age at diagnosis was categorised into two groups (<40 versus ≥40 years) based on clinical risk.

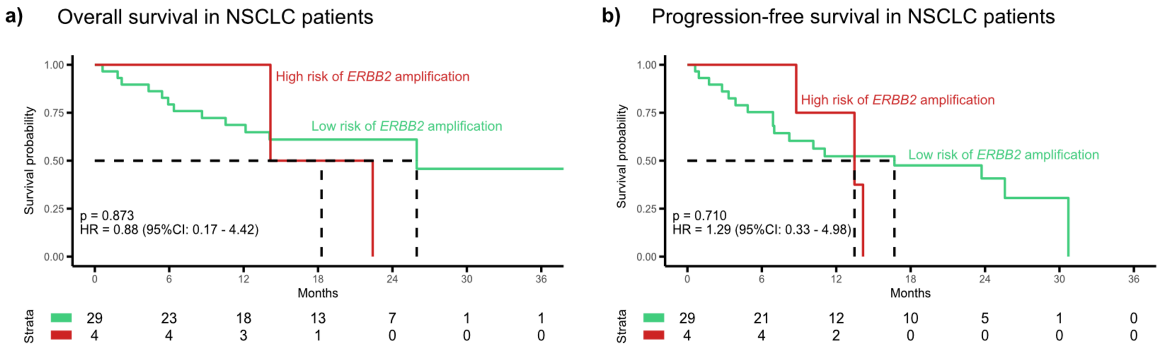


**Figure S29. Predicting EGFR tyrosine kinase inhibitor resistance via forecasting *ERBB2* amplification.** Kaplan-Meier curves showing (right) overall survival and (left) progression-free survival for NSCLC patients from the OSIRESP cohort predicted as having high (red) or low (green) risk of acquiring *ERBB2* amplification in the future. Cox proportional hazards regression models were applied to predict hazard ratios (HR) and p-values (p). The model was corrected for the fraction of genome altered, the presence of metastasis in the central nervous system (CNS) at treatment initiation, and the physical status based on the Eastern Cooperative Oncology Group (ECOG) scale at treatment initiation.

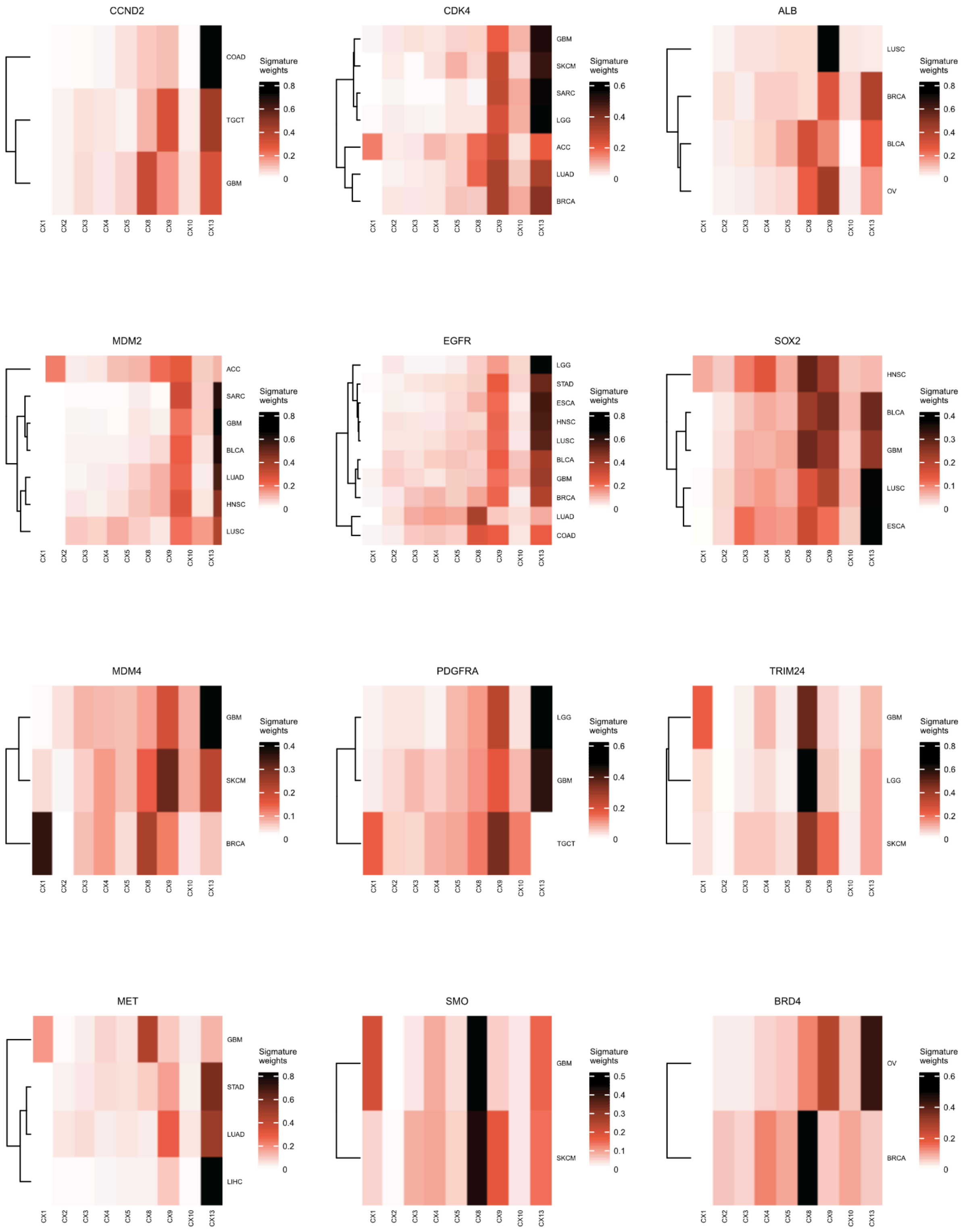


**Figure S30. Types of chromosomal instability underpinning oncogene amplification (Part 1).** Heatmaps represent the signature enrichment in amplified segments mapping oncogenes across tumour types. Only tumour types in which the given oncogene putatively confers a fitness advantage are included.

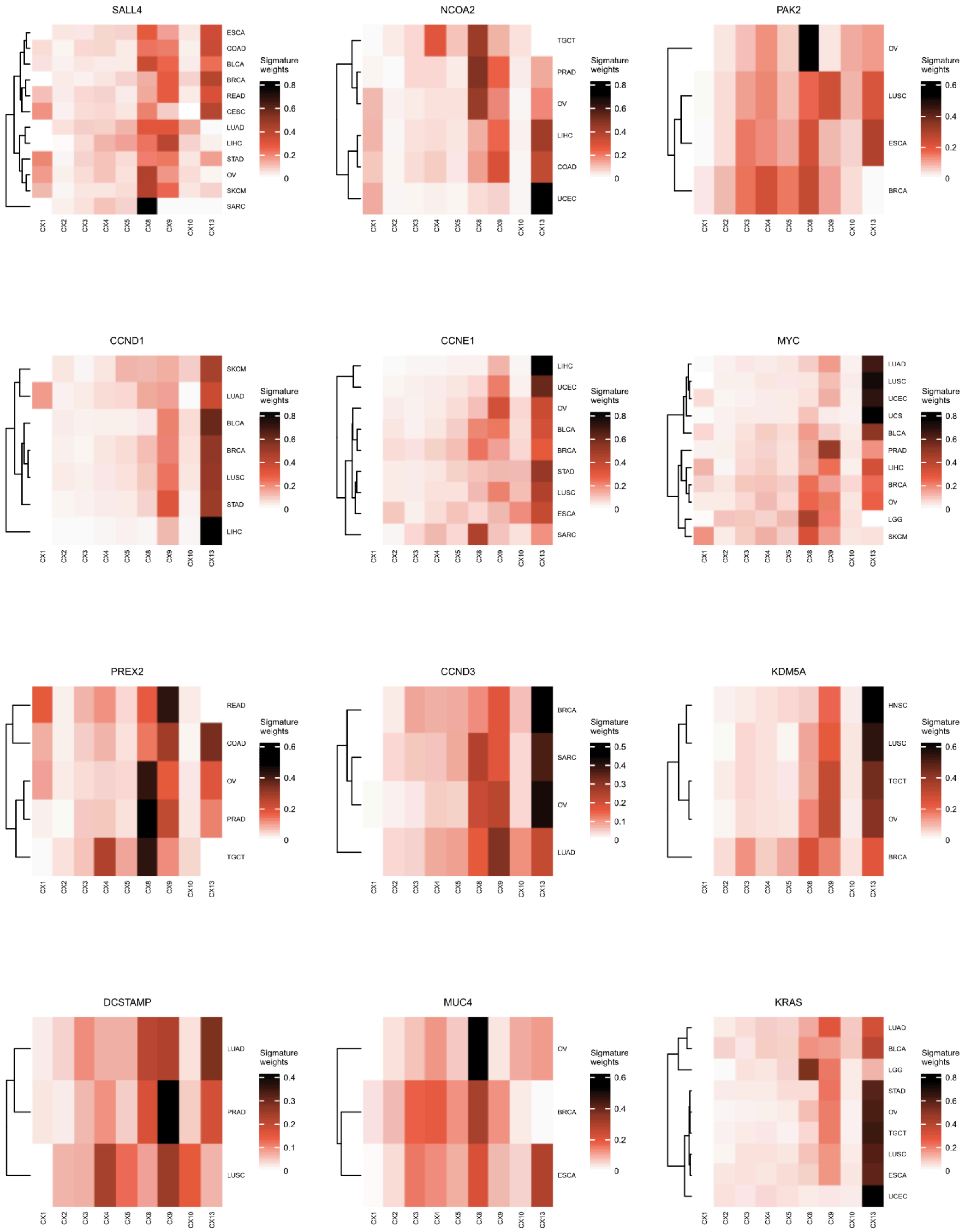


**Figure S30. Types of chromosomal instability underpinning oncogene amplification (Part 2).** Heatmaps represent the signature enrichment in amplified segments mapping oncogenes across tumour types. Only tumour types in which the given oncogene putatively confers a fitness advantage are included.

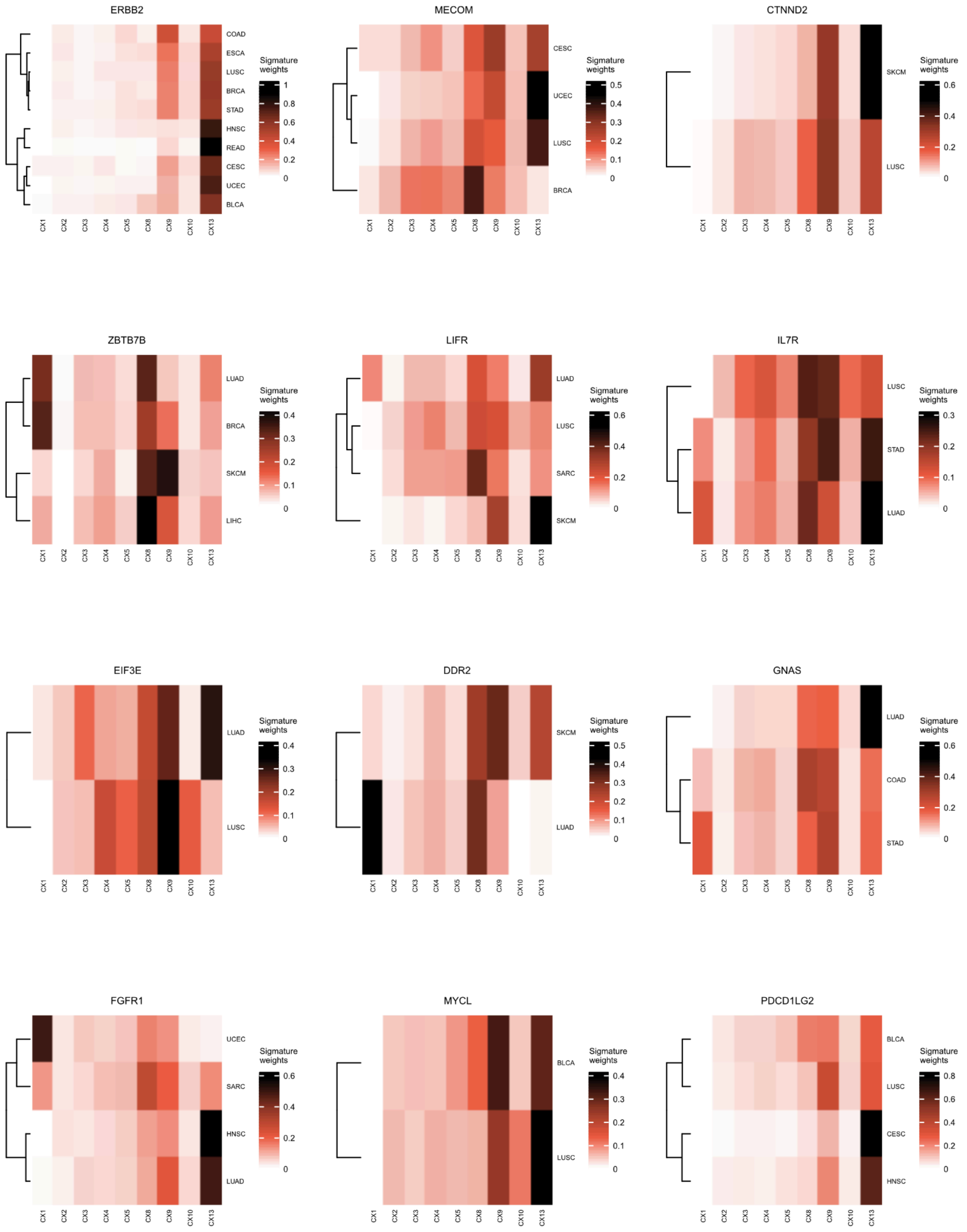


**Figure S30. Types of chromosomal instability underpinning oncogene amplification (Part 3).** Heatmaps represent the signature enrichment in amplified segments mapping oncogenes across tumour types. Only tumour types in which the given oncogene putatively confers a fitness advantage are included.

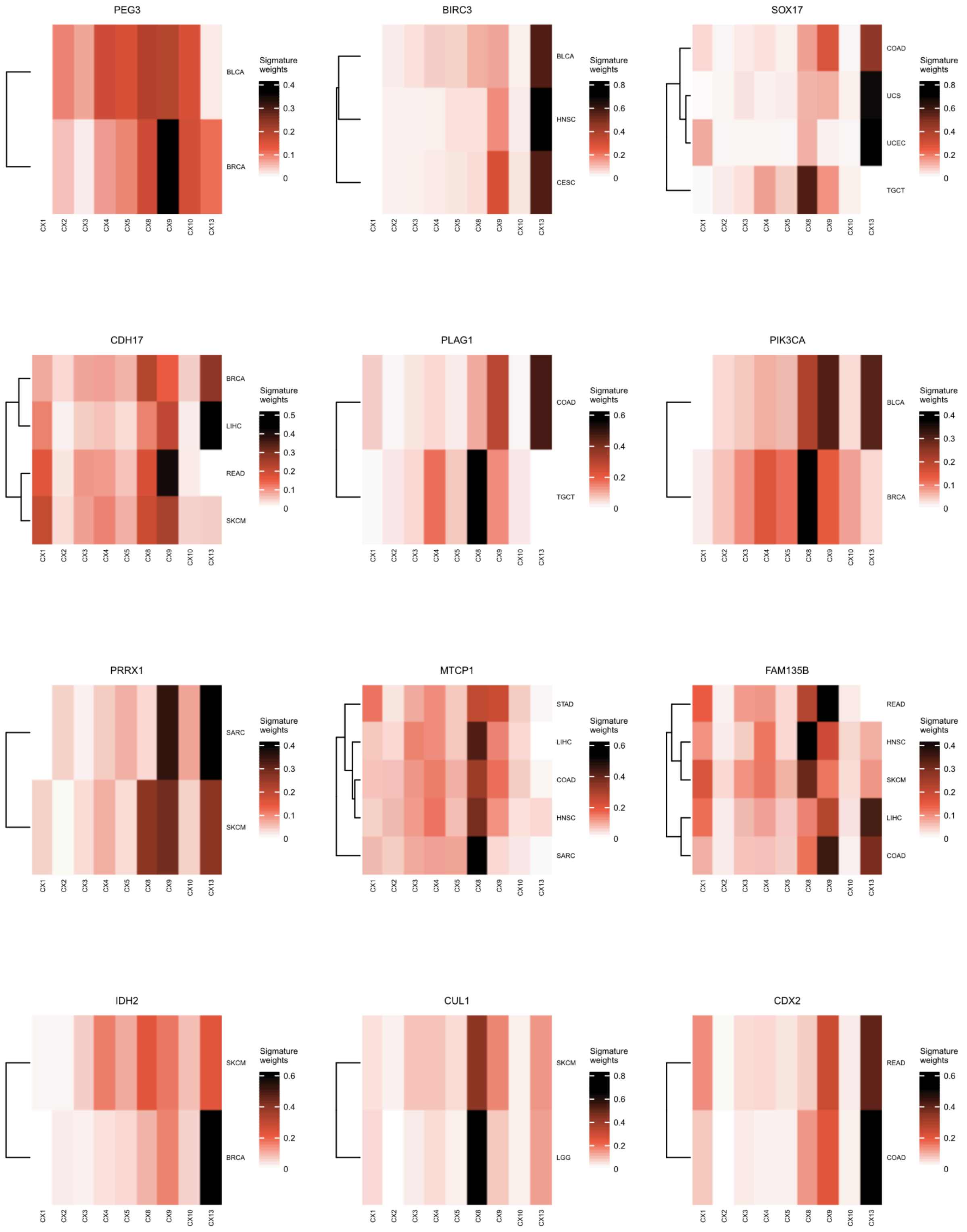


**Figure S30. Types of chromosomal instability underpinning oncogene amplification (Part 4).** Heatmaps represent the signature enrichment in amplified segments mapping oncogenes across tumour types. Only tumour types in which the given oncogene putatively confers a fitness advantage are included.

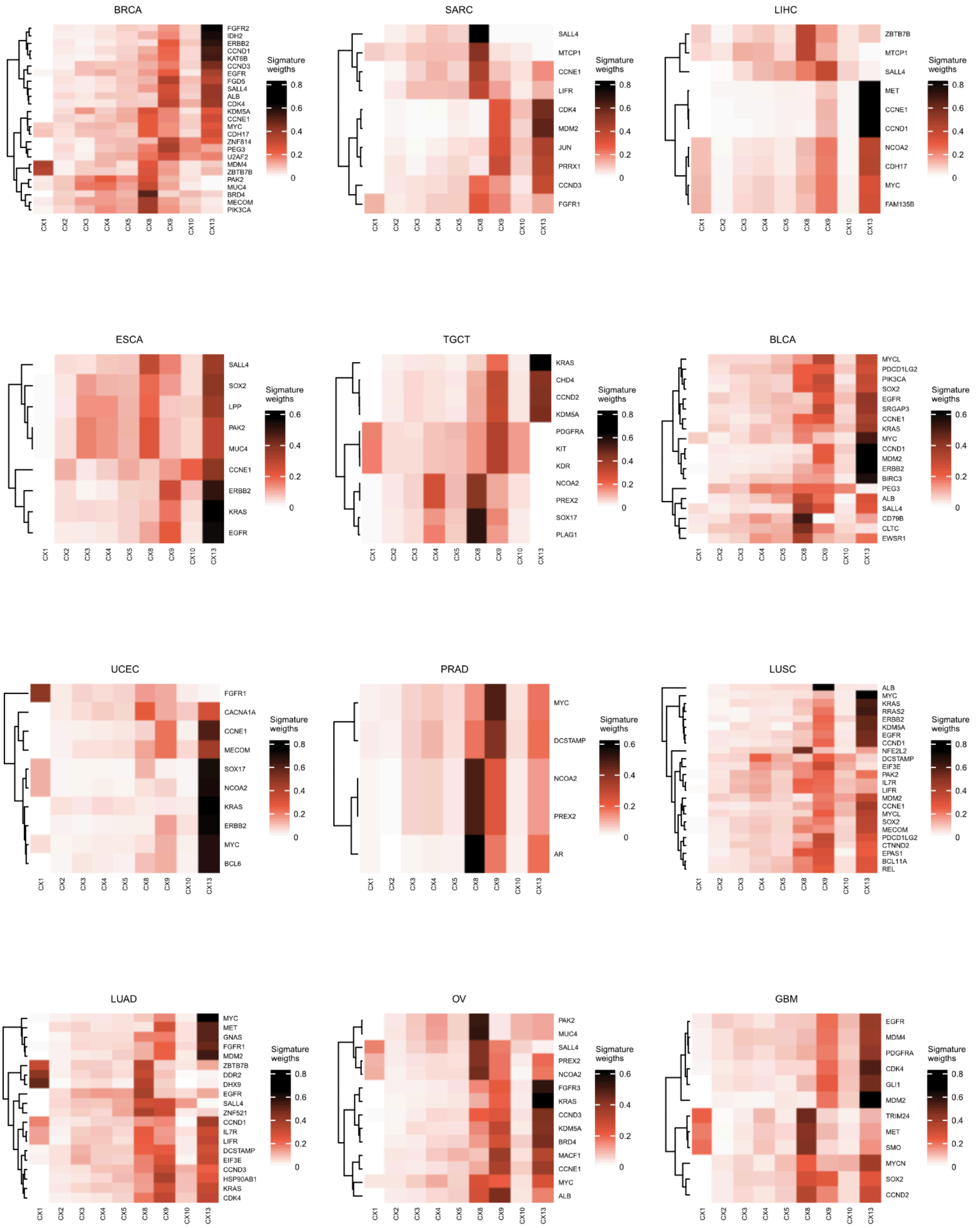


**Figure S31. Types of chromosomal instability underpinning oncogene amplification per tumour type (Part 1).** Heatmaps represent the signature enrichment in amplified segments mapping different oncogenes in a given tumour type. Only oncogenes positively selected to be amplified in a tumour type are included.

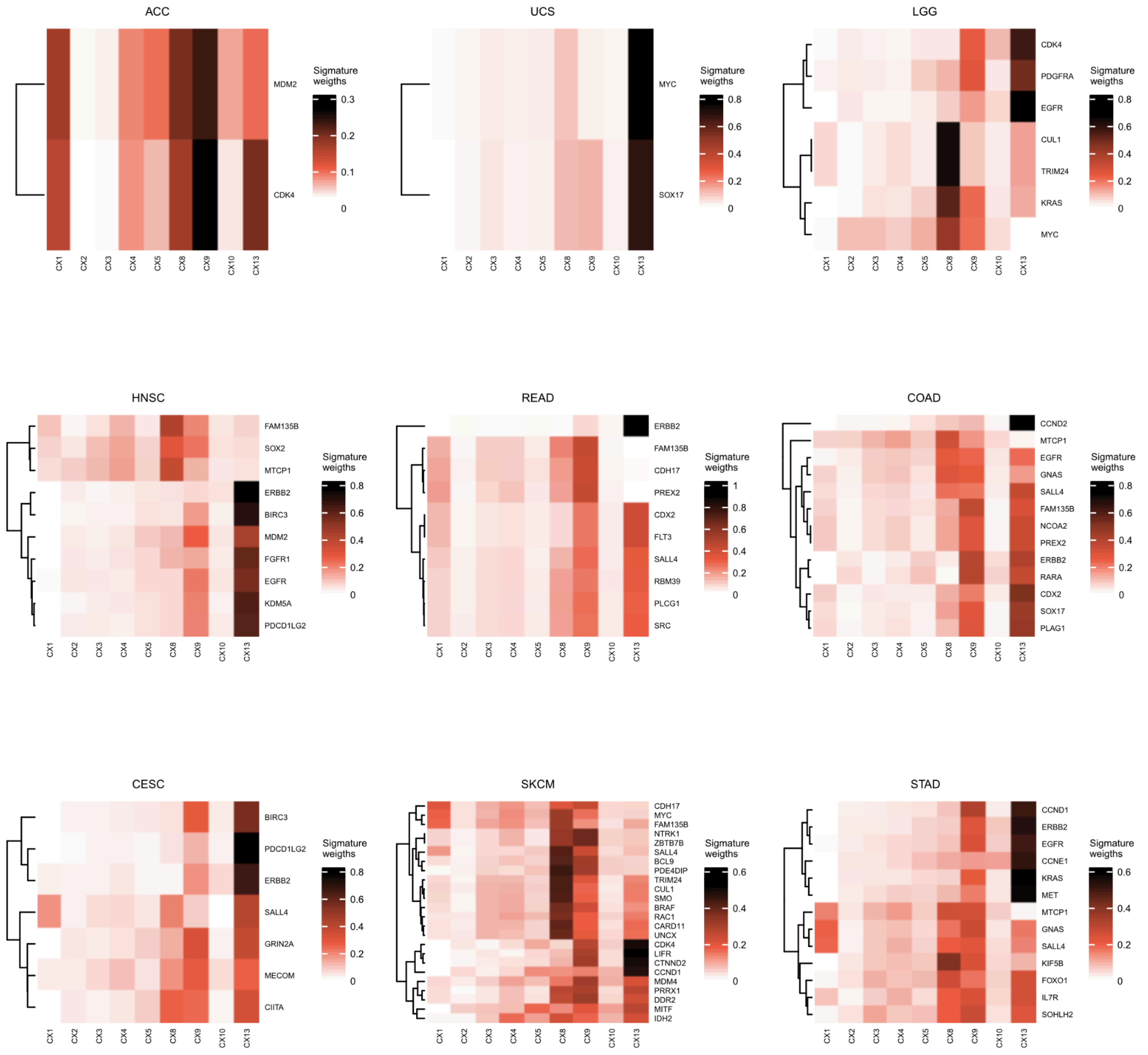


**Figure S31. Types of chromosomal instability underpinning oncogene amplification per tumour type (Part 2).** Heatmaps represent the signature enrichment in amplified segments mapping different oncogenes in a given tumour type. Only oncogenes positively selected to be amplified in a tumour type are included.

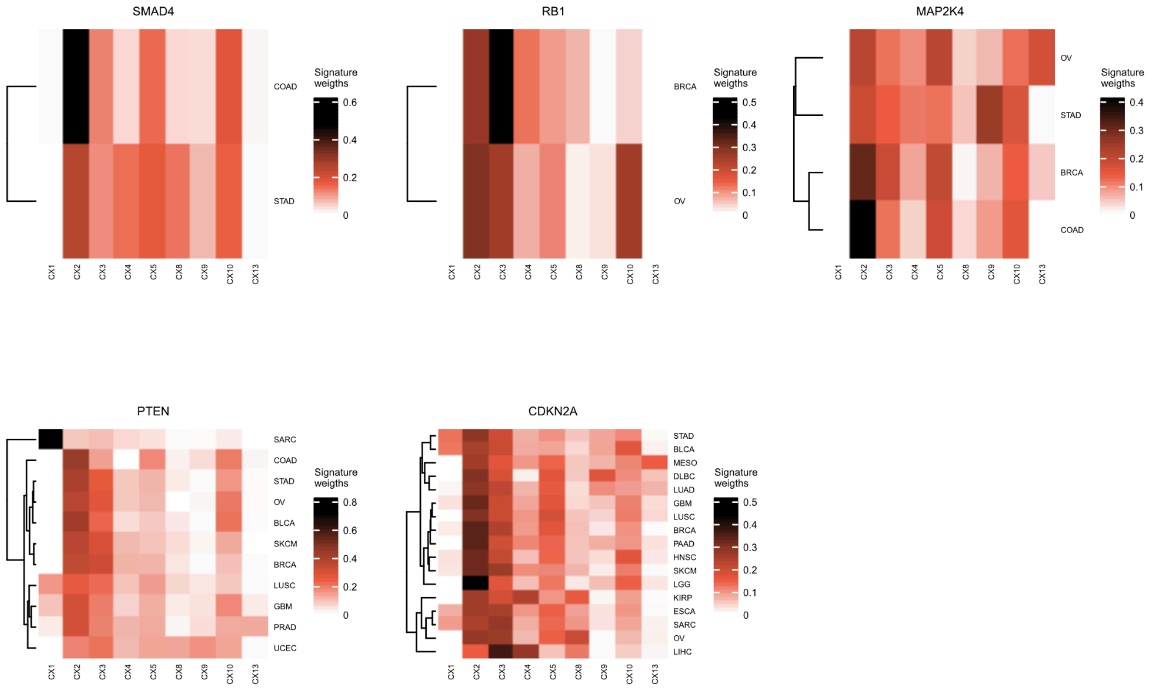


**Figure S32. Types of chromosomal instability underpinning homozygous deletion of tumour suppressor genes.** Heatmaps represent the signature enrichment in deleted segments mapping tumour suppressor genes across tumour types. Only tumour types in which the given tumour suppressor gene putatively confers a fitness advantage are included.

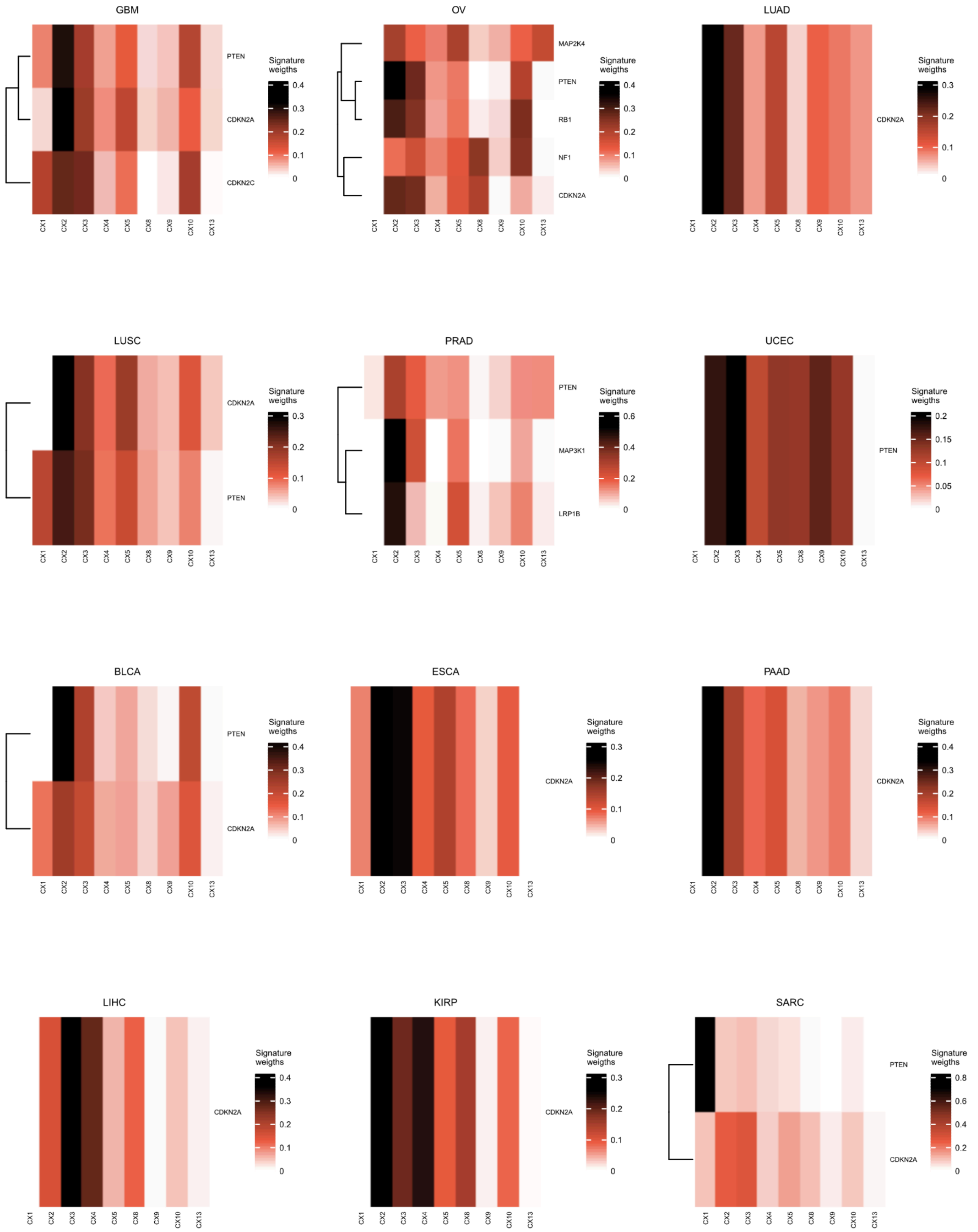


**Figure S33. Types of chromosomal instability underpinning homozygous deletion of tumour suppressor genes per tumour type (Part 1).** Heatmaps represent the signature enrichment in deleted segments mapping different tumour suppressor genes in a given tumour type. Only tumour suppressor genes positively selected to be deleted in a tumour type are included.

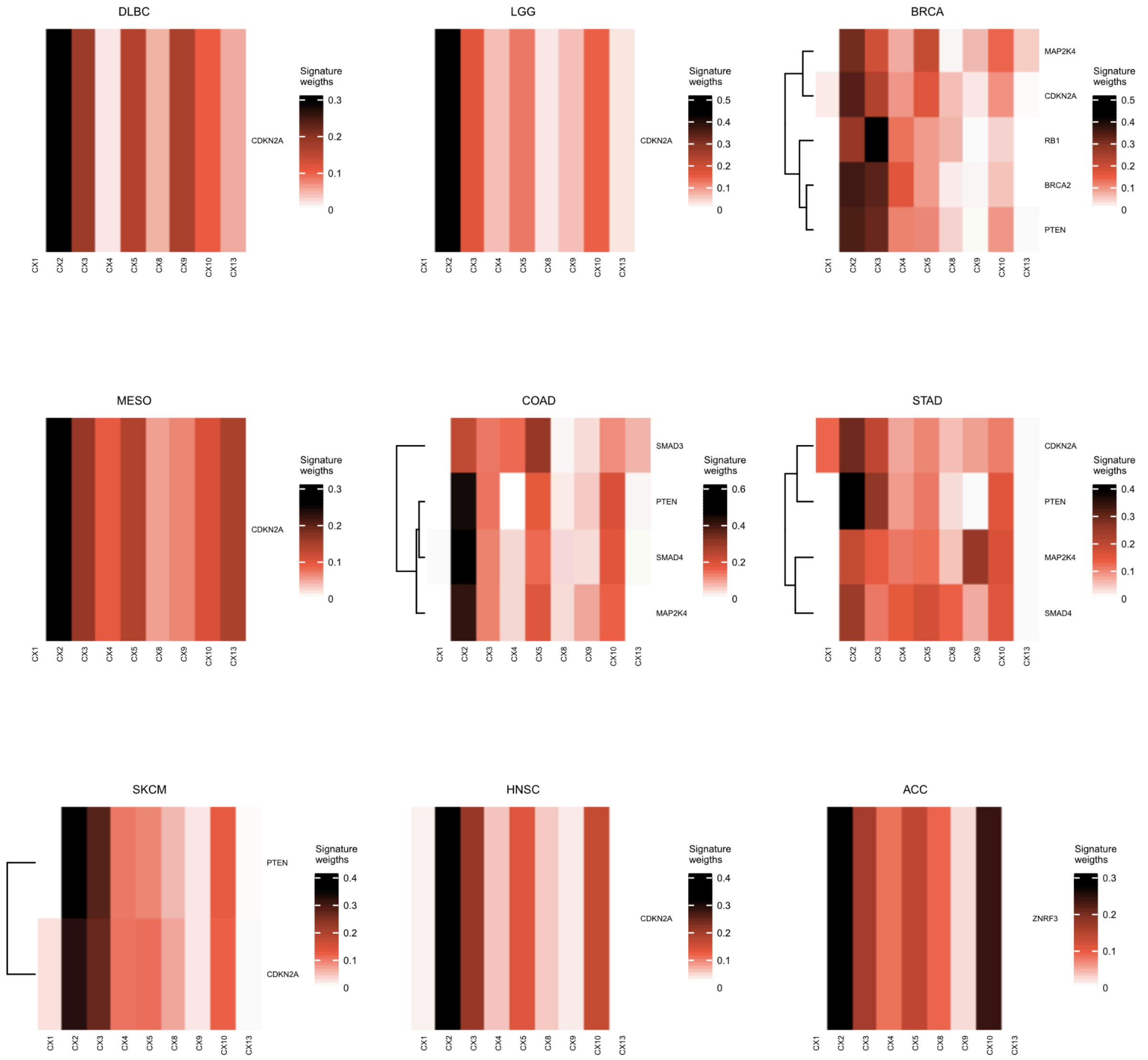


**Figure S33. Types of chromosomal instability underpinning homozygous deletion of tumour suppressor genes per tumour type (Part 2).** Heatmaps represent the signature enrichment in deleted segments mapping different tumour suppressor genes in a given tumour type. Only tumour suppressor genes positively selected to be deleted in a tumour type are included.

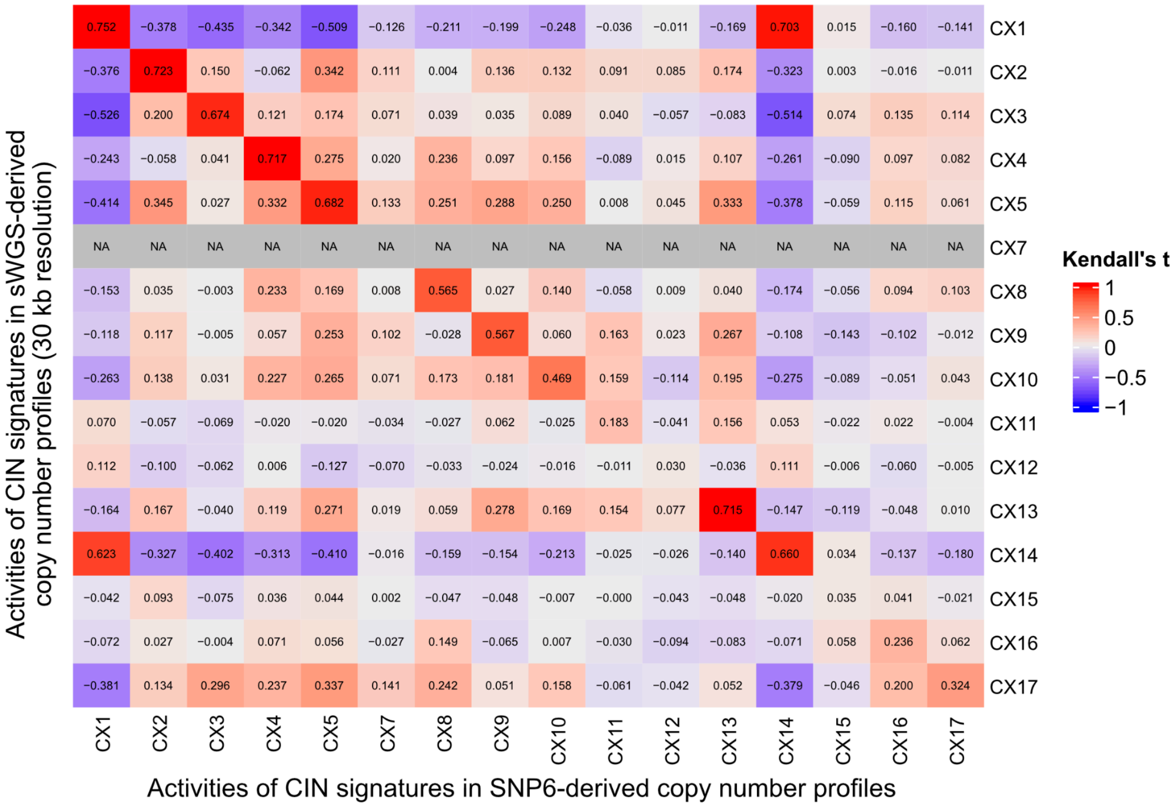


**Figure S34. Signature stability across different copy number profiling technologies.** The heatmap shows Kendall's tau correlation coefficients between CIN signature activities derived from SNP6 (x-axis) and sWGS (y-axis) data from the same set of 478 tumours. Red colour denotes a high positive correlation between activities, while blue colour denotes a high negative correlation. CX7 was exclusively identified in SNP6-derived copy number profiles.

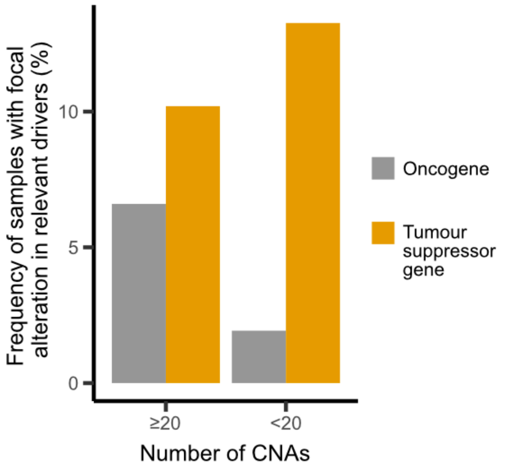


**Figure S35. Frequency of samples with focal copy number alterations in driver genes according to CIN levels.** Bar plot showing the frequency of TCGA samples with oncogene amplifications (grey) and homozygous deletions in tumour suppressor genes (yellow). Tumour samples are grouped as having detectable CIN (≥20 copy number alterations, CNAs) or not having detectable CIN (<20 CNAs).

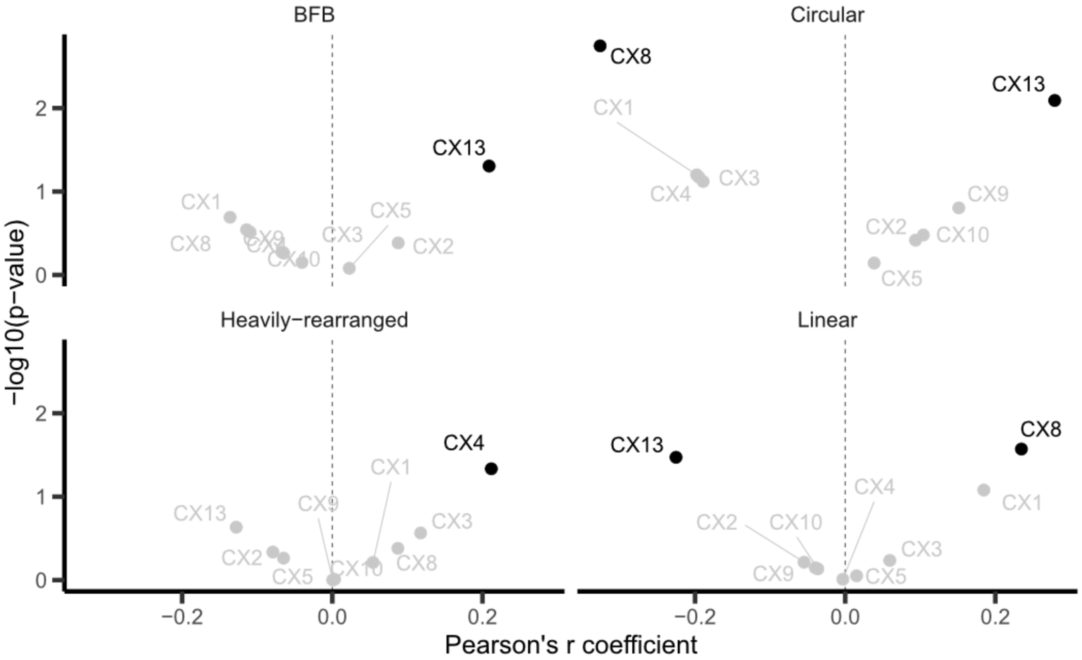


**Figure S36. Associations between signatures mapped to oncogene amplifications with amplicon structure.** Scatter plots showing the Pearson correlation between oncogene-specific signature weights and four amplicon types. Black colour denotes significant correlation.
