## Supplementary material for "Forecasting oncogene amplification and tumour suppressor deletion": Methods


**Data and code availability** **2**
**Data reporting and sources** **2**
**Software** **4**
**Experimental data** **6**
Patient samples and cell lines 6
Patient sample cohort 6
Human cancer cell lines 6
DNA extraction 7
DNA sequencing 7
Shallow whole genome sequencing (sWGS) 7
Whole exome sequencing (WES) 8
Single-cell whole genome sequencing (scWGS) 8
Read alignment 9
**Generating copy number profiles** **9**
Absolute copy number fitting from sWGS 9
Absolute copy number fitting from WES 9
Absolute copy number fitting from scWGS 10
Absolute copy number fitting from pseudobulk WGS 10
**Detection of focal amplifications** **10**
**Detection of homozygous deletions** **11**
**Forecasting driver gene amplifications and deletions** **11**
Estimating locus-specific mutation rates (μ) 12
Quantifying tumour mutational processes activity (ICX) 13
Estimating locus-specific mutational processes (ωD,CX) 13
Estimating causal mutational processes (ZD,CX) 14
Estimating topological constraints (XD,CX) 16
Estimating selection coefficients (s) 17
**Robustness analyses** **18**
Assessing stability across technologies 18
Assessing stability across bin resolutions 18
Robustness of using the universal set of signatures 19
Assessing signature assignment to individual events 20
Assessing number of individual events required for sample-level signature quantification 21
Assessing the threshold for defining focal amplifications 21

Assessing whether mutational processes can be detected in the absence of focal amplifications 22
Assessing the impact of the genomic context on amplicon configurations 22
Assessing the ability to capture mutation rates 23
**Model performance assessment 25**
Testing the predictive capacity of the forecasting framework 25
Assessing forecasting performance of the framework 27
Lung cancer longitudinal data 27
Prostate cancer longitudinal data 28
Hartwig Medical Foundation prostate tumour samples 28
Mateo et al. 2020 29
Pan-Prostate Cancer Group cohort 29
Integrated analysis 29
**Demonstrating clinical utility of the predictor 30**
Predicting prognosis 30
Predicting amplification-driven drug resistance 32
Lung cancer cell lines 32
Lung cancer patients 33
**Exploring mutational processes underlying driver alterations 34**
Exploring processes underlying oncogene amplifications 34
Exploring processes underlying tumour suppressor gene deletions 35
Supporting putative processes underlying oncogene amplifications 35
**Exploring acquisition and selection of oncogene amplifications 36**
Amplification acquisition 36
Positive selection test 36
Distribution of unique amplifications across the genome 37
Birth-death simulations 37
**References 39**

### Data and code availability

The source code for reproducing analyses and figures will be available at https://github.com/macintyrelab. All data required for reproducing these analyses will be available at figshare. Raw data generated in this study (both sWGS and WES data from lung cancer patients and cell lines) will be deposited in the European Genome-phenome Archive (EGA) under the accession number EGAXXXXXXXXXXX. Raw sequencing data generated by the TCGA portion of the PCAWG project have restricted access (TCGA dbGaP accession number: phs000178.v11.p8). Access can be obtained by applying to the TCGA Data Access Committee (DAC) via dbGaP. Raw sequencing data generated by Mateo *et al.*[1] is deposited at the European Nucleotide Archive under the accession number PRJEB32038. Raw sequencing data generated by Jia *et al.*[2] is deposited at the NCBI Sequence Read Archive under accession numbers SRP022942 and SRP022943. Access to raw, processed and clinical data from the Hartwig Medical Foundation can be requested here. All other data supporting the findings of this study are publicly available without restrictions.

### Data reporting and sources

No statistical methods were used to predetermine sample size, and the experiments were not randomised.

Here, we list the data used in this study and their respective sources. All data are publically accessible, except for the Pan Cancer Analysis of Whole Genomes (PCAWG) raw data (WGS) and the Hartwig Medical Foundation (HMF) clinical, processed and raw data (WGS). The access to the TCGA portion of the PCAWG data can be obtained by a Data Access Request through the Database of Genotypes and Phenotypes (dbGaP).

**Methods Table 1. Sources of data used in this study**

| **Data** | **Source** |
|---|---|
| TCGA copy number profiles | Drews et al., 2022[3]. Files hosted on https://github.com/VanLoo-lab/ascat |
| TCGA clinical data | TCGA Research Network: https://www.cancer.gov/tcga |
| Molecular subtype for TCGA low glioma cancer samples | Brain Lower Grade Glioma TCGA PanCancer data. Files downloaded via cBioportal: https://www.cbioportal.org/study/summary?id=lgg_tcga_pan_can_atlas_2018 |
| Copy number status of *CDKN2A* and *CDKN2B* for TCGA low glioma cancer samples | Brain Lower Grade Glioma TCGA PanCancer data. Files downloaded via cBioportal: https://www.cbioportal.org/study/summary?id=lgg_tcga_pan_can_atlas_2018 |
| PCAWG copy number profiles | Gerstung, et al. 2020[4]; Dentro, et al. 2021[5]. Files hosted on https://dcc.icgc.org/releases/PCAWG |
| PCAWG SV amplicons annotation | Kim et al., 2020[6] |

| | |
|---|---|
| PCAWG clinical annotation | Pan-Cancer Analysis of Whole Genomes Consortium[7]. Files hosted on https://dcc.icgc.org/releases/PCAWG/ |
| PPCG copy number profiles | Access to the data can be requested to the Pan-Prostate Cancer Group: http://panprostate.org/index.html |
| GLASS copy number profiles | Barthel et al., 2019[8]. Files hosted on synapse.org under the project ID syn17038081 |
| GLASS clinical data | Barthel et al., 2019[8]. Files hosted on synapse.org under the project ID syn17038081 |
| TRACERx lung copy number profiles | Al Bakir et al., 2023[9]. Files hosted on zenodo.org (https://doi.org/10.5281/zenodo.7649257) |
| Hartwig Medical Foundation copy number profiles | Priestley *et al.*, 2019[10]. Access to the data can be requested to the Hartwig Medical Foundation: https://www.hartwigmedicalfoundation.nl/en/data/data-access-request/ |
| Hartwig Medical Foundation clinical data | Priestley *et al.*, 2019[10]. Access to the data can be requested to the Hartwig Medical Foundation: https://www.hartwigmedicalfoundation.nl/en/data/data-access-request/ |
| Patient-matched primary and metastatic prostate cancer samples | Mateo *et al.*, 2020[1]. Files hosted on the European Nucleotide Archive with accession number PRJEB32038 |
| Raw single-cell DNA sequencing data from TNBC tumours and cell lines | Minussi et al., 2021[11]. Files hosted on the NCBI Sequence Read Archive under the accession number PRJNA629885 |
| Driver gene classification | Bailey et al., 2018[12]; Jassal et al., 2020[13]; Martínez-Jiménez et al., 2020[14]; Sondka et al., 2018[15] |
| List of non-cancer oncogenes | Ensembl GRCh37 genome browser: https://grch37.ensembl.org/index.html |
| GISTIC2.0 outputs for the TCGA cohort | Mermel et al., 2011[16]. Access to the results: https://portals.broadinstitute.org/tcga/home |
| Genomic and clinical data of a subset of patients from the OsiResp project | Generated in this project (see section "Experimental data") |
| Genomic data of parental and resistant lung cancer cell lines | Generated in this project (see section "Experimental data"), and publicly available in Jia et al., 2013[2]. Files hosted on the NCBI Sequence Read Archive under accession numbers SRP022942 and SRP022943. |

### Software

If not otherwise stated, all analyses were performed using the free statistical software R (v>4.0). If functions outside the standard packages were used, they are mentioned and cited throughout the text. A list of all packages used and their corresponding versions are presented in **Methods Table 2**. All software written by us as part of this study will be available at https://github.com/macintyrelab.

**Methods Table 2: R packages used as part of this study**

| Name | Version | Citation | Name | Version | Citation |
|---|---|---|---|---|---|
| QDNAseq | 1.21.0 | [17] | survminer | 0.4.9 | [18] |
| this.path | 1.2.0 | [19] | survival | 3.5-5 | [20] |
| BiocGenerics | 0.44.0 | [21] | PRROC | 1.3.1 | [22] |
| sysfonts | 0.8.8 | [23] | gridExtra | 2.3 | [24] |
| QDNAseqmod | 1.21.0 | Repo link | ggbeeswarm | 0.7.1 | [25] |
| randomcoloR | 1.1.0.1 | [26] | pROC | 1.18.0 | [27] |
| ROCit | 2.1.1 | [28] | GenomeInfoDb | 1.34.8 | [29] |
| CNpare | 0.99.0 | [30] | lubridate | 1.9.2 | [31] |
| forcats | 1.0.0 | [32] | stringr | 1.5.0 | [33] |
| purrr | 1.0.1 | [34] | readr | 2.1.4 | [35] |
| tidyr | 1.3.0 | [36] | tibble | 3.2.1 | [37] |
| tidyverse | 2.0.0 | [38] | rstudioapi | 0.14 | [39] |
| knitr | 1.42 | [40] | circlize | 0.4.15 | [41] |
| cowplot | 1.1.1 | [42] | flexmix | 2.3-19 | [43] |
| lattice | 0.20-45 | [44] | ggrepel | 0.9.3 | [45] |
| patchwork | 1.1.2 | [46] | reshape2 | 1.4.4 | [47] |
| mclust | 6.0.0 | [46] | dplyr | 1.1.1 | [48] |
| lsa | 0.73.3 | [49] | SnowballC | 0.7.0 | [50] |
| qgraph | 1.9.4 | [51] | igraph | 1.4.1 | [52] |
| RColorBrewer | 1.1-3 | [53] | lemon | 0.4.6 | [54] |
| ggthemes | 4.2.4 | [55] | ggpubr | 0.6.0 | [56] |
| ComplexHeatmap | 2.12.1 | [57] | YAPSA | 1.22.0 | [58] |
| ggplot2 | 3.4.2 | [59] | doMC | 1.3.8 | [60] |
| iterators | 1.0.14 | [61] | foreach | 1.5.2 | [60] |
| GenomicRanges | 1.50.0 | [62] | ASCAT | 3.1.2 | [63,64] |
| IRanges | 2.32.0 | [62] | Polychrome | 1.5.1 | [65] |
| S4Vectors | 0.36.0 | [66] | NMF | 0.27 | [67] |

| | | | | | |
|---|---|---|---|---|---|
| data.table | 1.14.8 | [68] | bigmemory | 4.6.4 | [69] |
| Biobase | 2.58.0 | [21] | Cairo | 1.6.2 | [70] |
| biomaRt | 2.56.1 | [71] | corrplot | 0.92 | [72] |
| doParallel | 1.0.17 | [73] | parallel | 4.4.0 | [74] |
| nnls | 1.5 | [75] | synchronicity | 1.3.10 | [69] |
| BiocManager | 1.30.23 | [76] | devtools | 2.4.5 | [77] |
| quadprog | 1.5.8 | [78] | | | |

### Experimental data

#### Patient samples and cell lines

##### Patient sample cohort

Clinical data and samples from the 33 patients with *EGFR*-mutant non-small cell lung cancer (NSCLC) treated with front-line osimertinib for advanced disease included in the OSIRESP cohort (ESR-20-20709) were used in this study (**Figure S12**). Samples for whole exome sequencing or shallow whole genome sequencing were collected from a total of 68 patients. Following sample processing and quality control, a total of 35 patients were identified as suitable for fitting copy number profiles and producing reliable CIN signature composition. Low quality samples were defined as samples with low purity, with insufficient number of total reads (<15 reads per bin per chromosome copy), or with noisy copy number profiles. The high dropout rate was due to the use of whole biopsy samples for sequencing (without macrodissection) resulting in high non-tumour DNA content within the collected sample used for DNA extraction (see section "DNA extraction" for further details).

All study procedures were done under the approval from the 12 de Octubre Ethics Committee protocol #20/653. Patients provided written, informed consent for participation in this study.

Response Evaluation Criteria in Solid Tumors (RECIST) guidelines[79] were used to measure response to osimertinib. Overall survival (OS) was calculated from the treatment start date to the date of death or loss of follow-up. Progression-free survival (PFS) was computed from the treatment start date to the date of disease progression, death or loss of follow-up. Following clinical data curation, a final cohort of 33 patients was included in the downstream analyses (**Figure S12**).

##### Human cancer cell lines

The 4 *EGFR*-mutant NSCLC cell lines (HCC827, HCC4006, PC9 and H1975) were obtained from the American Type Culture Collection (Manassas, VA). All cell lines were maintained in Roswell Park Memorial Institute (RPMI) 1640 medium (GIBCO, Carlsbad, CA) with 10% fetal bovine serum (FBS), penicillin (100 U/mL), and streptomycin (50 mg/mL) (Complete Medium) in a humidified $CO_2$ incubator at 37°C. All cells were passaged for less than 3 months before being renewed with frozen, early-passage stocks. Cells were regularly screened for mycoplasma using a MycoAlert Mycoplasma Detection Kit (Lonza). Before performing the experiments, cell line identities were confirmed by STR profiling and *EGFR* mutation was confirmed by droplet digital PCR (ddPCR) using the following ddPCR™ Mutation Detection Assays (Bio-Rad Laboratories): ddPCR *EGFR* Exon 19 Deletions Screening Kit (12002392), *EGFR* p.L858R (dHsaMDS463105111), and *EGFR* p.T790M (dHsaMDS759157834). Parental cell lines were sequenced via shallow whole genome sequencing.

Resistant clones were generated by long-term culture of individual cell lines with high concentration of a EGFR tyrosine kinase inhibitor (EGFR-TKI), as previously described[80].

Briefly, a total of 1x10$^{6}$ cells were seeded into 10-cm dishes and allowed to adhere overnight. The following day, the medium was replaced with fresh complete medium containing 2 µM erlotinib for HCC827, 10 µM erlotinib for HCC4006 and PC9, or 0.7 µM osimertinib for H1975. After approximately 9 days, most cells perished, leaving behind a small number of isolated survivor cells. Clearly separated colonies were isolated and transferred to 96-well plates between 8 and 12 weeks of drug treatment. From there, colonies were expanded successively from 96-well plates to 24-well plates, then to 6-well plates, until finally reaching confluency in a 10-cm plate, thereby establishing individual resistant clones. During the whole process and after generation of resistant clones, medium was replaced every 2-3 days with fresh drug at the same concentration used initially. Canonical resistant mutations, such as T790M (dHsaMDS759157834) and C797S (dHsaMDS759157834), were assessed by ddPCR and tested negative for all resistant clones.

Short term HCC827 resistant cells were generated by incubating 1.5x10$^{5}$ cells, plated the day before on 6-well plates, with either 1 µM erlotinib or a combination of 1 µM erlotinib and 0.5 µM MET inhibitor (PHA-665752). After 1 month, cells were recovered by trypsinization and single cell sorting was performed using an image based piezoelectric nanoliter dispenser (cellenONE, Scenion). Single cells were selected based on cell line specific parameters of cell diameter, elongation and circularity. Single cells were dispensed into 96-well PCR plates containing pre-dispensed 300 nL of 30 mM Tris buffer pH=8, then stored at −20°C.

### DNA extraction

Formalin-fixed, paraffin-embedded (FFPE) tissue blocks were cut as 5 µm sections. Microdissection was not used for recovering tumour-enriched regions. DNA was extracted from 10-25 sections using QIAamp DNA FFPE Tissue Kit from Qiagen (Cat. No. 56404) following manufacturer’s instructions.

DNA extraction from cell pellets of about 5x10$^{6}$ cells per sample was performed using the DNeasy® Blood & Tissue Kit from Qiagen (Cat. No. 69504). Cell pellets were equilibrated at room temperature for 15 minutes to then follow manufacturer's guidelines. DNA was finally eluted in Tris-HCl (pH=8, 10 mM) and was quantified using the ADNds Quant-iT™ PicoGreen™ kit from Invitrogen™ (Cat. No. P7589).

### DNA sequencing

#### Shallow whole genome sequencing (sWGS)

For parental lung cancer cell lines, extracted DNA at 5 ng/µL was used. 50 µL of sample was dispensed into a Covaris microTUBE AFA Fiber Pre-Slit Snap-Cap 6x16mm (Covaris, Cat. No. 520045) and DNA was mechanically fragmented for 150 seconds using the E220evolution focused-ultrasonicator from Covaris.

Due to the different quality among the clinical samples, different DNA inputs were manipulated, therefore samples were divided in groups of 1, 2 and 5 ng/µL. Samples with low volume availability were sheared using 10 µL in a microTUBE-15 AFA Beads H-Slit Strip V2 (Covaris, Cat. No. 520241) for 250 seconds, and the rest of them were sheared using 50 µL in a microTUBE AFA Fiber Pre-Slit Snap-Cap 6x16mm for 300 seconds in the ultrasonicator.

For both lung cancer cell lines and clinical samples, DNA fragmentation was aimed to obtain DNA fragments of about 200 bp. DNA profiles were checked using the LabChip DNA High Sensitivity Reagent kit (PerkinElmer, Cat. No. CLS760672) on the LabChip GX Touch Nucleic Acid Analyzer, and the Agilent High Sensitivity DNA Kit (Agilent, Cat. No. 5067-4626) on the 2100 Bioanalyzer Instrument.

For library preparation, ThruPLEX® DNA-Seq Kit (Takara, Cat. No. R400674) was executed following manufacturer's instructions. DNA index kits belonged to the same manufacturer and were compatible with Illumina sequencing instruments.

Library cleanup was carried out with Agencourt AMPure XP Reagent (Beckman Coulter, Cat. No. A63881) as recommended. Final quality control of libraries was assessed via sample quantification using the ADNds Quant-iT™ PicoGreen™ kit, and via library profiles using the LabChip DNA High Sensitivity Reagent kit.

Libraries were then pooled together in equal ratios and sequenced by the Genomics and CEGEN Units of the Spanish National Cancer Research Centre (CNIO) with the NextSeq™ 550 or the NovaSeq™ X Plus systems from Illumina aiming for 80 million reads per tumour sample and 10 million reads per parental cell line.

#### Whole exome sequencing (WES)

For sample preparation, a total of 200 ng of DNA was required. WES was conducted by a commercial service (Macrogen Inc., Seoul, Korea). Libraries were constructed using the SureSelect Human All Exon v6 (Agilent Technologies, Santa Clara, CA, USA). Sequencing of pair-end reads of 151 bp length was performed with the Novaseq™ X Plus system from Illumina, obtaining an average total reads across samples of 350 million reads (range 98-510 million reads).

#### Single-cell whole genome sequencing (scWGS)

Single-cell DNA whole genome sequence libraries were prepared using SMARTer PicoPLEX Gold Single Cell DNA-Seq Kit (PicoPlex Gold, Takara) following the manufacturer’s protocol. Cells were subjected to an enzymatic and heat lysis for efficient release of genomic DNA. Then, a pre-amplification step of the genomic material was performed, followed by a clean up step for removing the excess of pre-amplification primers. Finally, an amplification step was conducted for exponential amplification of the DNA libraries and Illumina-compatible indexing. Libraries quality and quantity were assessed with DNAds Quant-iT™ PicoGreen™ kit on FLUOROSKAN (ThermoFisher Scientific) and LabChip DNA High Sensitivity Reagent kit on LabChip GX Touch Nucleic Acid Analyzer (PerkinElmer) according to the supplier’s recommendations. Libraries were then pooled together in equal ratios and sequenced by the Genomics Unit of the Spanish National Cancer Research Centre (CNIO) using NextSeq™ 550 system from Illumina aiming for 1 million reads per cell.

#### Read alignment

Reads were aligned as single-end against the human genome assembly GRCh37 using BWA-MEM (v0.7.17)[81]. Duplicate reads were then identified and marked using samtools-markdup (v1.15)[82].

### Generating copy number profiles

After alignment, we fitted absolute copy number profiles from sequencing data generated across the different high-throughput technologies. Following absolute copy number fitting, samples were rated using a star system as previously described[83,84], and those with 1-star were discarded for downstream analyses.

#### Absolute copy number fitting from sWGS

We used the QDNAseq R package[17] to count reads within 30 kb bins (the resolution threshold determined to accurately call copy number signatures[3]). Bins mapped to centromeres and regions of undefined sequence in the reference genome hg19 were excluded. Read counts were initially corrected for the relationship between sequence mappability and GC content by ignoring the sex chromosomes (default procedure of the QDNAseq R package). Then, we performed a second correction step including the sex chromosomes following developer's recommendations. After the profile was segmented, we inferred absolute copy numbers for a combinations of purity and ploidy values (0.05<purity<1, and 1.5<ploidy<8) as follows:

$$cn = \frac{1}{purity} \times \left(\frac{r}{d} - 2 * (1 - purity)\right)$$

, where *purity* is the fraction of tumour cells in the sample, *r* is the read counts, and *d* is a constant proportional to the read depth and the average absolute copy number of the tumour and normal cells in the sample:

$$d = \frac{r}{(ploidy \times purity + 2 \times (1 - purity))}$$

The optimal mathematical solution was considered the one with the lowest root mean squared deviation (RMSD) between the non-rounded and the rounded copy numbers for all bins. All solutions were manually inspected to either confirm or correct for more appropriate fits.

For pure cancer models (i.e. cell lines and single cells), we used a range of purities from 0.9 to 1 for fitting absolute copy numbers.

#### Absolute copy number fitting from WES

Paired-end raw reads were aligned to the human hg19 reference genome. The resulting alignments were split into equally-sized bins of 50 kb in size. Bins were annotated with replication timing and a set of metadata inherited from QDNAseq: gc, mappability, residuals, bases and blacklist. The annotated bins were then interrogated for overlaps with target regions (bed file) and divided in two groups: 1) off-target bins, with a zero overlap with targeted regions. In this case, reads were counted normally in the entire bin, and 2) on-target bins, those with at least 1 overlap with a target region. In this case every overlap was deleted and the remaining off-target regions were pasted together forming a new, smaller bin, ready for read counting. GC correction was performed using LOESS. To correct for artificially high off-target read counts in parts of the genome with high sequence similarity to the target regions, we generated a score per bin that quantified the magnitude of this bias and used it for a single LOESS fit and correction. We segmented these data using a modified version of the circular binary segmentation (CBS) from the DNAcopy R package[85]. Absolute copy numbers were inferred as described above in the section "Absolute copy number fitting from sWGS".

#### Absolute copy number fitting from scWGS

To infer copy number profiles from single-cell DNA sequencing data, we defined 100-200 kb windows along the genome and applied the same methodology described in the "Absolute copy number from sWGS" section.

#### Absolute copy number fitting from pseudobulk WGS

We reconstructed bulk copy number profiles by concatenating reads of all single cells sequenced from a specific sample. We downsampled pseudobulk bams to appropriate read counts based on the bin size (30 kb), purity and ploidy (30 number of reads per bin per chromosome copy). Absolute copy number fitting was performed as described in the "Inferring absolute copy number from sWGS" section.

### Detection of focal amplifications

Focal amplifications in unpaired WGS, WES and SNP6 data were defined as genomic regions with absolute copy number higher or equal to 8. This threshold was selected after an extensive analysis using a range of threshold values (see section "Assessing the threshold for defining focal amplifications"), and is in line with thresholds previously used in the literature[6]. For sex chromosomes, we adjusted the absolute copy number value for defining focal amplifications to 6, considering that males possess only one copy of each sex chromosome.

In longitudinal analyses, the acquisition of a new focal amplification in the late/relapse/metastasis biopsies was defined as the gain of at least 6 copies of an autosomal genomic region with respect to the early time point sample. In male patients, for oncogenes located in sex chromosomes (i.e. *AR*), we defined the acquisition of amplification as the gain of at least 3 copies compared to an early time point.

### Detection of homozygous deletions

Homozygous deletions in unpaired WGS, WES and SNP6 data were defined as genomic regions with absolute copy number lower than 0.5.

### Forecasting driver gene amplifications and deletions

Here, we present a forecasting framework that takes as input a tumour copy number profile, and then outputs whether the tumour has a high or low risk of acquiring the amplification or deletion of the given driver gene(s) at some (undetermined) point in the future (**Figure 1**).

This framework is designed to integrate the major forces governing the emergence and expansion of oncogene amplifications or tumour suppressor gene (TSG) deletions within a tumour cell population: (1) the chance of acquiring a specific alteration in the genomic region where the driver gene is located (*locus-specific mutation rate*, $\mu$), and (2) the selective advantage conferred by the driver amplification/deletion in the cell that acquires it (*selection coefficient, s*). Analysis of single cell DNA sequence data and simulations support the feasibility of this approach (**Supplementary Note 3**).

Our framework outputs *P(D)*, the probability of acquiring an alteration of a given driver gene (*D*) in the future. This is computed via the product of two terms:

$$P(D) = \mu \times s = \left( \sum_{CX=1}^{9} I_{CX} \times w_{D,t,CX} \right) \times f_{D,t}$$

, where the $\mu_{D,t}$ represents the steady-state probability of the mutation occurring at the locus of interest in a single cell within the tumour population, and the $s_{D,t}$ represents the selective advantage conferred by the amplification or deletion of the driver of interest. The *locus-specific mutation rate* ($\mu_{D,t}$) is derived by combining estimates of the mutational processes present in the input tumour genome $I_{CX}$ (details of the computation can be found in the section "Quantifying tumour mutational processes activity") with estimates of the mutational processes that can cause the amplification or deletion of the driver gene(s) of interest (*D*) in a given tumour type (*t*) denoted as $\omega_{D,t,CX}$ (computation details described in the section "Estimating locus-specific mutational processes"). The *selection coefficient* ($s_{D,t}$) is approximated by the frequency of the driver alteration at a cohort level $f_{D,t}$ (computation described in the section "Estimating selection coefficients").

If the tumour type (*t*) is not specified at input, a pan-cancer probability can be computed by averaging the *P(D)* for all tumour types in which the amplification or deletion of the gene of interest (*D*) is putatively under positive selection (see section "Estimating selection coefficient"). For each tumour type (*t*), we first multiply the locus-specific mutation rate ($\mu_{D,t}$) by the driver-specific cohort frequency ($f_{D,t}$). Then, the oncogene-specific, tumour-specific probabilities are averaged across all tumour types. This approach allows us to assign a greater contribution to the pan-cancer amplification risk from tumour types where the driver alteration provides a stronger positive selection:

$$P(D) = \left\langle \left( \sum_{CX=1}^{9} I_{CX} \times w_{D,t,CX} \right) \times f_{D,t} \right\rangle_t$$

, where $<...>_t$ indicates the average over all tumour types in which the driver alteration is under positive selection.

Since the homozygous deletion of a tumor suppressor gene typically involves two independent genomic events (see **Supplementary Note 4**), the risk of acquiring such a deletion decreases as the copy number state of the TSG increases. To account for this, we adjust the deletion probability by dividing it by the current copy number of the TSG in the input tumor genome.

*P(D)* can be subjected to a predetermined threshold to generate a final binarised prediction that forecasts the risk of acquiring an oncogene amplification and/or a TSG deletion as either high or low (**Figure 1b**). How the threshold is determined is described in **Supplementary Note 2**. For oncogene amplifications, tumour samples with a *P(D)* higher or equal than the threshold are classified as having high risk of acquiring the amplification of the given oncogene. In contrast, samples with a *P(D)* below the threshold or without detectable CIN (< 20 CNAs) are classified as having low amplification risk. For tumor suppressor gene homozygous deletions, samples with a *P(D)* above the threshold and ≥ 15 CNAs (the minimum required for accurately quantifying sample-level activities, **Figure S13**) are classified as having high risk of acquiring a homozygous TSG deletion. Samples not meeting this criteria are classified as having low risk.

Given that each driver gene has distinct prior probabilities of being altered and selected depending on the disease context, an appropriate setting is needed for learning the framework parameters and determining the optimal threshold for each driver gene and cohort. The modular structure of our forecasting framework provides flexibility, allowing users to tailor it to align with specific clinical and/or biological research settings (see **Supplementary Note 2**).

#### Estimating locus-specific mutation rates (μ)

Here, we describe how we estimate the probability that a focal amplification or homozygous deletion will occur at the gene of interest in a single cell within the tumour population (**Figure 1a**). To achieve this goal, we assume that the locus-specific mutation rate depends on the mutational processes operating in a tumour sample and their ability to cause focal amplifications or deletions specifically at the genomic region where the driver gene is located:

$$\mu_{D,t} = \sum_{CX=1}^{9} I_{CX} \times w_{D,t,CX}$$

, where $I_{CX}$ represents the activities of each signature ($CX$) in a given input tumour sample (see section "Quantifying tumour mutational processes activity" for details on how this is estimated), and $\omega_{D,t,CX}$ represents the weights of each signature ($CX$) at the genomic location harbouring the driver gene ($D$) for a given tumour type ($t$) (see section "Estimating local-specific causal mutational processes" for details on how this is estimated). Note: a min-max normalization is used to scale the locus-specific mutation rate ($\mu_{D,t}$) of a given sample to a reference cohort to bound the value between 0 and 1. It is recommended to use the same reference cohort that is used to determine the optimal threshold (**Supplementary Note 2**).

#### Quantifying tumour mutational processes activity ($I_{CX}$)

$I_{CX}$ reflects the activity levels of different mutational processes causing the accumulation of numerical and structural DNA changes across an input tumour sample. To quantify the activity of mutational processes from the input tumour copy number profile, we use our previous CIN signature approach[3] modified to a 3-feature signature encoding space (see below for details). We first compute a sample-by-component sum-of-posterior vector from the input tumour copy number profile. This is done by summing up the posterior probability 1x41 component vector for all copy number events of the sample. Then, we apply the linear combination decomposition function from YAPSA[58] to the input vector to derive a sample-by-signatures vector representing the activity of different types of mutational processes. To ensure robustness of small signature activities, we identify signature-specific thresholds, following the same procedure as in our previous work[3]. Then, we set activities to zero if they are below the signature-specific threshold. The sample-by-activity vector is finally normalised to sum to 1.

#### Estimating locus-specific mutational processes ($\omega_{D,CX}$)

To estimate $\omega_{D,t,CX}$, representing the mutational processes ($CX$) that can cause specific oncogene amplifications or tumour suppressor gene homozygous deletions for a given tumour type ($t$), we use a collection of examples of amplifications and deletions from the TCGA. We first estimate the putative causal mutational process(es) responsible for generating focal alterations where the driver gene(s) is located, $Z_{D,t,CX}$ (see section "Estimating causal mutational processes" for details). Then, we learn topological constraints that might bias this estimation, $X_{D,t,CX}$ (see section "Estimating topological constraints" for details). Finally, the contribution of each signature to the amplification or deletion of the driver gene(s) is calculated by correcting $Z_{D,t,CX}$ by $X_{D,t,CX}$:

$$w_{D,t,CX} = \frac{Z_{D,t,CX}}{X_{D,t,CX}}$$

Note: as TSG homozygous deletions occur early in tumour evolution (see **Supplementary Note 4** for further discussion), we base our estimation on all samples with ≥15 copy number alterations and reliable signature quantification. In the case of oncogene amplifications, we use all samples with detectable CIN (≥20 copy number alterations) according to the threshold estimated in our previous work[3].

#### Estimating causal mutational processes ($Z_{D,CX}$)

To estimate $Z_{D,t,CX}$, the putative causal CIN signatures for each oncogene amplification or tumour suppressor homozygous deletion, we had to modify our original CIN signature encoding as it did not facilitate mapping of signatures to individual copy number events[3]. Thus, we designed a new feature encoding space that could enable mapping to individual events while maintaining the ability to quantify the original CIN signature compendium. This new encoding consists of three features: the segment size, represented by 22 components; the difference in copy number between adjacent segments, represented by 10 components, and the breakpoints per 5 Mb side, represented by 9 components. With this new encoding, we are able to compute the probability that a given event belongs to any one of the feature components, represented by a 41-component probability vector. Using this approach, we can ultimately assign putative causal signatures to amplified or deleted segments harbouring a given oncogene or tumour suppressor gene.

In the case of oncogenes, we combine probability vectors from all segments spanning oncogene amplifications resulting in an oncogene-by-component probabilities matrix, which is then normalised per feature. We then average feature component probabilities to compute an oncogene-specific vector of feature component probabilities per tumour type. All oncogene-specific vectors are then combined to generate the "oncogene by copy number feature" matrix **Φ** (with elements $\phi_{OG,c}$) per tumour type (**Figure S15**).

For tumour suppressor genes (TSG), we adopt the same procedure to generate a "TSG by copy number feature" matrix **Φ** (with elements $\phi_{TSG,c}$) per tumour type (**Figure S15**), but with one modification to the changepoint feature calculation: given that homozygous deletions may originate from an initial single-copy loss and signals may be disrupted by subsequent whole-genome doubling (WGD) events in CIN samples (see **Supplementary Note 4** for further discussion, and **Figures S15-16**), we correct the probabilities assigned to the changepoint feature in homozygous deleted events: for 0-copy segments, we assign a weight of 1 to the changepoint feature component representing 1-copy change, while a weight of 0 is assigned to the other changepoint components.

To define a signature background for enrichment calculations, we follow a similar approach to the oncogene setting, but use all copy number events (*p*) in the tumour-specific TCGA cohort (*t*) to generate a "passenger event by component probabilities" matrix **Φ** (with elements $\phi_{p,c}$) per tumour type. For simplicity, in the following description of the method, we use $\phi_{e,c}$ as a general notation for $\phi_{OG,c}$ , $\phi_{TSG,c}$, and $\phi_{p,c}$, while allowing substitution with the corresponding term whenever appropriate.

To compute putative causal signatures, we multiply the "event by copy number features" matrix (**Φ**) and the "CIN signature definition map" matrix (**σ**). The "CIN signature definition map" matrix (with elements $\sigma_{c,CX}$) refers to the definitions of the CIN signatures (*CX*) in the 3-feature components (*c*) space. For each sample (*s*), the resulting event-by-signature matrix is then multiplied by the sample-level signature activities $I_{CX}^{(s)}$ (see section "Quantifying tumour mutational processes activity") to compute activity-corrected signature scores for each copy number event (*e*), which is finally normalised per signature (*CX*). Activity-corrected event-by-signature scores of all samples are finally joined into a single matrix **ε**:

$$\varepsilon_{e,CX} = \frac{I_{CX}^{(s)}[\mathbf{\Phi}^{(s)} \cdot \boldsymbol{\sigma}]_{e,CX}}{\sum_{CX'=1}^{9} I_{CX'}^{(s)}[\mathbf{\Phi}^{(s)} \cdot \boldsymbol{\sigma}]_{e,CX'}}$$

Given a driver gene (*D*), we then calculate the relative contribution of a CIN signature to the focal alteration for a given tumour type ($q_{D,t,CX}$) by averaging the activity-corrected CIN signature score of all altered events spanning a given driver gene in a specific tumour type as follows:

$$q_{D,t,CX} = \frac{\sum_{a=1}^{A_{D,t}} \varepsilon_{a,CX}}{A_{D,t}}$$

, where $\varepsilon_{a,CX}$ represents the activity-corrected score of one of the CIN signatures (*CX*) for a particular altered event (*a*), and $A_{D,t}$ represents the number of altered segments spanning a particular driver gene (*D*) in a given tumour type (*t*). Only universal CIN signatures are included in this calculation (see section "Assessing stability across technologies").

Next, we compute the relative contribution of a CIN signature to the acquisition of any copy number event in a specific tumour type ($q_{t,CX}$) by averaging the activity-corrected CIN signature scores of all copy number events found in the tumour-specific training cohort as follows:

$$q_{t,CX} = \frac{\sum_{p=1}^{E_t} \varepsilon_{p,CX}}{E_t}$$

, where $\varepsilon_{p,CX}$ represents the activity-corrected score of one of the CIN signatures (*CX*) for a particular copy number event (*p*), and $E_t$ represents the total number of copy number events observed in the training cohort of a given tumour type (*t*).

Finally, we generate the "causal mutational processes" term by computing the tumour-specific driver-specific CIN signature weights ($Z_{D,t,CX}$). These weights are derived by normalising the relative contribution of a CIN signature to the driver alteration in a given tumour type ($q_{D,t,CX}$) to the relative contribution of a CIN signature to the acquisition of any copy number event in any tumour type ($q_{t,CX}$) as follows:

$$Z_{D,t,CX} = \frac{q_{D,t,CX}}{q_{t,CX}}$$

, where $Z_{D,t,CX}$ is finally normalised so that the sum of all signature weights (*CX*) equals 1.

This procedure applied to the TCGA, for each tumour type, allowed us to construct two "causal mutational processes" matrices containing: 1) the weights of the 9 universal CIN signatures for 271 oncogenes ($Z_{OG,t,CX}$), and 2) the weights of the 9 universal CIN signatures for 306 tumour suppressor genes ($Z_{TSG,t,CX}$). See **Supplementary Note 5** for an analysis of the biological implications of these matrices.

##### Estimating topological constraints ($X_{D,CX}$)

The size or amplitude of a copy number alteration caused by a mutational process in a particular genomic location may be influenced by the local chromatin structure[86,87]; or by selective constraints, for example the presence of neighbouring genes to which the extension of the copy number alterations may be beneficial or detrimental[88,89]. This may bias our estimates of $Z_{D,CX}$ causing us to assign the wrong causal mutational processes. Thus we need to correct for any topological constraints. Through a comprehensive analysis of the landscape of focal amplifications and homozygous deletions across multiple samples, we hypothesise we can infer how these alterations are constrained by DNA topology and estimate a correction factor.

To do this, we use our 3-feature copy number encoding to observe biases in the size, changepoint and breakpoint density of driver alterations over a null distribution estimated from non-driver or "passenger" alterations (described in detail below). Note: given that homozygous genome loss is commonly deleterious and thus heavily negatively selected[90], we are not able to account for local topological constraints for TSG homozygous deletions (see **Supplementary Note 4** for further discussion).

For each tumour type (*t*) and oncogene (*OG*), the null distribution is estimated by dissecting configurations of non-oncogene ("passenger") amplifications in tumor samples harboring amplification of the given oncogene. Specifically, we first calculate the posterior probability of each passenger amplified segment belonging to each feature component, resulting in a 1x41 posterior probability component vector. These component vectors from all passenger amplified events were then averaged to compute a passenger-by-component probability vector $\beta_{OG,t,c}$. We repeat this procedure to generate a similar oncogene-by-component vector but with the driver events, $\alpha_{OG,t,c}$. **Figure S14** summarises the configurations of amplified segments harbouring a specific oncogene in the TCGA cohort.

We then generated a "topological constraints" term by dividing the regional biases constraining the configuration of oncogene amplifications ($\alpha_{OG,t,c}$) by the regional biases constraining the configuration of passenger amplifications ($\beta_{OG,t,c}$). To integrate this term into the forecasting framework, we expressed the matrices in the signature space by multiplying with the signature-by-component definition matrix ($\boldsymbol{\sigma}$) as follows:

$$X_{OG,t,CX} = \frac{[\boldsymbol{\alpha} \cdot \boldsymbol{\sigma}]_{OG,t,CX}}{[\boldsymbol{\beta} \cdot \boldsymbol{\sigma}]_{OG,t,CX}}$$

Since CX1 is associated with mitotic errors and is likely unaffected by local chromatin structure, we assigned a weight of 1 to CX1 in the resulting oncogene-by-signature matrix and then renormalized per oncogene to sum to 1.

#### Estimating selection coefficients (s)

Here, we describe how we estimate the selective advantage an oncogene amplification or a TSG homozygous deletion confers to the cell that acquires it (**Figure 1a**). This estimation relies on previous literature[89,91–93] indicating that, for driver amplifications and deletions showing evidence of positive selection in a specific tumour type, the tumour-specific frequency computed from large-scale cohorts (which may have variable mutation rates) serves as a reasonable proxy for *s* (see **Supplementary Note 1** for further discussion).

We initially created a list of oncogenes and TSGs by integrating and curating the annotations provided by Bailey *et al*. 2018[12], Jassal *et al*. 2020[13], Martínez-Jiménez *et al*. 2020[14], and Sondka *et al*. 2018[15]. This yielded a total of 271 oncogenes and 306 TSGs. For each tumour type, we selected the oncogenes and TSGs that were focally amplified and deleted, respectively, in at least 5 samples of the TCGA tumour-specific cohort. This resulted in a total of 929 oncogene/tumour type pairs, including 22 different tumour types and 188 oncogenes; and in a total of 54 TSG/tumour type pairs, including 22 different tumour types and 16 tumour suppressor genes. Then, we confirmed that the amplification or deletion of those driver genes showed evidence of a selective advantage in the tumour type by leveraging results obtained by GISTIC2.0[16] applied to 10,844 tumours from the TCGA cohort (https://portals.broadinstitute.org/tcga/home, data version: "2015-06-01 stddata__2015_04_02 regular peel-off"). From our list of driver genes/tumour type pairs, we only retained those driver genes located in an amplification or deletion peak that were significantly altered in a specific tumour type (q-value ≤ 0.25). This resulted in a total of 257 oncogene/tumour type pairs where the amplification may be under positive selection (**Figure S1**), and a total of 43 TSG/tumour type pairs where the homozygous deletion may be under positive selection (**Figure S2**).

These data allowed us to construct a "cohort frequency" matrix (*f*) by computing the frequency of samples carrying the alteration of a given driver gene per tumour type, as follows:

$$f_{D,t} = \frac{A_{D,t}}{S_t}$$

, where $f_{D,t}$ represents the frequency of carriers of a given driver (*D*) alteration for a tumour type (*t*), $A_{D,t}$ is the number of samples with the alteration of a given driver gene *(D)* and tumour type *(t)*, and $S_t$ represents the total number of samples for a given tumour type (t).

This approach applied to TCGA generated two "driver-specific cohort frequency" matrices: one for oncogenes ($f_{OG,t}$), a 271 oncogenes by 33 tumor types matrix, and another for tumor suppressor genes ($f_{TSG,t}$), a 306 TSGs by 33 tumor types matrix.

Frequencies of driver genes not included in the oncogene/tumour type and TSG/tumour type combinations were set to zero in the "driver-specific cohort frequency" matrices.

### Robustness analyses

#### Assessing stability across technologies

CIN signatures were derived using TCGA SNP 6.0 array data because this represents the largest pan-cancer collection of high-resolution copy number profiled tumours to date. In our previous work, we evaluated the robustness of our approach across different high-throughput sequencing technologies by comparing signature definitions and activities. As expected, we saw a drop in similarity scores for the activities and signature definitions between SNP6 and sWGS data[3].

Here, we extended this analysis by means of testing the ability to capture our compendium of signatures in sWGS, which is the technology used for obtaining copy number profiles from the experimental data included in this study (see section "Experimental data"). For this analysis, we used 478 TCGA samples with detectable levels of CIN that were also part of the PCAWG project. WGS data from the 478 TCGA/PCAWG samples was downsampled to 15 number of reads per bin per chromosome copy (according to the ploidy, purity and bin size) for mimicking sWGS. We fitted absolute copy number profiles (see section "Absolute copy number fittings from sWGS"), extracted the 5 fundamental copy number features but using the new changepoint encoding (see companion work), quantified activities for the 16 compendium signatures (CX6 excluded) using YAPSA[58], and compared them to the SNP6-derived signature activities using Kendall's tau.

This analysis showed the deterioration of some signatures across technologies (**Figure S16**). Signatures showing robust signals across technologies were classified as universal signatures (CX1-5, CX8-10, and CX13) whereas others were classified as SNP6-specific signatures (CX7, CX11-12, and CX14-17). CX14 was not considered universal because its signal was captured by CX1 in sWGS data. It is important to note here that SNP6-specific signatures can also be detected in sWGS, but the copy number changes captured by the signatures manifest differently across the two technologies.

To test the robustness of the forecasting framework across sequencing technologies, we then computed amplification scores in the SNP6- and sWGS-derived copy number profiles from the 478 samples included in both the TCGA and the PCAWG project. Oncogene-specific amplification scores were computed using a pan-cancer level procedure, and then averaged per sample for comparisons. We used a Pearson correlation to compare sample pairs and showed a high correlation of scores obtained from different technologies ($r = 0.85$, p-value < 2.2e-16; **Figure S17**).

#### Assessing stability across bin resolutions

Given that the gold-standard bin size for quantifying our compendium of CIN signatures is 30 kb[3], and the bin size used for deriving copy number profiles from WES was 50 kb (see section "Absolute copy number fitting from WES"), we explored the robustness of our CIN signatures across bin resolutions by comparing activities.

Here, we segmented copy number profiles from the 478 TCGA/PCAWG samples using 8 different bin sizes (30, 40, 50, 60, 70, 80, 90 and 100 kb). Once absolute copy number profiles were fitted, we quantified the CIN signatures using the 3-feature encoding per sample. We then computed cosine similarities of SNP6-derived signature activities with those obtained at each bin resolution (**Figure S18a**). In addition, we compared signature activities derived at 30 kb (gold-standard) and 50 kb (resolution used for WES data) using Kendall's tau (**Figure S18b**). Altogether, these comparisons indicate our novel signatures are stable across bin resolutions.

#### Robustness of using the universal set of signatures

The use of a signature subset for decoding mutational processes operating in tumours could erroneously attribute alterations caused by unrepresented signatures to the signature set. This presents a fundamental challenge in the mutational signatures field, potentially affecting our ability to predict amplifications driven by these unrepresented signatures. Previous attempts to mitigate this effect have involved *de novo* signature discovery within the cohort of interest and subsequently mapping them back to a compendium, which may identify unmapped signatures. However, this approach is only feasible when there is a sufficiently large cohort for *de novo* discovery. Unfortunately, this condition is not met for many of our cohorts with oncogene amplifications. Therefore, in order to strike a balance and facilitate technology-agnostic forecasting, we decided to exclusively use the universal signatures.

To evaluate the extent of copy number events that may be erroneously assigned to the set of universal signatures, we applied our mapped approach using the whole set of CIN signatures and allocated each copy number event to the signature with the highest associated probability. Subsequently, we estimated the rate of events assigned to a CIN signature not included in the universal set. This analysis revealed a misassignment rate of 22.5% (145,554 out of 647,630 events), with most of them (17%) assigned to a signature linked to subclonal changes with unknown aetiology (CX7). The average misassignment rate per sample was 23.3 ± 14.4%.

To further warrant the use of universal signatures to build our forecasting framework, we explored the impact of the presence of these misassigned events within copy number profiles on the signature quantification task. We compared activities of universal signatures quantified in TCGA profiles including and excluding copy number events not mapped to the universal signature set using both Pearson's correlation (average Pearson's r across signatures of 0.84; **Methods Table 3**) and cosine similarities (average cosine similarity across signatures of 0.91; **Methods Table 3**).

**Methods Table 3. Concordance of sample-level activities after excluding misassigned events**

| CIN signature | Pearson's r | Cosine similarity |
|---|---|---|
| CX1 | 0.86 | 0.92 |
| CX2 | 0.73 | 0.90 |
| CX3 | 0.83 | 0.97 |
| CX4 | 0.94 | 0.96 |
| CX5 | 0.80 | 0.88 |
| CX8 | 0.91 | 0.92 |
| CX9 | 0.72 | 0.79 |
| CX10 | 0.85 | 0.92 |
| CX13 | 0.92 | 0.92 |

In addition, we tested whether quantifying the universal set of CIN signatures was sufficient for predicting amplifications. To do this, we performed a Pearson's correlation to compare amplification scores computed by including all 16 signatures (CX6 was excluded since it is not captured with the new feature encoding) and those obtained using only the 9 universal signatures. Oncogene-specific amplification scores were computed using a pan-cancer level procedure, and then averaged per sample for comparisons. Results demonstrated a high correlation between the scores obtained with both signature sets in the TCGA dataset ($r = 0.86$, p-value < 2.2e-16; **Figure S19**).

Finally, as the mutational processes active in a specific cohort may be different from those captured by the 9 universal signatures, we also conducted a comparison between the amplification scores computed using the universal signature set to those derived from a cohort-specific set of signatures. To undertake this analysis, we leveraged the TCGA-BRCA cohort, which comprises the largest sample size within the TCGA dataset (n=730), as the *de novo* signature extraction process requires a substantial cohort. This extraction was performed using non-negative matrix factorisation (NMF) on the input matrix as detailed in our previous work[3] (code available here: https://github.com/markowetzlab/CINSignatureDiscovery). A total of 6 breast-specific signatures were identified, all of which were mapped to one of the signatures included in our compendium of CIN signatures (CX2, CX3, CX4, CX7, CX9, and CX14). We averaged the oncogene-specific amplification scores per sample for comparisons. Results demonstrated a high correlation between the scores obtained using both signature sets in the TCGA-BRCA dataset ($r = 0.87$, p-value < 2.2e-16; **Figure S20**).

Altogether, our findings revealed the putatively low impact of applying a fixed signature repertoire for sample-level quantification and for forecasting focal amplifications.

#### Assessing signature assignment to individual events

Our approach for mapping CIN signatures to individual copy number events outputs a score that reflects the likelihood of an event being attributed to each signature. This flexibility accommodates situations where copy number events exhibit features resembling more than one signature, and also avoids assigning a specific CIN type to a copy number event which has been produced by an uncertain or unrepresented mutational process.

In an attempt to assess the reliability of our mapping approach, we allocated each copy number event to the signature with the highest associated probability. Subsequently, we assessed the accuracy of these assignments by calculating the fraction of events displaying similar features to the definition of the signature assigned. The signature with the highest cosine similarity in comparing the event-by-component vector and the signature-by-component vector was considered as the most similar. This analysis revealed an overall accuracy of 72.2% (479,565 out of 647,630 events), denoting a substantial alignment between the mapped signature and features defining an individual copy number event. **Methods Table 4** summarises mapping accuracy for each CIN signature.

**Methods Table 4. Fraction of events correctly assigned to the CIN signature with the most similar definition.**

| CIN signature | Events correctly assigned (%) |
|---|---|
| CX1 | 52.3 |
| CX2 | 75.8 |

| | |
|---|---|
| CX3 | 76.8 |
| CX4 | 78.1 |
| CX5 | 100 |
| CX8 | 59.3 |
| CX9 | 60.5 |
| CX10 | 89.0 |
| CX13 | 57.9 |

#### Assessing number of individual events required for sample-level signature quantification

We performed an analysis to determine the minimum number of copy number events required within a sample to accurately replicate its signature composition. To achieve this, we progressively increased the number of events of a sample profile via random selection. We then quantified sample-level signature activities by using the linear combination decomposition function from YAPSA on the input matrix (sample-by-component sum-of-posterior matrix). In each iteration of this decomposition, we expanded the information within the input matrix by progressively introducing an increasing number of events, ranging from 1 to 100. We then evaluated whether a sample's signature composition (defined by its three dominant signatures) was correctly captured depending on the number of events. This procedure was repeated 1,000 times with 1,000 TCGA samples being randomly selected for each run. A minimum of 15 copy number events were needed to correctly decompose >90% cases (**Figure S13**).

This procedure was also expanded to encompass individual signatures. For each specific signature, we performed random selection of copy number events from the subset of 200 TCGA samples exhibiting the highest signature activity. Then, we quantified sample-level signature activities and evaluated the success rate of detecting the signature. This procedure was done 100 times per signature. Our findings indicated that an average of 7 copy number events was sufficient for calling an individual signature with >90% of accuracy (**Figure S21**).

#### Assessing the threshold for defining focal amplifications

To select the optimal threshold for defining focal amplifications, we conducted an extensive analysis of our predictor's performance across a range of thresholds following two strategies: 1) we used absolute copy number values (ranging from ≥ 6 to ≥ 10 copies); and 2) we used copy number values relative to tumour ploidy (ranging from ≥ 4 to ≥ 8 copies respect to tumour's ploidy). For each threshold value, we computed the DNA topology effects at pan-cancer level and incorporated them in the framework for forecasting the amplification of a given oncogene. We then assessed the model performance by applying it back to the training cohort using a retrospective strategy, where signature activities were quantified after removing amplified segments harbouring the specific oncogene. Only samples from tumour types in which the oncogene putatively confers a fitness advantage were used for testing model performance. Performance was evaluated by comparing amplification scores between samples with and without oncogene amplifications using Wilcoxon two-sided test, and by computing the area under the receiver operating characteristic curve (AUC) (**Figure S22**). Although we observed marginal differences in performance across these different threshold values, higher thresholds tended to show improved performance. However, higher threshold values resulted in fewer amplifications available to train the model, limiting our ability to predict across as many oncogenes. We obtained similar positive performance using the ploidy-aware strategy, which indirectly may take into account whether a WGD event has occurred. We therefore chose a threshold of ≥ 8 as it represented a balance between performance and inclusivity, and was in line with the threshold previously used in the literature[6].

#### Assessing whether mutational processes can be detected in the absence of focal amplifications

We designed the forecasting framework with the aim of predicting future oncogene amplification at an early tumour evolution point. For that reason, here we tested the ability to capture DNA damage processes causing focal amplifications in the absence of these copy number events. To do so, we removed all copy number segments with a copy number value equal or higher than 8 in samples from the TCGA cohort. Then, we quantified the activity level of the 9 universal signatures at sample level as previously described (see section "Quantifying tumour mutational processes activity"). Results showed that CIN signatures associated with low-, medium- and high-level amplifications (CX8, CX9 and CX13, respectively) can be detected in a tumour even if the tumour does not yet have an amplification (**Figure S23**).

#### Assessing the impact of the genomic context on amplicon configurations

Previous research has demonstrated that breakpoints within a chromosome arm are enriched in regions with high densities of driver genes, suggesting that the segment size of copy number alterations may be influenced by the fitness effects of neighbouring driver genes[89]. To address this, we designed the forecasting framework to separate biases in amplicon configuration (topological constraints term, $X_{D,CX}$) from the underlying mutational processes driving oncogene amplification (local mutational processes term, $Z_{D,CX}$). Here, we evaluated whether the fitness effects of neighboring genes were effectively removed, thereby isolating the mutational process responsible for oncogene amplification. For each oncogene, we calculated the distance to surrounding TSGs located on the same chromosome, normalizing these distances by chromosome length. We then assessed correlations between the distance of the oncogene to the nearest TSG and both amplicon size and the number of flanking breakpoints (**Figure S24**). This analysis revealed no significant correlations, suggesting that the presence of neighbouring TSGs may not influence amplicon configurations.

#### Assessing the ability to capture mutation rates

To test the ability of our framework to infer oncogene amplification predisposition, we simulated 10 sample cohorts with 625 samples where mutation rates can be known with certainty. These simulated samples exhibited mutational processes driving copy number alterations (including focal amplifications) across 100 genomic regions (including the oncogene locus), which were influenced by specific cohort-level topological constraints.

For each of the 10 cohorts, we simulated 100 genomic regions and assumed that each tumour sample harboured exactly 1 copy number event per region. The first genomic region was designated as the oncogene locus (driver), while the remaining 99 were considered as passenger regions. Since copy number configurations may be biased by local topological features, we incorporated DNA topology features at each genomic region by generating a topological constraints matrix, $\mathbf{X}$. For each region $i$, we generated a 41-dimensional vector $X_i$:

$$X_i = [x_{i1}, x_{i2}, \ldots, x_{i41}]$$

, which defined the genomic region in the feature component encoding space. This encoding space consisted of the three fundamental copy number features: the segment size (22 components), the copy number change (10 components), and the breakpoints per side (9 components). We fitted three independent Gaussian distributions, one per feature, to generate the values in $X_i$. Specifically, for a given component $j_k$ from the feature $k$, its value was drawn from

$$g_{ij_k} \sim \mathcal{N}\left(\mu_{ik}, \sigma_k^2\right)$$

, where the standard deviation $\sigma_k$ was fixed at a sufficiently high level to ensure smoothed topological constraints, while the mean $\mu_{ik}$ was placed in a feature component that was randomly selected. Each feature-specific distribution was normalized to sum to 1, and the three normalized distributions were then concatenated to generate the 41-component vector per genomic region. As we defined $i = 1$ as the oncogene locus, the first row of $X_1$ defined the oncogene-specific topological constraints, while the subsequent rows represented passenger regions.

Once the cohort-specific topological constraints matrix $\mathbf{X}$ was established, we simulated 625 samples per cohort. First, we simulated the underlying mutational processes acting on a sample by modelling the activity of five signatures representing different types of chromosomal instability: CX1, mitotic errors; CX3, impaired homologous recombination; and CX8, CX9, and CX13, replication stress causing amplifications. For each sample, we generated a random signature activity vector $I_{CX}$ (with $\sum_{CX=1}^{5} I_{CX} = 1$). Next, for each of the 100 genomic regions, we randomly selected a signature causing an alteration based on the weights in $I_{CX}$, and multiply the corresponding signature definition vector by the region-specific topological constraints vector $X_i$. The resulting vector is renormalized so that components values of each feature sum to 1. Then, one component per feature is randomly sampled (based on the weights given by the vector) to determine the configuration of the copy number alteration in the region. This entire process is repeated to create $10n$ samples, where $n$ represents the desired final sample size (in practice, $n = 625$). To emulate varying cohort-level frequencies of driver amplification, we selected from these $10n$ samples a subset of $n \cdot f$ samples in which the oncogene region exhibits an amplification (defined as an event with at least five additional copies compared to the neighboring segment, i.e. changepoints components 8 to 10) and another subset of $n \cdot (1 - f)$ samples where no amplification occurred. This selection process was repeated for 18 distinct values of $f$, ranging from 0.01 to 0.5, reflecting the frequency range observed in TCGA.

To perform downstream independent testing analyses, we also generated a replicate for each of the 10 sample cohorts, maintaining the same topological constraints matrix while using a different random seed for the generation of sample-level signature activity vectors and copy number events.

Once simulations were performed, we next extracted the risk of acquiring an amplification in the oncogene locus for each sample. To achieve this, given the sample-level signature activity vector $I_{CX}$, we multiplied the signature-by-components definition matrix $\tilde{\sigma}$ (5 signatures x 41 components) and $I_{CX}$ to generate a sample-by-components vector $\mathbf{v}$:

$$\mathbf{v} = \tilde{\sigma} \cdot \mathbf{I}_{CX}$$

We then extracted the corresponding values for the changepoint feature components $\mathbf{v}_{\text{chp}}$, which was multiplied by the changepoint feature components of the topological constraints defining the genomic region harbouring the oncogene (denoted as $(X_1)_{\text{chp}}$). The product of these two vectors, normalized to sum 1, yields the probability distribution for each changepoint feature component $j$ sample:

$$p_j = \frac{(v_j)_{\text{chp}} \cdot (x_{1,j})_{\text{chp}}}{\sum_{j'} (v'_j)_{\text{chp}} \cdot (x_{1,j'})_{\text{chp}}}$$

The risk of acquiring an oncogene amplification for a given sample was defined as the sum of the probabilities of changepoint components 8 to 10, which correspond to focal amplifications (i.e. events showing a change of at least 5 copies respect to the neighboring segment). If $A$ represents these changepoint components, the acquisition risk of driver amplification for a given sample is calculated as:

$$P(D) = \sum_{j \in A} p_j$$

Finally, we inferred oncogene amplification risk using our forecasting framework. For each simulated cohort, we estimated the “local mutational processes” ($Z_{D,CX}$) and the “topological constraints” ($X_{D,CX}$) terms of our forecasting framework to then compute the “locus-specific mutational processes” term ($\omega_{D,CX}$), which was ultimately combined with the sample-level signature activities ($I_{CX}$) to infer the mutation rate driving amplifications in the oncogene region in a given simulated sample. The framework terms were learnt in the training cohorts and subsequently applied to both training and testing cohorts to evaluate performance. We observed strong correlations between the simulated and inferred risk of oncogene amplification acquisition, with higher correlation coefficients when topological constraints were taken into account for computing locus-specific mutational processes (Training sets: average Pearson's r across simulated cohorts of 0.794 with topological correction and 0.776 without correction; Testing sets: average Pearson's r across simulated cohorts of 0.796 with topological correction and 0.778 without correction; **Figure S25a**). We also evaluated how well the model captured topological constraints at the oncogene locus by computing cosine similarities between estimated and the simulated constraints (average cosine similarities of 0.926; **Figure S25b**). We finally assessed the ability to capture signatures causing oncogene amplifications by comparing the estimated $\omega_{D,CX}$ to the proportion of oncogenes amplified by each signature per cohort (average cosine similarities of 0.939; **Figure S25b**).

### Model performance assessment

#### Testing the predictive capacity of the forecasting framework

We assessed the predictive capacity of our forecasting framework by retrospectively evaluating models accuracy in the training cohort (TCGA, The Cancer Genome Atlas[94], comprising 7,880 samples from 33 tumour types) and in two independent cohorts (PCAWG, Pan-Cancer Whole Genome project[7], comprising 2,114 samples from 40 tumour types; and HMF; Hartwig Medical Foundation dataset[10], comprising 4,784 samples from 64 tumour types).

To assess the performance of each forecasting model on these large-scale datasets, we first selected samples from the specific tumour type for which the model parameters were trained. Additionally, when homozygous deletions in TSGs were forecasted, we further excluded samples without detectable CIN, given the recurrence of early TSG homozygous deletions that cannot be positively forecasted without additional CIN that makes the sample amenable to CIN signature quantification (**Figure S13**). Then, we labelled each sample based on the presence of the copy number alteration in the target driver gene (i.e. an homozygous deletion of the target tumour suppressor gene or a high-amplitude amplification of the target oncogene).

To estimate how accurately the model could predict these labels, we computed sample-level copy number signature activities ($l_{CX}$) for all the labelled samples. To prevent target leakage, prior to computing $l_{CX}$, we removed *in silico* the target oncogene amplification or TSG homozygous deletion and its adjacent segments. Using the $\omega_{D,CX}$ and $s_D$ terms trained in TCGA, we computed $P(D)$ for each sample and assessed how well this value aligned with the assigned labels by calculating the area under the receiver operating characteristic curve (AUC). We repeated this procedure for all tumour-specific and pan-cancer models. Only models for which at least 5 samples harboured an amplification were tested in each cohort. For each model, we used AUC as a measure of both predictive capacity and potential overfitting to the TCGA dataset (**Table S1**). We recommend to only apply out-of-the-box the models that demonstrate predictive capacity (AUC > 0.7) and have been validated in at least one independent cohort (PCAWG and/or HMF).

For models meeting this predictive criteria, we further evaluated three thresholding procedures, namely:

1. Younden's index: This procedure estimates the threshold as the *P(D)* value that maximises the Youden's index (sensitivity + specificity -1) in the training cohort.

2. Expected alteration frequency: It estimates the threshold as the *P(D)* value that equals the fraction of predicted high-risk samples in the training cohort to the expected rate of samples with the gene amplification or deletion. In practice, the threshold is set to the quantile $Q_{1-x}$ of the distribution of *P(D) values* in the reference cohort, where x represents the expert- or literature-derived expected frequency of driver alteration in the forecasting scenario. For the purpose of this high-throughput evaluation, we automatically set this expert-rate *x* to be the frequency of samples with the driver alteration in the training cohort. The threshold derived with this strategy places greater emphasis on specificity.

3. Minimum *P(D)* in carriers: It estimates the threshold as the lowest *P(D)* value across all cases with deletion or amplification of the gene of interest in the training cohort. The threshold derived with this strategy places greater emphasis on sensitivity.

All thresholding approaches require a training or reference cohort for derivation. To assess how the similarity between the reference and the target cohorts impact on threshold suitability, we evaluated two distinct scenarios. First, we simulated an ideal setting with a well-matched reference cohort by splitting the original cohort (either TCGA, PCAWG or HMF) into a training subset, representing 70% of the dataset, and a held-out testing subset, representing the remaining 30% of the dataset. To preserve the proportional representation of samples with and without the target copy number alteration across the training and testing datasets, we applied stratified subsampling. Specifically, 70% of the negative cases (without the copy number alteration) and 70% of the positive cases (with the copy number alteration) were randomly assigned to the training subset, while the remaining 30% were assigned to the testing subset. We set the threshold through any of the three procedures described above in the training subset and then evaluated the sensitivity, specificity and Youden's index resulting from applying these thresholds to the testing subset. We repeated this procedure 100 times and reported the average values across the 100 iterations (**Figure S4**). Second, we simulated a scenario with a less-matched reference cohort (i.e., differing sequencing technology and/or clinico-pathological status and/or biological characteristics) by training thresholds on TCGA and evaluating them on the independent PCAWG and HMF cohorts (**Figure S5**). Throughout this process, the threshold for *P(D)* binarization was optimized, while model parameters remained fixed as initially estimated in TCGA. Threshold performance was assessed by calculating sensitivity, specificity and Youden's index. Only models that showed evidence of potential predictive capacity were included in this evaluation (i.e. AUC > 0.7 in TCGA and HMF and/or PCAWG). To ensure robust evaluation, we only performed these assessments in cohorts with at least 10 cases exhibiting the target driver copy number alteration.

#### Assessing forecasting performance of the framework

We assessed the performance of our forecasting framework to predict acquisition of focal amplifications using longitudinal data. We performed the prospective assessment in 100 lung cancer patients from the TRACERx cohort[9]; and 44 prostate cancer patients from the [Pan-Prostate Cancer Group project](), the Hartwig Medical Foundation dataset[10], and from Mateo et al.[1]. **Table S2** includes a list of studies with patient-matched sample pairs that were excluded due to constraints such as limited cohort size, unavailable sequencing data or genome-wide copy number profiles, and lack of recurrent oncogene amplifications acquired.

##### Lung cancer longitudinal data

We downloaded copy number profiles from a total of 694 lung cancer samples (476 primary and 218 metastatic tumours) from 126 patients included in the TRACERx 421 cohort[9]. All samples have detectable CIN (≥ 20 CNAs).

In line with the results shown by Al Bakir et al.[9], the focal amplification of the *HIST1H3B* oncogene (copy number value ≥ 8) was exclusively observed in metastatic samples and may therefore confer increased metastatic potential. Consequently, we aimed at forecasting *HIST1H3B* amplification by applying our forecasting framework in primary lung cancer samples. Due to the metastatic-specific nature of *HIST1H3B* amplification, we were unable to train the model using the TCGA-LUSC and TCGA-LUAD cohort (only 2/541 TCGA lung samples exhibit this amplification). In addition, there is no evidence of positive selection for the amplification of this oncogene in any TCGA tumour-specific cohort[16]. For that reason, we learnt the different terms of our prediction model using metastatic tumours and computed the *HIST1H3B*-specific signature weights. Training was performed in an independent subset of metastases where no matched primary tumour sample was available for forecasting. Only one metastatic sample per patient was used in the learning step, prioritising samples collected from metachronous metastases in cases where a patient had multiple metastatic samples.

To forecast *HIST1H3B* amplification by applying our approach, we then took as input the copy number profiles of primary samples. We selected only those primary samples seeding metastases. Samples with a purity lower than 0.25 were excluded to avoid including samples with inaccurate copy number segmentation. We randomly selected one primary/metastasis pair per patient. For those patients that acquire *HIST1H3B* amplification in at least one metastatic sample, we prioritised the pair where the primary tumour seeded the metastasis harbouring the oncogene amplification. The acquisition of a *HIST1H3B* amplification in the metastatic biopsy was defined as the gain of at least 6 copies with respect to the primary sample. We computed amplification scores for a total of 100 primary samples, and classified samples as having high or low risk of acquiring *HIST1H3B* amplification in the metastasis. We established the classification threshold based on the frequency at which this amplification is observed in lung cancer (8-10%)[9,95]. To assess the performance of our forecasting framework, we computed the sensitivity and specificity to predict the oncogene amplification and evaluated the area under the ROC curve (AUC).

To demonstrate that the presence of amplifications and copy number alterations is not sufficient to forecast the acquisition of *HIST1H3B* amplification, we tested the predictive signal of four alternative genomic features: 1) having or not having CIN; 2) the number of copy number alterations; 3) having or not having amplifications; and 4) the number of amplifications. None of these three genomic features demonstrated the ability to predict *HIST1H3B* amplification, as reflected by achieving AUC values of 0.50, 0.56, 0.33 and 0.33, respectively (**Figure S6b**).

#### Prostate cancer longitudinal data

Here, we tested the ability of our forecasting framework to predict the acquisition of *AR* amplification as a resistance mechanism to androgen deprivation therapy (ADT) in prostate cancer. To do so, we collated and/or processed copy number profiles from the following 3 cohorts:

##### Hartwig Medical Foundation prostate tumour samples

We downloaded copy number profiles from a total of 5,200 samples released by the Hartwig Medical Foundation (HMF). Copy number profiles were derived from whole genome sequencing data using PURPLE as previously described[10,96].

As PURPLE[96] determines the allele specific copy number of every base of the genome, the genome binning resolution is significantly higher compared to the resolution of the copy number profiles used to derive feature components of our signature encoding (see section "Feature distributions and mixture modelling"). To avoid incorrect mapping of signatures due to differences in segmentation resolution, copy number profiles were binned into 30 kb bins and then resegmented. In our previous work[3], we estimated this bin size as appropriate to have a segmentation agreement between copy number calls derived from SNP6 arrays and WGS/WES. The copy number value of each 30 kb bin was computed by averaging the copy number value of the segments spanning a given bin. For bins spanning multiple copy number segments, this resegmentation may generate artificial segments with a 30 kb of length. Segments with a 30 kb length were therefore removed to avoid this artificial oversegmentation, and we then applied a smoothing procedure for merging continuous segments with a ± 0.1 difference in copy number.

To validate our forecasting framework, we focused on a subset of 308 longitudinally collected tumour pairs available within the HMF cohort, with a specific emphasis on 39 pairs where the primary tissue of origin was the prostate. All metastatic prostate cancer sample pairs were manually inspected and had good quality copy number profiles with sufficient segments supporting the relatedness between samples in the pair. We excluded 27 of these pairs because the initial sample already harboured an amplification in *AR* (copy number value > 3).

##### Mateo et al. 2020

We downloaded sWGS data from a total of 102 FFPE samples from 51 prostate patients[1]. Patient-matched primary tumour and metastatic castration resistant biopsies were available for all patients. Copy number profiles were derived as previously described (see section "Absolute copy number fitting from sWGS"). We excluded 35 pre/post-treatment sample pairs due to either low quality copy number profiles (i.e. oversegmented or low purity) or a lack of alignment with the profile of the corresponding pre/post-treatment sample.

##### Pan-Prostate Cancer Group cohort

We processed WGS data and derived copy number profiles from 16 time-separated pairs from patients included in the Pan-Prostate Cancer Group (PPCG). To have a comparable segmentation resolution with the reference cohort (TCGA), copy number profiles were derived after downsampling WGS data to SNP6 positions (as recommended by us in our previous work[3]). We obtained allele counts from alleleCounter (v4.1.0, default parameters with the "--dense-snps" option; https://github.com/cancerit/alleleCount) at SNP6 loci. For each sample, loci with at least 1 count in the tumour and 10 counts in the matched normal sample were used to derive logR and BAF using the default parameters of the *ascat.prepareHTS* function from the ASCAT repository (https://github.com/VanLoo-lab/ascat). LogR was then corrected for GC content and replication timing using the reference correction files for SNP6 positions provided in the ASCAT repository. Finally, the copy number calling was performed by running ASCAT[63] with a penalty of 70 and gamma equal to 1. For purity/ploidy selection, we fixed a range of ±0.2 with respect to the purity/ploidy estimates curated by the PPCG consortium by running Battenberg[97].

##### Integrated analysis

After processing and merging data from the 3 different cohorts, we applied our framework to forecast *AR* amplification in a total of 44 longitudinal pairs using as input the copy number profile derived from the early biopsy of each patient. We used the *AR*-specific, PRAD-specific signature weights learnt using the TCGA-PRAD cohort to compute amplification risk scores. Then, samples were classified as having high or low risk of acquiring *AR* amplification in the later biopsy by applying an optimal threshold determined in the TCGA-PRAD cohort. This threshold was established based on the known frequency of *AR* amplification acquisition after androgen deprivation therapy (45-52%)[98]. We specifically selected the score at which 45% of the TCGA-PRAD samples would be classified as having high amplification risk. Samples with no detectable CIN were classified in the low risk group.

We finally assessed the predictor performance by computing the sensitivity and specificity of forecasting *AR* amplification, and also by evaluating the area under the ROC curve (AUC). The acquisition of *AR* amplification was determined as the gain of more than 2 copies with respect to the early biopsy. Given that the amplification of the enhancer region is frequently observed to be amplified after ADT treatment[99], we scanned a genomic region consisting of 0.5Mb before the transcriptional start site and 1Mb after the coding region.

As in the lung cancer longitudinal data, we did not observe a better predictive signal for other alternative genomic features (presence of CIN: AUC=0.49; number of copy number alterations: AUC=0.58; presence of amplification: AUC=0.73; number of amplifications: AUC=0.78; **Figure S6a**).

### Demonstrating clinical utility of the predictor

#### Predicting prognosis

We applied our forecasting framework to compute the risk scores of acquiring *CDK4* amplification, *PDGFRA* amplification and *CDKN2A* deletion in 426 low-grade gliomas (LGG) from the TCGA project and 47 LGGs from the the Glioma Longitudinal Analysis Consortium (GLASS) dataset[8].

ASCAT-derived TCGA copy number profiles were downloaded from https://github.com/VanLoo-lab/ascat, and clinical metadata and genomic data for the TCGA cohort were downloaded from the TCGA website (https://portal.gdc.cancer.gov/) or via cBioPortal[100].

In the case of the GLASS cohort, we downloaded copy number profiles from a total of 504 diffuse glioma samples[8]. Copy number profiles were derived from WGS or WES data as previously described[8]. To avoid incorrect mapping of signatures due to differences in segmentation resolution, we applied the same resegmentation procedure used for processing the HMF profiles (see details in section “Hartwig Medical Foundation prostate tumour samples”). Once copy number profiles were processed, we excluded glioblastoma samples and samples with unknown histological subtype to exclusively include low-grade glioma patients in the downstream analyses. Given that the TCGA-LGG cohort consists of primary tumours, we only applied our model to early biopsies. Only one sample per patient was considered in the analysis. If a patient had multiple sample pairs, we prioritised the pair where the early biopsy was obtained from the primary tumour. If this was not feasible, we randomly selected one of the available sample pairs.

We first applied a univariate (Kaplan-Meier estimates) and a multivariate method (Cox proportional hazard model) for comparing overall survival between LGG patients classified based on the current WHO Classification of Tumors of the Central Nervous System[101–103] (**Figure 3a, Figure S8a**).

Next, given the link of *CDK4* and *PDGFRA* amplification status to poor prognosis[104–106], we further classified LGG patients with intermediate prognosis based on their amplification status in these two oncogenes (**Figure S7a**). Survival analyses were then performed for comparing overall survival across groups as previously described (**Figure S7b**).

To demonstrate the potential of implementing our forecasting framework into the brain tumour diagnosis algorithm, we further classified IDHmut-non-codeleted LGG patients without *CDK4/PDGFRA* amplifications and *CDKN2A* deletions by applying our forecasting framework (**Figure 3b**). Survival analyses were then performed for comparing overall survival across groups as previously described (**Figure 3b, Figure S8b**). The implementation details for each model used in sample classification are outlined below:

- Forecasting *CDK4* amplifications: We used the *CDK4*-specific, LGG-specific signature weights learnt from the TCGA-LGG cohort for computing amplification scores. Then, the frequency at which the amplification is found in the TCGA-LGG cohort (~10%) was used to define the threshold for classifying those patients without *CDK4* amplification but with different risks of acquiring it in the future. To classify GLASS patients as having high or low risk of acquiring *CDK4* amplification, we applied the frequency-based threshold learnt using the TCGA-LGG cohort (reference cohort).

- Forecasting *PDGFRA* amplifications: We used the *PDGFRA*-specific, LGG-specific signature weights learnt from the TCGA-LGG cohort for computing amplification scores. Then, the frequency at which the amplification is found in the TCGA-LGG cohort (~5%) was used to define the threshold for classifying those patients without *PDGFRA* amplification but with different risks of acquiring it in the future. To classify GLASS patients as having high or low risk of acquiring *PDGFRA* amplification, we applied the frequency-based threshold learnt using the TCGA-LGG cohort (reference cohort).

- Forecasting *CDKN2A* homozygous deletions: We used the *CDKN2A*-specific, LGG-specific signature weights learnt from the TCGA-LGG cohort for computing deletion scores. Deletion scores were computed for all LGG tumour samples, and scores were subsequently divided by the copy number value of the TSG in the sample. This allowed to correct the risk by the probability of hits needed to acquire an homozygous deletion. Then, the frequency at which the homozygous deletion is found in the TCGA-LGG cohort (~15%) was used to define the threshold for classifying those patients without *CDKN2A* homozygous deletion but with different risks of acquiring it in the future. To classify GLASS patients as having high or low risk of acquiring *CDKN2A* deletions, we applied the frequency-based threshold learnt using the TCGA-LGG cohort (reference cohort). Samples with no detectable CIN (< 20 CNAs) were included in the low risk group when deletion scores were lower than the threshold, otherwise they were classified as high risk of acquiring *CDKN2A* deletion.

Kaplan-Meier estimates (function *survfit*) and Cox proportional hazard models (function *coxph*) were performed using the survival and survminer R packages. Cox proportional hazard models were corrected by the fraction of genome altered and age at diagnosis. Age at diagnosis was categorised into two groups (< 40 versus ≥40 years) based on clinical risk[107]. Those LGG samples with a deletion of *CDKN2B* were excluded in the analysis due to its significant correlation with worse clinical behaviour and shorter patient survival[108]. We verified the fit of the proportional hazards assumption using the function *cox.zph*.

Previous research has highlighted the pivotal role of ecDNA in gliomas progression, with the majority of oncogene amplifications in gliomas occurring extrachromosomally[109]. To validate our forecasting framework, we demonstrated that the sample-level activity of CX13, a signature linked to ecDNA that showed a significant enrichment in *CDK4* and *PDGFRA* in both low-grade gliomas and glioblastomas (**Figure S31**), was not sufficient for predicting overall survival in the TCGA-LGG cohort (**Figure S26**). In addition, we also demonstrated that the sample-level activity of CX2, a signature linked to extensive deletions across the genome that showed a significant enrichment in *CDKN2A* in both low-grade gliomas and glioblastomas (**Figure S33**), was not sufficient for predicting overall survival in the TCGA-LGG cohort (**Figure S27**).

The amplification and/or activation of *MYC* at diagnosis has been previously associated with poorer post-progression survival, likely due to inducing temozolomide resistance via promoting hypermutation[110]. We applied our framework to forecast *MYC* amplification in tumour samples from the TCGA-LGG cohort. To this purpose, we used the *MYC*-specific, LGG-specific signature weights learnt from the TCGA-LGG cohort for computing amplification scores. The frequency of the amplification in the sample subset (~3%) was used as the threshold for classifying patients without *MYC* amplification as having a high or low risk of acquiring it in the future. For this analysis, apart from excluding samples with *CDKN2A/B* deletions, we also removed samples with *CDK4/PDGFRA* amplification. Notably, only two of the samples were classified as having high risk of acquiring both *MYC* and *CDK4/PDGFRA* amplification, suggesting that distinct biological processes may underlie the amplification of each oncogene. No significant differences in overall survival were observed between samples with low and high *MYC* amplification risk (HR = 1.1, p-value = 0.87; independent of fraction of genome altered, molecular subtype, and age at diagnosis; **Figure S28**). Patients with *MYC* amplification indeed showed similar survival intervals to those without the amplification (HR = 2.4, p-value = 0.25; independent of fraction of genome altered, molecular subtype, and age at diagnosis; **Figure S28**). Therefore, forecasting *MYC* amplification appears to lack prognostic potential in this cohort.

### Predicting amplification-driven drug resistance

#### Lung cancer cell lines

We computed *MET* amplification scores in four parental *EGFR*-mutant NSCLC cell lines, including HCC827, HCC4006, PC9 and H1975, and observed the acquisition of *MET* amplification in the resistant clones emerged after long-term culture in the presence of EGFR-TKI (see section “Experimental data: Human cancer cell lines” for further details). To apply our forecasting framework, we used the parental copy number profiles as input files and applied the *MET*-specific signature weights learnt from the TCGA cohort. Droplet digital PCR (ddPCR) was used to detect *MET* amplification in both the parental and resistant cell lines, using the ddPCR™ Copy Number Assay: MET, Human (dHsaCP1000038). We used the ddPCR™ Copy Number Assay: AGO1, Human (dHsaCP2500349) as a reference. Copy number was estimated as the ratio of the concentration of *MET* with respect to *AGO1* using the Bio‑Rad QuantaSoft software. *MET* amplification was absent in the parental cultures. An acquisition of *MET* amplification-driven resistance was considered when resistant clones showed 2-fold increase of the parental value (**Figure S9**).

To validate results observed in our experimental design, we also downloaded sequencing data from an independent study in which 3 *EGFR*-mutant NSCLC cell lines (HCC827, HCC4006 and PC9) were long-term cultured in the presence of erlotinib until the emergence of resistant clones[2]. Copy number profiles were derived from WGS and/or WES data as previously described (see section “Generating copy number profiles”), and then *MET* amplification scores were computed in the parental cell lines. We observed the same result as that obtained from our experimental design, with the highest amplification score assigned to the HCC827 parental cell line. This was the unique cell line reported to acquire *MET* amplification as the resistant mechanism to erlotinib treatment (**Figure 4a, Figure S10**).

To confirm *MET* amplification was responsible for EGFR-TKI resistance in HCC827 cells, we sorted 64 single cells treated with EGFR-TKI and 64 single cells treated with a combination of EGFR-TKI and METi (see section “Human lung cancer cell lines”), from which 93 cells were sequenced (see section “Single-cell whole genome sequencing”). After quality control, we derived copy number profiles for up to 35 cells per treatment (see section “Generating copy number profiles”). The absolute number of copies of *MET* was then determined from each single-cell copy number profile, and the cells with at least 8 copies were classified as having a focal *MET* amplification. As a control, we also sorted 96 single cells from the parental HCC827 cell line and successfully derived copy number profiles for 59 of them. None of these parental HCC827 cells exhibited *MET* amplification (**Figure S11**).

#### Lung cancer patients

We applied our forecasting framework to compute the *MET* amplification scores in 33 primary *EGFR*-mutant non-small cell lung cancers (NSCLC) treated with osimertinib (**Figure S12**). These patients were part of the OSIRESP cohort (see section “Experimental data: Patient sample cohort”). In this case, we also used the *MET*-specific signature weights learnt from the TCGA cohort for computing the predictions. Given the lack of a training cohort with similar clinical and biological context, we used *EGFR*-mutant NSCLC cell lines for estimating the appropriate threshold for classifying patients as having high or low risk of *MET* amplification. We selected a score of amplification risk that accurately separates the cell lines that acquire and do not acquire *MET* amplification as the resistance mechanism (0.06).

For comparison purposes, we grouped patients classified as having “complete response” and “partial response” according to RECIST criteria in one group, and patients classified as “progressive disease” and “stable disease” according to RECIST criteria in another group. Six out of the 13 non-responders patients were classified as having high risk of acquiring *MET* amplification, while only 2 out of 20 responders patients were included in the high risk group.

In order to further test the clinical utility of our forecasting framework, we performed a survival analysis to compare both the overall survival (OS) and progression-free survival (PFS) between patients predicted as having high or low risk of acquiring *MET* amplification (and therefore as being likely to show resistance to osimertinib via *MET* amplification or show response or resistance via other mechanism, respectively). Kaplan-Meier curves were used to illustrate the differences in OS and PFS between the two risk groups. Cox proportional hazards regression models were applied to predict hazard ratios. Cox proportional hazard models were corrected by the fraction of genome altered, the presence of metastases in the central nervous system (CNS) at treatment initiation, and the physical activity based on the Eastern Cooperative Oncology Group (ECOG) scale at treatment initiation. We verified the fit of the proportional hazards assumption using the function *cox.zph* from the survival R package.

The amplification of *ERBB2* has been previously linked to resistance to the anti-EGFR antibody cetuximab in EGFR-mutant lung cancer patients[111]. We applied our framework to forecast *ERBB2* amplification in samples from the OSIRESP cohort. To this purpose, we used the *ERBB2*-specific signature weights learnt from the TCGA cohort for computing amplification scores. Given the lack of both a training cohort and an *in vitro* study, the frequency at which *ERBB2* amplification-driven resistance is expected after EGFR inhibition (12%)[111] was used as a threshold for classifying patients as having a high or low risk of acquiring it in the future. No significant differences were observed between samples with low and high *ERBB2* amplification risk in overall survival (HR = 0.88, p-value = 0.873; independent of the fraction of genome altered, the presence of CNS metastases at treatment initiation, and the ECOG status at treatment initiation; **Figure S29**) and in progression-free survival (HR = 1.29, p-value = 0.710; independent of the fraction of genome altered, the presence of CNS metastases at treatment initiation, and the ECOG status at treatment initiation; **Figure S29**). Therefore, forecasting *ERBB2* amplification appears to lack potential to predict osimertinib resistance.

### Exploring mutational processes underlying driver alterations

#### Exploring processes underlying oncogene amplifications

Computing the enrichment of CIN signatures in oncogene amplifications (signature weights) allowed us to link the observed oncogene amplifications directly back to putative causal CIN signatures.

We applied the procedure for computing the oncogene-specific signature weights to 6,335 genome-wide copy number profiles from 33 cancer types derived from SNP6 array data from the TCGA. From a total of 271 oncogenes included in our list, we limited this exploration analysis to those oncogenes that are putatively selected to be amplified in a given tumour type (**Figure S1**). In total, we computed signature weight vectors ($\omega_{OG,t,CX}$) for 352 oncogene/tumour type pairs. We first explored the contribution of the 9 CIN signatures to oncogene amplification at a pan-cancer level. Unlike in the context of a specific tumour, we must capture features impacting on the acquisition and expansion of oncogene amplifications across tumour types to correctly estimate the pan-cancer CIN signature weights for each oncogene. Hence, we first multiplied the oncogene-specific CIN signature weights by the amplification frequency observed in the TCGA tumour-specific cohort. Then, the corrected signature weights per oncogene were averaged across all tumour types in which the given oncogene amplification confers a positive fitness effect. Performing this weighted average allowed to assign greater contribution to the tumour types where the amplification is more frequently observed. The *Heatmap* function from *ComplexHeatmap*[57] was applied for identifying groups of oncogenes with amplifications putatively caused by similar mutational processes. The number of clusters was set to 8 (**Supplementary Note Figure 3a**).

We next explored the contribution of CIN signatures to oncogene amplifications across 21 tumour types and 95 different oncogenes (**Figures S30-S31**). Hierarchical clustering (using the *hclust* function from the *stats* package in R[74]) was used for grouping tumour types without forcing the number of clusters. Interestingly, we observed both shared and variable patterns of signature enrichment across tumour types for given oncogenes. We observed some oncogenes with similar CIN signatures weights pan-cancer (i.e. *CDK4* or *CCND1*), while others showed more diverse patterns across tumour types (i.e. *MYC* or *PDGFRA*), likely due to cell-of-origin differences in replicative, transcriptional, repair and chromatin dynamics[112].

#### Exploring processes underlying tumour suppressor gene deletions

We applied the same enrichment strategy to identify the putative CIN signatures associated with homozygous deletions of 12 tumour suppressor genes that are under positive selection across various tumour types, resulting in a total of 52 tumour suppressor gene/tumour type pairs (**Figure S2**). As for oncogene amplifications, this exploration was performed at pan-cancer and tumour type-specific level. The *Heatmap* function from *ComplexHeatmap*[57] was applied for clustering tumour suppressor genes with homozygous deletions caused by similar mutational processes (**Supplementary Note Figure 3b, Figure S32**). We also grouped tumour types with similar CIN signatures weights in a specific tumour suppressor gene (**Figure S33**).

#### Supporting putative processes underlying oncogene amplifications

We tested for a correlation between the frequency at which an specific oncogene is present in the different amplicon categories and the signatures enriched in the amplified segments spanning the given oncogene. In detail, we downloaded the complete list of amplicons found in TCGA and PCAWG from Kim et al. (2020)[6], which were classified into four categories as circular, breakage-fusion-bridge (BFB), heavily-rearranged, and linear. We then computed the frequency of each amplicon category harbouring a specific oncogene. This frequency was assumed to be a readout of the preferred mechanism to amplify a specific oncogene. For each category, we then used a Pearson correlation to assess the association between the amplicon type frequency and the signature enrichment per oncogene. This analysis was performed for all 271 oncogenes included in our list.

### Exploring acquisition and selection of oncogene amplifications

#### Amplification acquisition

We explored the acquisition of amplifications during recent karyotypic diversification using publicly available single-cell sequencing data from 12 human triple negative breast cancer (TNBC) samples obtained in two prior studies[11].

To do so, we first classified focal amplifications detected in single cells as shared or unique. We defined shared amplifications as genomic regions with absolute copy number higher or equal to 8 at bulk resolution. Potential unique amplifications were then detected as gains of at least 6 copies with respect to the pseudobulk copy number profile. To truly determine that these potential amplifications were present in only a single cell, we took a heuristic approach, motivated by the uncertainty in the location of the breakpoints during segmentation of single cell copy number profiles. In brief, we extended each of the putative unique amplifications by a 1 Mb window per segment side, and looked for overlaps with the amplifications present in other cells. If we found overlaps, the amplification was classified as shared, and otherwise as unique.

Focal amplifications present in one single cell (unique amplifications) must have recently been generated, while amplifications shared across cells (i.e. detectable at bulk resolution) must have been acquired previously in the evolution of a tumour. We compared the number of unique focal amplifications detected across all single cells with the number of clonally-expanded focal amplifications seen at pseudobulk resolution (**Supplementary Note Figure 2b**).

### Positive selection test

We tested the role of selection in sculpting the landscape of focal amplifications in a tumour. Briefly, we evaluated enrichment in the number of i) unique amplifications in these 12 TNBC samples and ii) shared amplifications in 145 TCGA-TNBC samples mapping to oncogenes against a background distribution derived by using a Monte Carlo approach (**Supplementary Note Figure 2c**).

First, we empirically selected the 10 oncogenes most frequently amplified in the TCGA-TNBC cohort and counted the number of focal amplifications mapping to those 10 tumour-specific oncogenes. The background distribution was then derived by counting the number of focal amplifications mapping 10 non-cancer genes which were randomly selected from a list of 53,989 total non-cancer genes annotated in Ensembl GRCh37 genome assembly. In total, we ran this procedure 1,000 times.

Statistical significance was measured by calculating the empirical p-value. Empirical p-values were computed as the fraction of iterations where the number of amplifications within the 10 non-cancer genes (background) was greater than the number of amplifications observed within the 10 TNBC-specific most recurrently amplified oncogenes.

### Distribution of unique amplifications across the genome

We tested whether the acquisition of amplifications was uniform across the genome. To this end, we explored the genomic distribution of unique amplifications in single cells from 12 TNBC samples[11,113].

For each sample, we initially mapped all observed unique amplifications to their respective genomic regions, arranging them in sequential order from the start to the end of the chromosome. We next calculated the distance of each unique amplification to its nearest adjacent unique amplification. These distances were then normalised by the total length of the chromosome. Only unique amplifications located in autosomal chromosomes were considered in this analysis.

Next, for each autosomal chromosome, we estimated the expected distance between adjacent unique amplifications, assuming a homogeneous amplification rate across the genome. This distance was computed by dividing the chromosome length by the total number of unique amplifications observed on it. We normalised this distance by the chromosome length to facilitate the comparison across chromosomes, resulting in the expected distance being equal to 1/(number unique amplifications + 1).

Finally, we transformed those distances to a relative measure by subtracting the expected distances to the observed distances. This relative distance facilitates interpretation: a distribution centred at 0 implies an even distribution of unique amplifications across the genome. We determined if our distance distribution significantly deviated from this expectation by performing a one-sample Wilcoxon two-sided test, with the null hypothesis assuming a mean distance equal to 0 and the alternative hypothesis suggesting a mean distance different from 0 (**Supplementary Note Figure 2d**).

#### Birth-death simulations

We performed simulations to model the acquisition of an oncogenic amplification with varied amplification rates and selection coefficients. We model cancer evolution as a stochastic system of birth (occurring at rate $\beta$) and death (occurring at rate $\alpha$) events. In this process, cells without oncogene amplification can acquire it in the process of cell division at rate $\mu$ (**Supplementary Note Figure 2e**). This means that the acquisition of oncogene amplification occurs at overall rate $\beta\mu$. In our simulations, we conferred to cells with oncogene amplification a fitness advantage ($s$), which increases the division rate by ($1+s$) and decreases death rate by ($1-s$). Overall, this means that tumour cells without oncogene amplification die with rate $\alpha$, divide yielding two cells without oncogene amplification with rate $\beta(1-\mu)$ and divide giving rise to one cell with oncogene amplification and another without with rate $\beta\mu$. Meanwhile, tumour cells with oncogene amplification die with rate $\alpha(1-s)$ and divide with rate $\beta(1+s)$, giving rise to two cells with oncogene amplification.

We used the Gillespie algorithm[114] to implement the simulations in practice. For all simulations, we used the following fixed parameters: starting population of 1,000 cells, cell division rate of 0.3 and cell death rate of 0.15. In the different simulations, we tested a range of oncogene amplification rates (from 5e-6 to 0.1) and selection coefficients (from 0 (neutral selection) to 1). Simulations of tumour growth were run until the population size reached 200,000 cells or until 20,000 generations.

In total, we ran 20 different simulations for each of the combinations of amplification rate and selection coefficient. For each simulation, we calculated the fraction of cells that acquired the oncogene amplification and computed the average across simulations with the same parameters (**Supplementary Note Figure 2f**).
